# Antibody optimization enabled by artificial intelligence predictions of binding affinity and naturalness

**DOI:** 10.1101/2022.08.16.504181

**Authors:** Sharrol Bachas, Goran Rakocevic, David Spencer, Anand V. Sastry, Robel Haile, John M. Sutton, George Kasun, Andrew Stachyra, Jahir M. Gutierrez, Edriss Yassine, Borka Medjo, Vincent Blay, Christa Kohnert, Jennifer T. Stanton, Alexander Brown, Nebojsa Tijanic, Cailen McCloskey, Rebecca Viazzo, Rebecca Consbruck, Hayley Carter, Simon Levine, Shaheed Abdulhaqq, Jacob Shaul, Abigail B. Ventura, Randal S. Olson, Engin Yapici, Joshua Meier, Sean McClain, Matthew Weinstock, Gregory Hannum, Ariel Schwartz, Miles Gander, Roberto Spreafico

**Affiliations:** Absci Corporation, Vancouver (WA), USA

## Abstract

Traditional antibody optimization approaches involve screening a small subset of the available sequence space, often resulting in drug candidates with suboptimal binding affinity, developability or immunogenicity. Based on two distinct antibodies, we demonstrate that deep contextual language models trained on high-throughput affinity data can quantitatively predict binding of unseen antibody sequence variants. These variants span a *K*_*D*_ range of three orders of magnitude over a large mutational space. Our models reveal strong epistatic effects, which highlight the need for intelligent screening approaches. In addition, we introduce the modeling of “naturalness”, a metric that scores antibody variants for similarity to natural immunoglobulins. We show that naturalness is associated with measures of drug developability and immunogenicity, and that it can be optimized alongside binding affinity using a genetic algorithm. This approach promises to accelerate and improve antibody engineering, and may increase the success rate in developing novel antibody and related drug candidates.

## Introduction

Despite billions of dollars of investment every year, only an estimated 4 % of drug leads succeed in their journey from discovery to launch [1]. Even worse, only 18 % of drug leads that pass preclinical trials eventually pass phase I and II trials, suggesting that most drug candidates are unsafe or ineffective [2]. While much of this failure rate is attributable to incomplete understanding of the underlying biology and pathology, insufficient drug lead optimization contributes to a large number of failures [3].

Traditional antibody screening approaches can only explore small regions of the sequence space. This may constrain results to sequences with suboptimal properties such as insufficient binding affinity, developability limitations, and poor immunogenicity profiles [4]. By contrast, deep mutagenesis coupled with screening or selection allows for the exploration of a larger antibody sequence space, potentially yielding more and better drug leads [5]. However, deep mutagenesis comes with its own challenges. For example, most mutations degrade the binding affinity of antibodies rather than improve it, which greatly reduces screening efficiency. Moreover, the size of the antibody sequence variant space grows exponentially with mutational load (i.e. the number of mutations simultaneously introduced into each sequence variant) and quickly exceeds the capacity of experimental assays by orders of magnitude. In addition, most antibody screening approaches are limited to screening only one property at a time, restricting the simultaneous optimization of drug potency and developability. Because improving a property may negatively impact others, simultaneous, rather than sequential, optimization of antibody properties is a preferable therapeutic strategy [6].

Deep neural networks are an emerging tool that can help overcome the limitations of experimental screening capacity [7]. The general approach involves training a model on experimental data and applying it to predict which sequences are most likely to improve the measured trait. Several promising approaches have been proposed [8–14], but only two studies have had *in silico* predictions validated in the lab [15, 16]. While being valuable demonstrations, previous models are limited by throughput and the use of binary (rather than continuous) readouts, which can compromise their accuracy at high mutational loads.

In this study, we demonstrate our capability to improve the binding affinity of an antibody for its target antigen using deep contextual language models and quantitative, high-throughput experimental binding affinity data. We show that models can quantitatively predict binding affinities of unseen antibody variants with high accuracy, enabling virtual screenings and augmenting the accessible sequence space by orders of magnitude. In this sense, the trained learner can serve as an oracle, assigning functional annotations from just sequence [17, 18]. We confirm predictions and consequent designs in the lab, with a much higher success rate than would be attained with traditional screening.

An additional concern for antibody screening approaches is that the improvement of binding affinity can negatively affect developability and immunogenicity properties [19]. This issue would remain unaddressed by machine learning models trained to optimize affinity without regard for other properties. Here we introduce natural antibody sequences into our language models, allowing us to characterize the *naturalness* of any given sequence for a host species. We find that high naturalness scores are associated with improved immunogenicity and developability metrics, thereby highlighting the importance of simultaneously optimizing multiple antibody properties during drug lead screening. To address this task, we present a genetic algorithm for the efficient identification of sequences with both strong binding affinity and high naturalness.

## Results

### Deep language models can predict binding affinity of sequence variants

We hypothesized that artificial intelligence (AI) models based on deep neural networks could learn the mapping between variants of a biological sequence (such as an antibody) and quantitative readouts (such as binding affinity) from experimental data. With this capability, AI models could be used to simulate experiments *in silico* for novel sequences, thereby accessing more variants with improved properties at a lower cost (fig. 1).

**Figure 1.**
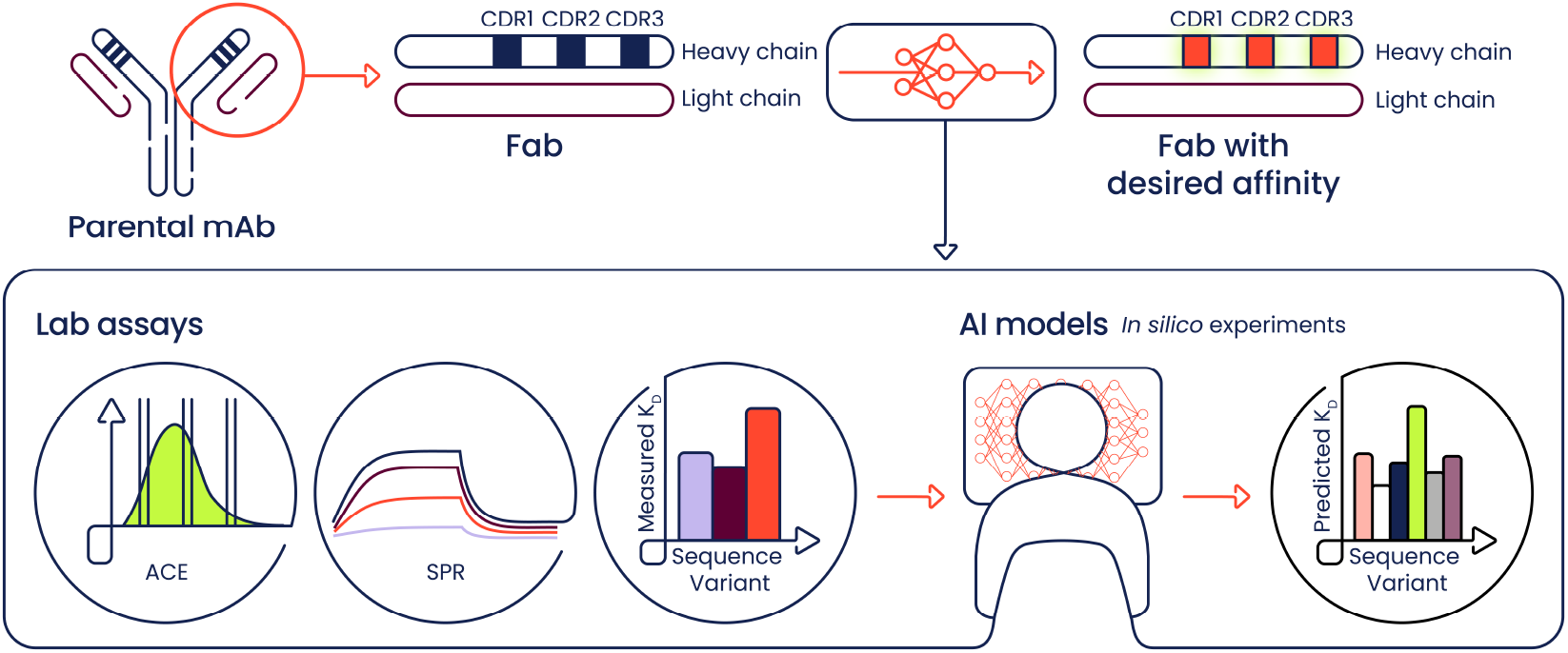
AI-augmented antibody optimization. Deep learning models fed with ACE or SPR measurements can quantitatively predict affinities of novel sequence variants, thereby enabling the *in silico* design of antibodies with desired binding properties.

Training of deep learning models requires large, high-quality datasets. To generate high-throughput measurements of antibody binding affinities, we developed the Activity-specific Cell-Enrichment (ACE) assay (fig. S1), a method based on Fluorescence-Activated Cell Sorting (FACS) and Next-Generation Sequencing (NGS). The assay is an improved version of our prior work [20]. The ACE assay leverages intracellular, soluble overexpression of folded antibodies in the SoluPro™ *E. coli* B Strain. Cells expressing antibody variants are fixed, permeabilized and stained with fluorescently-labeled antigen and scaffold-targeting probes. Cells are then binned and sorted based on binding affinity and expression level of variants. Finally, the collected DNA sequences are amplified via PCR and sequenced. ACE scores are calculated from sequencing read counts (See Methods) and are proportional to binding affinities.

In order to assess whether the sequence-affinity relationship can be modeled and predicted, we generated variants of the HER2-binding antibody trastuzumab in Fragment antigen-binding (Fab) format. Mutagenesis of CDRH2 and CDRH3 was prioritized as these regions accommodate the highest density of paratope residues, both in general and for trastuzumab [21, 22]. Across this study, up to five simultaneous amino acid substitutions were introduced randomly in the parent antibody, in up to two CDRs, allowing all natural amino acids except cysteine (excluded to avoid potential disulfide bond-related liabilities). Table 1 summarizes the datasets used to train models.

**Table 1.** Trastuzumab variant datasets. Characteristics of datasets used to train and evaluate models. The positions hosting substitutions (IMGT numbering), number of simultaneous substitutions (mutational load), and allowed amino acids (all except cysteine) define the combinatorial sequence space. A subset of sequences was sampled from the combinatorial sequence space according to the indicated design strategy to build libraries for screening by the ACE assay or SPR. The numbers of QC-passing amino acid sequence variants upon screening and analysis are shown, broken down by mutational load. ^*^ Random sampling of combinatorial space. ^**^ Uniform sampling by affinity from the trast-1 dataset. ^***^ Random sampling of combinatorial space per mutational load bin, with defined prevalence ratios of mutational load bins. Quadruple and quintuple mutants were used only to assess the performance of predictions from models trained with up to triple mutants.

| Dataset |  | trast-1 | trast-2 | trast-3 |
| --- | --- | --- | --- | --- |
| Screening |  | ACE | SPR | ACE |
| Mutated CDRH2 positions |  | - | - | 10 (55-66) |
| Mutated CDRH3 positions |  | 8 (107-114) | 8 (107-114) | 10 (107-116) |
| Mutational load |  | Up to double mutations | Up to double mutations | Up to triple mutations |
| Allowed natural AAs |  | 19 (no Cys) | 19 (no Cys) | 19 (no Cys) |
| Combinatorial space |  | 9,217 | 9,217 | 6,710,401 |
| Design |  | Random* | Uniform** | Random, stratified*** |
| # Measured AA variants |  | 8,932 | 215 | 52,596 |
| Number of mutations in AA variants | 0 | 1 | 1 | 1 |
|  | 1 | 142 | 23 | 315 |
|  | 2 | 8,789 | 191 | 4,054 |
|  | 3 | - | - | 44,704 |
|  | 4 | - | - | 1,992 |
|  | 5 | - | - | 1,530 |

In addition to high-throughput (HT) ACE data, we also leveraged low-throughput, but highly accurate SPR *K*_*D*_ readouts to assess binding affinity. SPR was used for *(i)* targeted re-screening of sequence variants upon primary screening with the ACE assay; and *(ii)* to validate model predictions.

As a proof of concept for our workflow, we created a library containing all sequence variants with up to two mutations across eight positions of trastuzumab CDRH3 (fig. 2A). Using the ACE assay, we measured the binding affinity of 8,932 variants (97 % of the combinatorial space) to create the trast-1 dataset (table 1). We trained a deep language model using 90 % of the trast-1 dataset and evaluated the model predictions using the remaining 10 % as hold-out data. The measured and predicted ACE scores for the hold-out dataset were highly correlated, indicating that the language model could predict binding affinity with high accuracy (fig. 2B).

**Figure 2.**
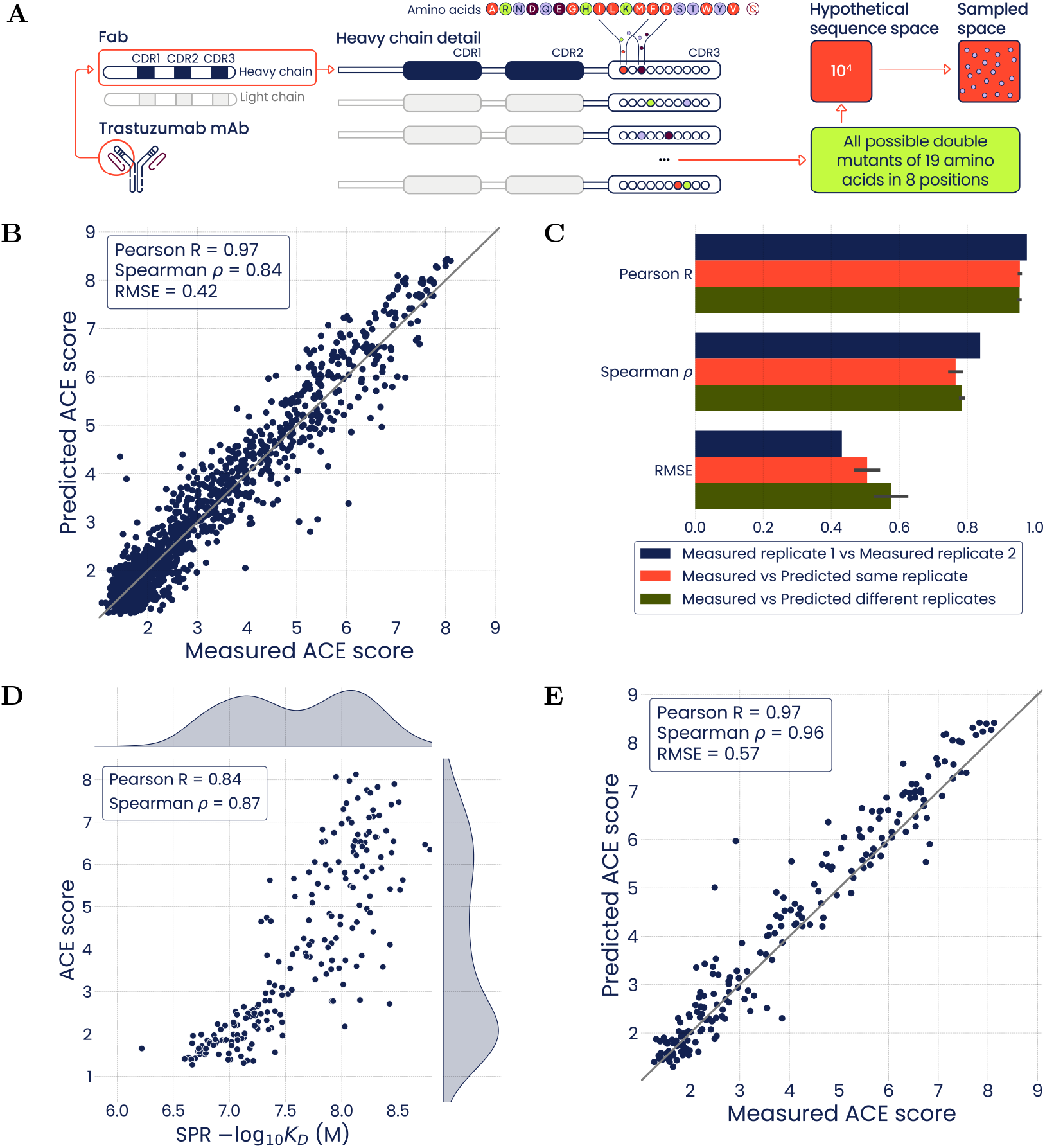
Deep language models trained with the ACE assay generated trast-1 dataset quantitatively predict antibody binding affinity. **(A)** Illustration of the combinatorial mutagenesis strategy of the trast-1 dataset: up to double mutants in 8 positions of the CDRH3 of trastuzumab, screened using the ACE assay. **(B)** Predictive performance of a model trained on ACE assay scores of variants from 90 % of trast-1, evaluated on the remaining 10 % of sequences. **(C)** Comparative analysis of replicate ACE assay measurements and ACE assay scores predicted from models trained on individual ACE assay replicates. Error bars are 95 % confidence intervals. **(D)** Correlation between ACE assay affinity score and log-transformed SPR *K*_*D*_ measurements. Plot shows ACE assay scores from trast-1 for sequence variants intersecting with trast-2. **(E)** Predictive performance against a hold-out set uniformly distributed with respect to binding affinity (ACE scores from trast-1 for sequences shown in panel **D**).

Inaccuracy in predictions is affected by both modeling errors and experimental noise. To disentangle these two effects, we looked at the agreement between measurement replicates using the same metrics we previously used to assess the predictive performance of our models (fig. S2A). Evaluating the model performance relative to the agreement of measurement replicates indicated that most of the prediction error could be attributed to experimental noise (fig. 2C, fig. S2B).

The hold-out set evaluated in fig. 2B was randomly drawn from the trast-1 dataset. Therefore, training and hold-out sets had similar distributions of ACE scores, with a prevalence of low-affinity binders due to the detrimental effect of most mutations. This design of training and hold-out sets addressed the question of whether models can simulate experiments *in silico*. A more challenging test would involve assessing predictions using a hold-out set distributed uniformly with respect to binding affinities. This hold-out set would be enriched in strong binders relative to the training set. To reduce the prevalence of weak binders in this new hold-out set, we sampled >200 sequences from the trast-1 dataset. The sampled sequences were rescreened by SPR to create the trast-2 dataset (table 1). As expected, we observed strong agreement between ACE scores and SPR-derived- log_10_ *K*_*D*_ values of trast-2 sequences (fig. 2D), and confirmed the near-uniform distribution of binding affinities for this dataset. We then used the trast-2 sequences as a hold-out set for models trained with trast-1 ACE scores, which confirmed strong predictive performances (fig. 2E).

Since we collected SPR measurements for the trast-2 dataset (fig. 2D), we investigated whether this dataset alone was sufficient to train a deep language model to directly predict equilibrium dissociation constants. Due to the relatively small size of the dataset (n=215), all models were trained using 10-fold cross-validation and model performance was evaluated using pooled out-of-fold predictions. We first trained a model to predict - log_10_ *K*_*D*_ values, and found that the correlation between measured and predicted values was slightly lower than that observed with the high-throughput trast-1 dataset (fig. 3A). However, 87 % of predicted binding affinities deviated by less than half of a log from their respective measured values. As in the case of trast-1, we also evaluated the trast-2 results relative to the best possible performance defined as the degree of agreement between measurement replicates (fig. 3B, fig. S3A-B).

**Figure 3.**
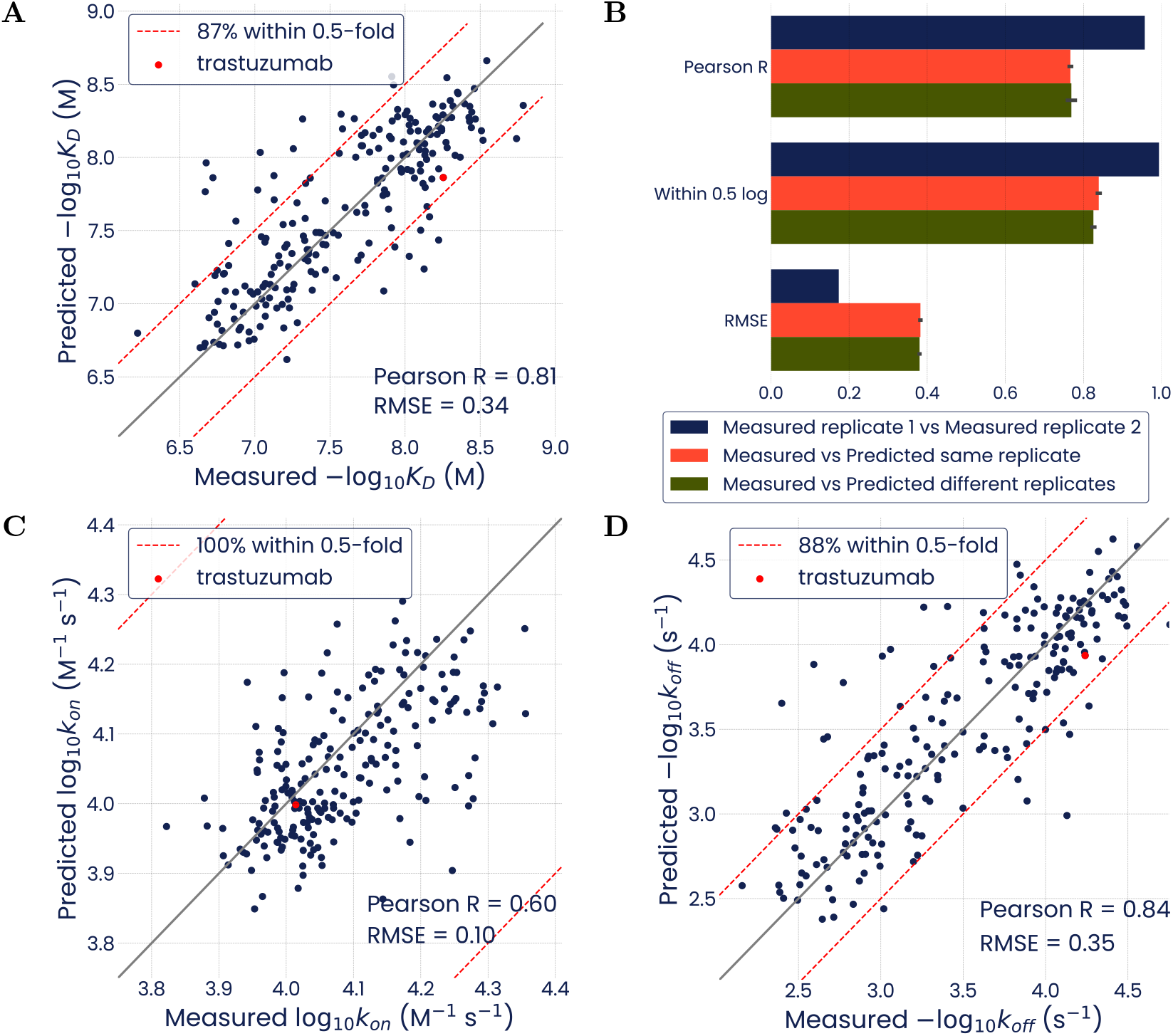
Deep language models trained with the SPR-generated trast-2 dataset quantitatively predict antibody binding affinity. Performance is evaluated by pooled 10-fold cross-validation. **(A)** Predictions from a model trained on SPR-measured - log_10_ *K*_*D*_ values. **(B)** Comparative analysis of replicate - log^10^ *K*_*D*_ measurements and- log^10^ *K*_*D*_ predicted from models trained on individual SPR replicates. Error bars are 95 % confidence intervals. **(C)** Predictions from a model trained on log^10^ *k*_*on*_ values. **(D)** Predictions from a model trained on - log^10^ *k*_*off*_ values.

In addition to equilibrium binding constants, SPR provides association (*k*_*on*_) and dissociation (*k*_*off*_) coefficients. Models trained to predict these coefficients also performed well (fig. 3C-D, figs. S4 and S5), opening the possibility for AI to aid the specific engineering of association and dissociation properties, in addition to the overall binding affinity. Note that the lower correlation coefficient observed for *k*_*on*_ was due to the small range of observed variation. Similarly, the agreement of measurement replicates was also lower for *k*_*on*_ than for *k*_*off*_, which further underscores the need to consider measurement noise when assessing prediction performances.

Finally, we asked whether a model simultaneously trained with two affinity data types could improve the performance compared to a model fed with just a single data type. For this, we supplemented the trast-2 model with trast-1 ACE assay data, using a multi-task training setting. We found that this model slightly out-performed the original model trained only on trast-2 SPR data (fig. S6).

All models trained on the trast-1 and trast-2 datasets were deep language models pre-trained on immunoglobulin sequences from the OAS database (see Methods). We compared these models against baselines, either using a 90:10 train:hold-out split from the trast-1 dataset or a pooled 10-fold cross-validation from the trast-2 dataset. For the first baseline, we trained a deep language model with an identical architecture but no pre-training (i.e. randomly-initialized weights) to evaluate the impact of transfer learning. For the second baseline, we trained gradient boosted trees using the XGBoost package [23] to determine if deep language models boosted predictive accuracy relative to “shallow” machine learning. The pre-trained model out-performed both baselines for both the trast-1 and trast-2 datasets (fig. S7), with a stronger benefit seen for the smaller trast-2 dataset, in line with previous observations [24].

To understand why pre-training improves model performance, we inspected model embeddings from all combinations of pre-training vs. no pre-training, and fine-tuning vs. no fine-tuning (fig. S8). Even without fine-tuning, embeddings from OAS pre-training appear to have structure, with distinct patches enriched for high (or low) binding affinities. This organization simplifies subsequent fine-tuning with binding data, such that the model weights could be more easily updated to provide enhanced binding affinity predictions.

### Model-guided design of improved antibody variants

Having demonstrated AI prediction performances using hold-out sets and cross-validation, we moved to using models to design sets of sequences with desired binding properties followed by validation with dedicated SPR experiments.

To begin, we tasked a model trained on the trast-2 dataset with designing 50 sequences spanning two orders of magnitude of equilibrium dissociation constants (design set A). This model-enabled design involved exhaustively making predictions for all variants in the combinatorial sequence space, followed by sampling of sequences with predicted binding affinities consistent with requirements. We found an excellent agreement between the predictions and validations for the design set A (fig. 4A, fig. S9A-B).

**Figure 4.**
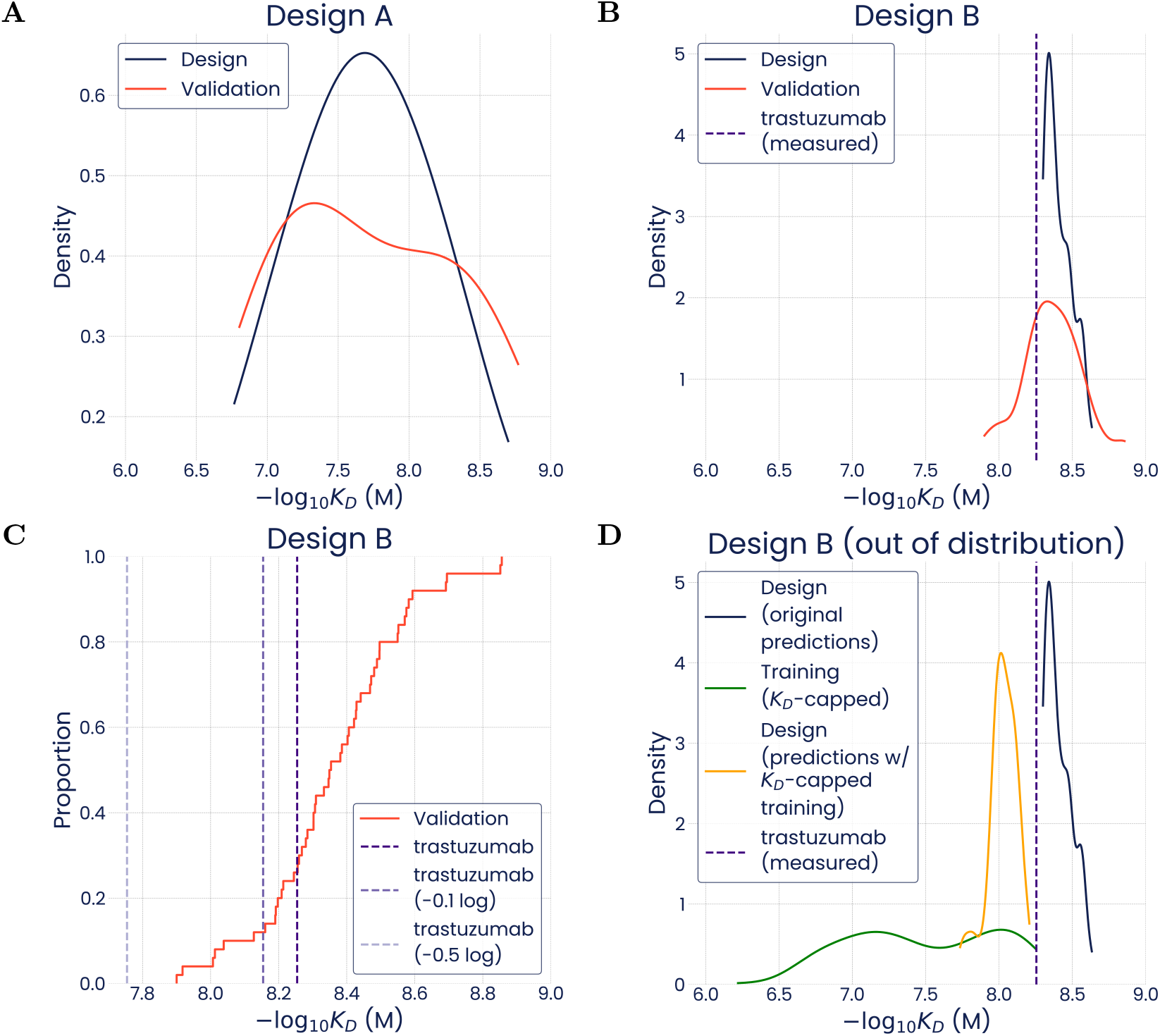
Deep language models trained with the SPR-generated trast-2 dataset can design unseen sequence variants that validate in independent SPR experiments. **(A)** Density plot of predicted (Design) and measured (Validation) binding affinities of 50 sequences designed to span about 2 orders of magnitude of *K*_*D*_s (set A). **(B)** Density plot of predicted (Design) and measured (Validation) binding affinities of 50 sequences designed to bind HER2 more tightly than parental trastuzumab (set B). **(C)** Empirical distribution function (ECDF) of the measured (Validation) binding affinities of the 50 sequences from design set B. Lines indicate the measured log_10_ *K*_*D*_ of trastuzumab (or deviations by −0.1 or −0.5 log). **(D)** Density plot of binding affinities from set B as predicted by a model trained with the full trast-2 dataset as in panel **B** (Design, original predictions) or as re-predicted (Design, predictions with *K*_*D*_-capped training) by a model trained on a trast-2 dataset version depleted of any variant binding more strongly than parental trastuzumab (Training, *K*_*D*_-capped).

We then considered a more challenging case, the design of variants with tighter binding than trastuzumab (design set B). As in the previous design, we validated 50 sequences by SPR and found that 74 % of variants were indeed tighter binders than the parental antibody (fig. 4B-C), and 100 % complied with the design specification within a tolerance of 0.5 log (fig. 4C, fig. S9C). This performance is competitive when considering replicate SPR measurements; a similar fraction of top binders from one replicate pass the threshold in the next (fig. S9D).

Because of the small - log_10_ *K*_*D*_ range spanned in this design, correlation between predictions and measurements was low (fig. S9C). As similarly observed in k_*on*_ modeling (fig. S4), if the affinity range is narrow, even measurement replicates correlate poorly with each other (fig. S9D). In contrast to correlation, other metrics, such as RMSE and the fraction of predictions deviating less than 0.5 log from measurements, remained in line with previously observed performance (fig. S9C). These metrics are generally more informative when considering sequences within a narrow affinity range.

The validation results for design set B compare very favorably against a naive, wet-lab-only approach to library screening, in which the fraction of binders tighter than trastuzumab is minimal (fig. S10). The strong enrichment for variants of interest provided by AI models can thus greatly facilitate antibody optimization (fig. 1).

As mentioned above, the model used to design sequence set B was trained on the trast-2 dataset, which included some binders stronger than trastuzumab (fig. 3A). We investigated whether a model that was never fed any sequence as extreme (affinity-wise) as those it was tasked to design could still prioritize top binders. This question is of practical value, as some campaigns may start from training sets devoid of high-affinity sequences. To test the performance of our models in out-of-distribution affinity prediction, we dropped those sequences with higher affinity than trastuzumab from the trast-2 training set. We then trained a model using the remaining data and predicted the affinity of sequences in the design set B. We found that the model was no longer able to make accurate *K*_*D*_ predictions for design B. Nonetheless, the model did place the binding affinities of design B variants at the top of its predictive distribution (fig. 4D). This result demonstrates that AI can enable the prioritization of high-affinity sequences even if laboratory experiments generating training data did not span the full affinity range.

### AI predictive performance is maintained when scaling to a larger sequence space

To evaluate the accuracy of predictions in a large sequence space, we performed combinatorial mutagenesis of up to three simultaneous mutations in CDRH2 and CDRH3, ten positions each. We constructed a library by sampling less than 1 % of this sequence space, and measured the binding affinity of the sampled sequence variants using the ACE assay (trast-3, table 1, fig. 5A). We then trained a model using 80 % of the trast-3 data, and evaluated its performance on the remaining 20 % of hold-out sequences. The model predictions were accurate (fig. 5B). As a negative control, we confirmed that a model trained on a dataset with randomly shuffled ACE scores had no predictive power (fig. S11). Since the trast-3 sequence space is vast and we routinely observe a high correlation between ACE scores and SPR-measured- log_10_ *K*_*D*_ values (e.g., fig. 2D), models were trained and evaluated directly on ACE data.

**Figure 5.**
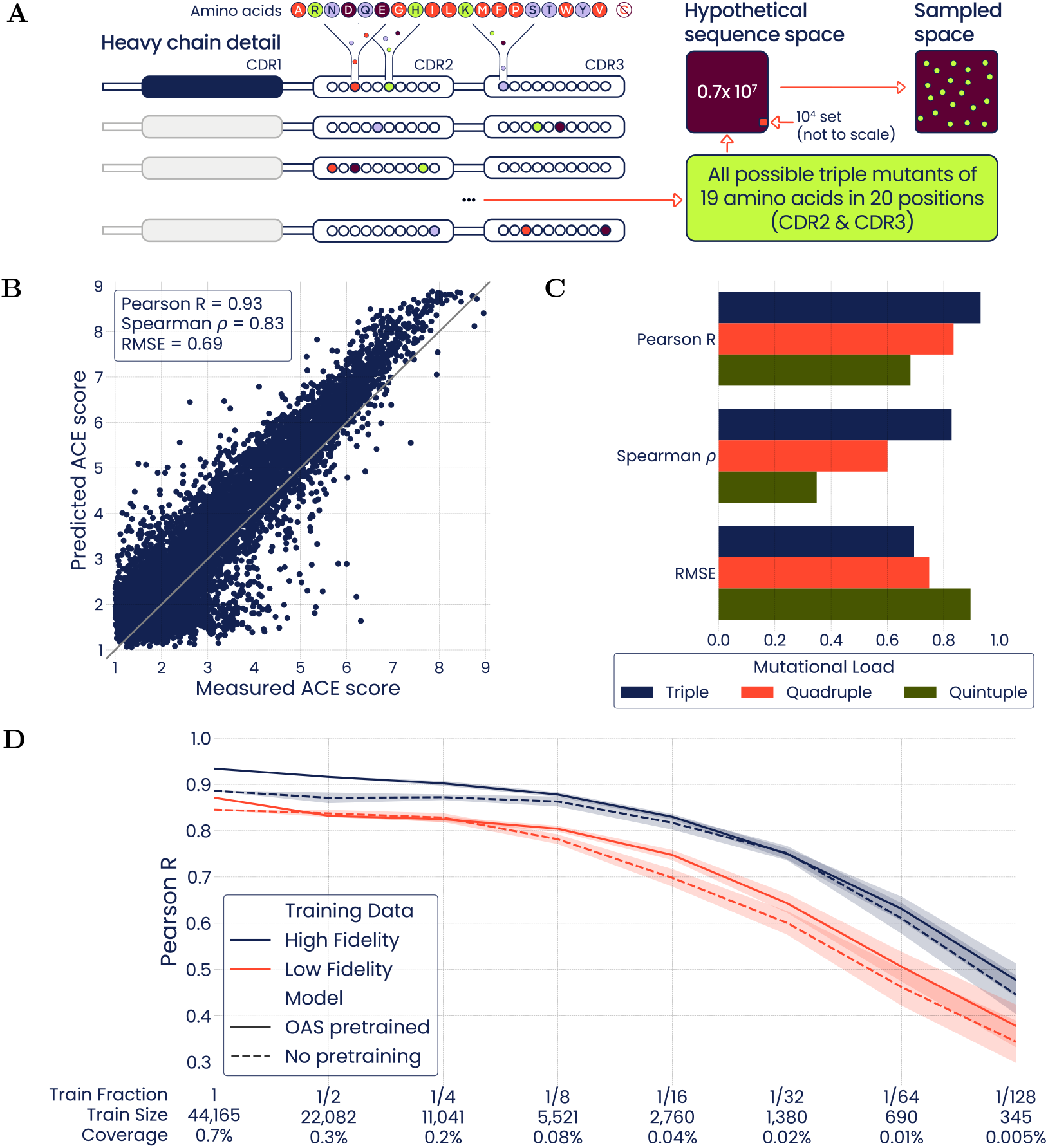
High-throughput binding scores from the ACE-generated trast-3 dataset can expand predictive capabilities to a larger mutational space. **(A)** Illustration of the combinatorial mutagenesis strategy of the trast-3 dataset: up to triple mutants in 20 positions (10 in CDRH2, 10 in CDRH3) of trastuzumab, screened using the ACE assay. **(B)** Predictive performance of a model trained on the trast-3 dataset, with 20 % of data in the hold-out set. **(C)** Models trained on up to triple mutants were validated against a hold-out set of up to triple mutants, and against hold-out sets of quadruple and quintuple mutants, thereby extrapolating predictions to a higher mutational load than seen in the training set. **(D)** Line plot showing model accuracy on a common hold-out validation set across different training set sizes. Shaded regions indicate standard deviations across folds. For each training subset size, we show the performance of the OAS pre-trained model and a randomly-initialized model, each trained using subsets of the high-fidelity trast-3 dataset or a low-fidelity version of the dataset. Under each subset size, we report the fraction of training data used, the size of the training dataset, and the fraction of the sequence space covered by the training subset.

Given the predictive accuracy of the trast-3 model on variants with up to three mutations away from the trastuzumab sequence, we tested whether the model could accurately predict the ACE scores of variants with four or five mutations (fig. 5C). The model predicted ACE scores of quadruple mutants (fig. S12A) with slightly lower (but still actionable) accuracy than those of triple mutants. Although the prediction accuracy for quintuple mutants was much lower (fig. S12B), the model could still discriminate between high- and low-affinity binders. These results show that the triple mutant model can be extrapolated to quantitatively predict binding scores for up to four simultaneous mutations, and qualitatively predict binding scores for five mutations from the parental sequence.

### Deep language models are highly sample efficient

The predictive power of any deep learning model is highly dependent on the quality and quantity of its training data. The trast-3 dataset contains binding affinities for about 50,000 unique antibody sequences, covering 0.7 % of the complete combinatorial sequence space for this design (table 1). To determine the relationship between model performance and the quality and quantity of the training dataset, we trained a cohort of models to predict affinity from a range of dataset sizes sampled from datasets of varying fidelity (fig. 5D). We treated the original trast-3 dataset as a high-fidelity dataset, and created a low-fidelity dataset by isolating a single DNA variant for each protein sequence from a single FACS sort replicate (see Methods). The size of the training subsets ranged from 44,165 sequences (the full training dataset), through 350 sequences (1/128 of the full training dataset), and models were evaluated on a common hold-out validation dataset containing 10 % of all sequences in the high-fidelity dataset. At each training subset size, we compared the performance of four models: *(1)* OAS pre-trained models trained on a subset from the high-fidelity dataset; *(2)* OAS pre-trained models trained on a subset from the low-fidelity dataset; *(3)* randomly-initialized models trained on a subset from the high-fidelity dataset; and *(4)* randomly initialized models trained on a subset from the low-fidelity subset.

As the size of the training dataset decreased, the model performance degraded. Models trained on low-fidelity data consistently performed poorer than their counterparts trained on high-fidelity data, highlighting the importance of high-quality experimental assays. Pre-training the model with immunoglobulin sequences from the OAS dataset generally improved its performance (fig. 5D). Given that the model required at least 2,760 sequences to maintain a Pearson’s R above 0.8, it is impractical to model this (or larger) sequence space using only SPR training data; higher-throughput assays such as the ACE assay are required.

Since the Pearson correlation coefficient remained above 0.8 for all high-fidelity training subsets covering at least 0.04 % of the potential search space, the model learned to predict roughly 2,500 sequences for every sequence in the training set. Therefore, deep language models can expand the search space of an experimental dataset by orders of magnitude.

### Deep language models enable interpretable analysis of the antibody binding landscape

Once trained, deep neural networks can be used as oracles to predict binding affinity scores for all sequences within the combinatorial space matching the design of the training set. Fast and accurate predictions of how antibody properties would be affected by sequence engineering can help guide design strategies.

To gain insight into the binding landscape of trastuzumab variants, we exhaustively evaluated the effect of all single, double and triple mutations in CDRH2 and CDRH3.

Trastuzumab has a high binding affinity for its target antigen HER2 (- log_10_ *K*_*D*_ of 8.25 M in Fab format, see fig. 3A). Thus, most mutations were predicted to have a detrimental effect on the binding affinity (fig. 6). When considering multiple mutations, we also found that most combinations were predicted to have a detrimental effect on the binding affinity (fig. S13). In particular, positions 55, 107, 111, 112, and 113, were often predicted to have a detrimental effect when mutated (fig. S13) and tended to interact epistatically with other mutations. This pointed to a strong contribution to binding affinity from these residues, in agreement with previous alanine scanning and structural studies [22].

**Figure 6.**
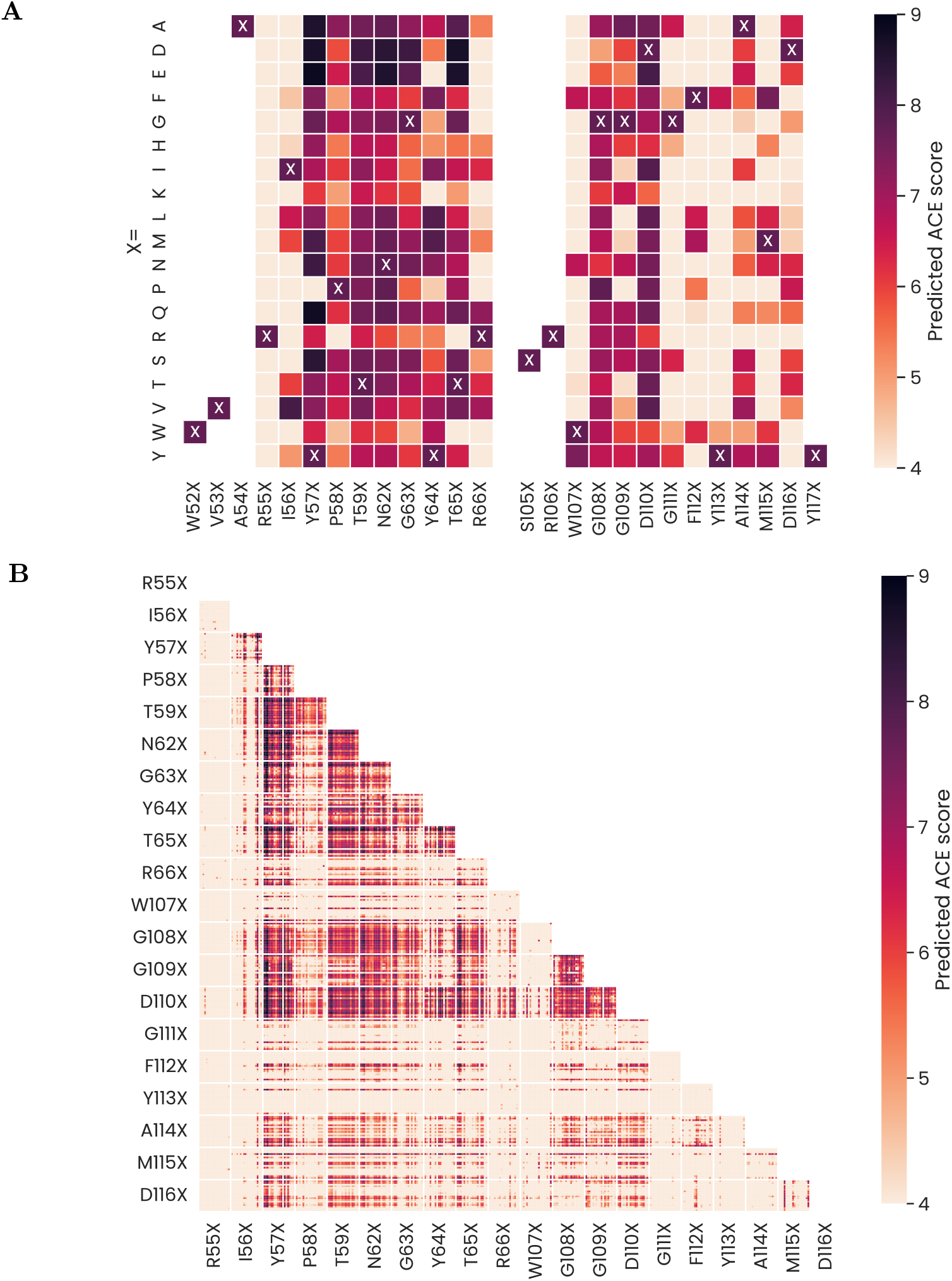
Global sequence-affinity mapping of trastuzumab variants. Predicted binding affinities for **(A)** single or **(B)** double mutants from a model trained on the trast-3 dataset. Positions holding mutations comprised CDRH2 (10 positions starting with R55) and CDRH3 (10 positions starting with W107). The reference trastuzumab sequence is highlighted with crosses. Mutations at each position include all possible substitutions with natural amino acids except cysteine, sorted alphabetically (i.e., *X* ∈ [*A, D, E, F, G, H, I, K, L, M, N, P, Q, R, S, T, V, W, Y*]).

Analyzing the incremental effects of mutations across variants revealed that positions 59, 62, and 110 were relatively tolerant to mutations (fig. S13). This suggests that they make a relatively small contribution to binding, and may offer ideal handles to optimize other antibody properties without perturbing affinity for the antigen.

Some single mutations in CDRH2, such as Y57D/E, N62E or T65D/E, were predicted to increase binding affinity (fig. 6). Beyond single mutants, combining multiple mutations may also provide improved high-affinity variants. In fact, as the mutational load increased, the number of predicted high-affinity sequences increased, although their proportion was reduced. For instance, 2 (0.56 %) of the single mutants, 192 (0.31 %) of the double mutants, and 7,063 (0.11 %) of the triple mutants had high (>8.7) predicted ACE scores in the trast-3 dataset.

We carried out a clustering analysis of model-derived embeddings of high-affinity sequences (predicted ACE score >8.0). While the space of triple mutants offered many potential high-affinity candidate sequences, these tended to form compact clusters involving specific substitutions in a few positions, as shown in fig. S14. Notably, mutation Y57D/E was observed in several clusters. Also, most high-affinity triple mutants had two or three mutations in the CDRH2 (particularly in positions 57 and 62 or adjacent positions), while fewer solutions involved one mutation in CDRH2 and two mutations in CDRH3. This finding highlights the key role of the CDRH2 region in antigen binding by trastuzumab, as also noted by others [22, 25].

We also found that the impact of a given mutation on binding affinity varied widely with the presence of other mutations in the sequence, a phenomenon known as *contingency* [26]. In fig. S13, we observed that a given mutation can have a larger, smaller, or even opposite effect compared to the effect it would have on the parental trastuzumab sequence, depending on the presence of just another single mutation. In the presence of two mutations, the possible range of effects for an additional (third) mutation became wider (fig. S13).

In a similar vein, *epistasis* is the deviation from additivity in the effects of two co-occurring mutations compared to their individual effects [27]. The epistatic interaction between mutations for all double mutants of trastuzumab is depicted in fig. S15. Given the negative effect that many mutations had on binding affinity, antagonistic, positive epistasis is often observed (i.e., a double mutant displays a higher binding affinity than expected based on its constituent single mutants). This is particularly evident in pairs of mutations involving positions 55, 107, 111, 112, and 113, which are crucial to the binding affinity of trastuzumab [22]. Epistatic interactions are also highly contingent on the presence of other mutations in the sequence. The complex interaction between mutations directly affects the biochemical properties of antibodies.

Taken together, the diversity of high-affinity sequences and their dilution as a function of mutational load highlights the value of exhaustively evaluating the space of possible variants. Such large-scale evaluation is only feasible with the help of computational models. Our modeling results are in excellent agreement with previous functional and structural studies and can provide unique insight on how mutations interact to shape the binding affinity of antibodies. The pervasiveness of epistatic effects also highlights the need for flexible AI models to accurately guide antibody optimization.

### AI shows strong predictive performance on a second case study involving simultaneous binding predictions for three antigen variants

Our modeling approach established with trastuzumab can be readily extended to other antibodies. To demonstrate this, we leveraged public binding data of variants of the broadly neutralizing (bn) antibody CR9114 (see Supplementary Information) [28]. Since the bnAb CR9114 dataset provides binding data for three different influenza subtypes of the target antigen hemagglutinin (HA), we extended the model to support multi-task affinity predictions to multiple targets simultaneously. We also explored the ability of the model to combine classification and regression in a single mixture model, since many of the CR9114 variants lost binding to one or more HA subtypes. Lastly, we evaluated the impact of the training set size on the model performance.

Results (table 2 and figs. S16 to S21) showed that a single model could be trained to jointly predict affinities of a given antibody sequence against multiple distinct antigen targets. As expected, the predictive power of the model was lower for the FluB target compared to H1 and H3, since the full dataset contains only 193 positive FluB binders. This left only 19 positive examples when using a training set of 10 % and only 1-2 positive examples in training sets of 1 % and 0.1 % (a minimum of one positive and negative example for each target was required when selecting the cross-validation folds, see Supplementary Information). Nevertheless, even with as little as 19 training examples, 88 % of FluB predictions were within 0.5 log of their measured values when using initial weights pre-trained on the OAS dataset, compared to only 73 % when using random initial weights. Using pre-trained weights improved performance in all cases where the number of training examples was below 1,000. The mixture model was able to perform well on the classification tasks without significant loss of performance on the regression tasks compared to the regression only model. The balanced accuracy of the model predictions was above 0.84 in all cases where the training set contained at least 7 positive and 7 negative examples, achieving a 0.91 balanced accuracy score on the H3 binding task even with training sets of only 65 variants (7 positive and 58 negative variants on average).

**Table 2.**
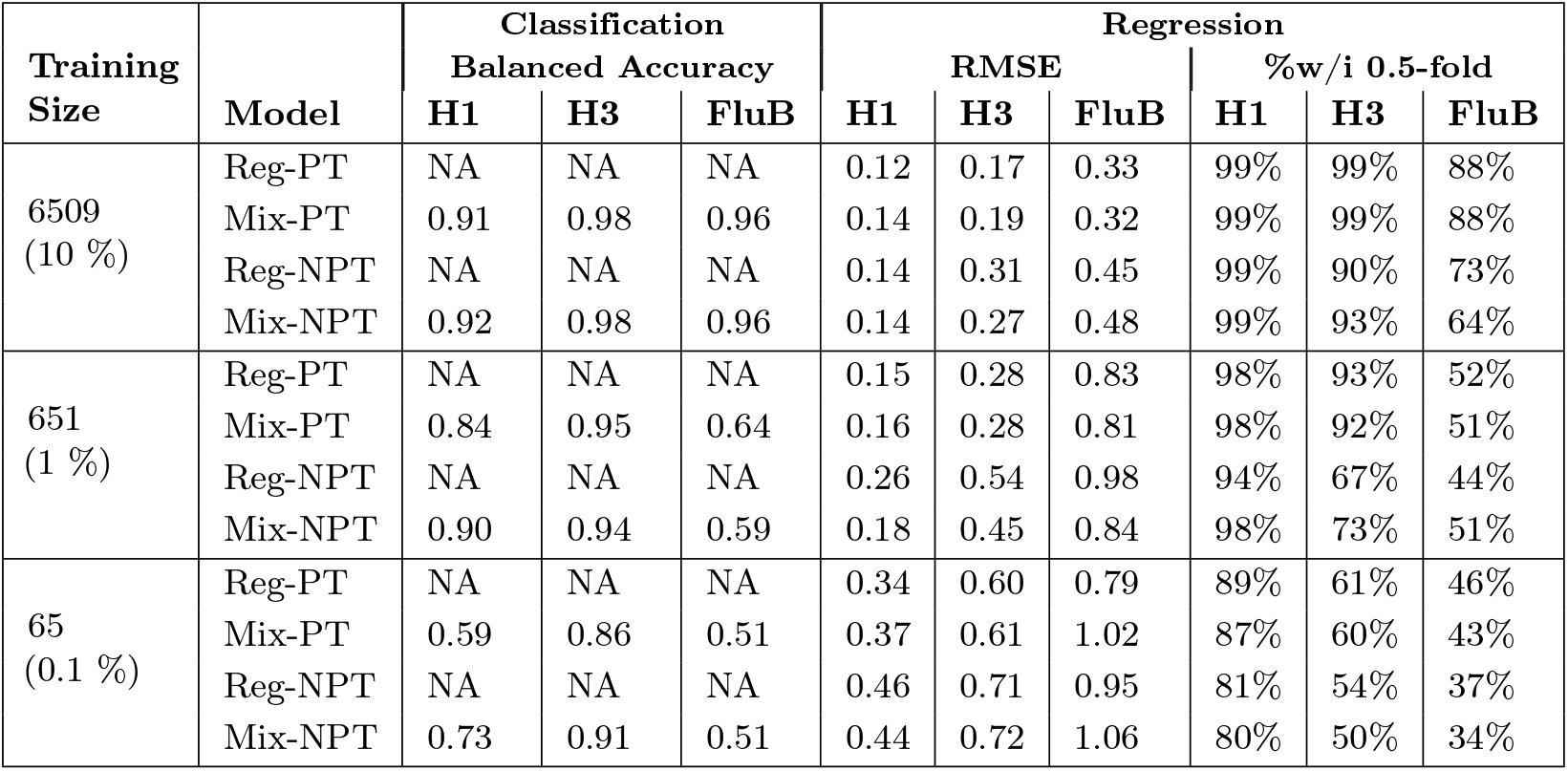
Joint model affinity prediction performance for CR9114 on multiple influenza strains of the hemagglutinin (HA) antigen. For each training set size (10 %, 1 %, 0.1 % of 65,091) four models were trained (*Reg* : Regression only model; *Mix* : Mixture classification/regression model; *PT* : initialized with pre-trained OAS-model weights; *NPT* : initialized with random weights). Results are shown for these models using pooled CV. The full CR9114 dataset includes 63,419 (97 %) H1, 7,174 (11 %) H3, and 198 (0.3 %) FluB positive binders.

### Optimizing antibody naturalness may mitigate development hurdles

The development of a candidate antibody into a therapeutic drug is a complex process with a high degree of pre-clinical and clinical risk. This risk is often due to numerous challenges related to production, formulation, efficacy and adverse reactions. Modeling these risks has been a tremendous challenge for the industry due to the difficulty in obtaining informative, abundant and relevant data.

We hypothesized that learning sequence patterns across natural antibodies from different species could be useful to identify and prioritize “human-like” antibody variants (as opposed to unnatural sequences) and, ultimately, mitigate drug development risks (fig. 7). To this aim, we took advantage of our OAS pre-trained language models to evaluate antibody sequences for their *naturalness* (see Methods). We define naturalness as a score computed by pre-trained language models that measures how likely it is for a given antibody sequence to be derived from a species of interest, including human. Thus, naturalness might be used as a guiding metric in antibody design and engineering.

**Figure 7.**
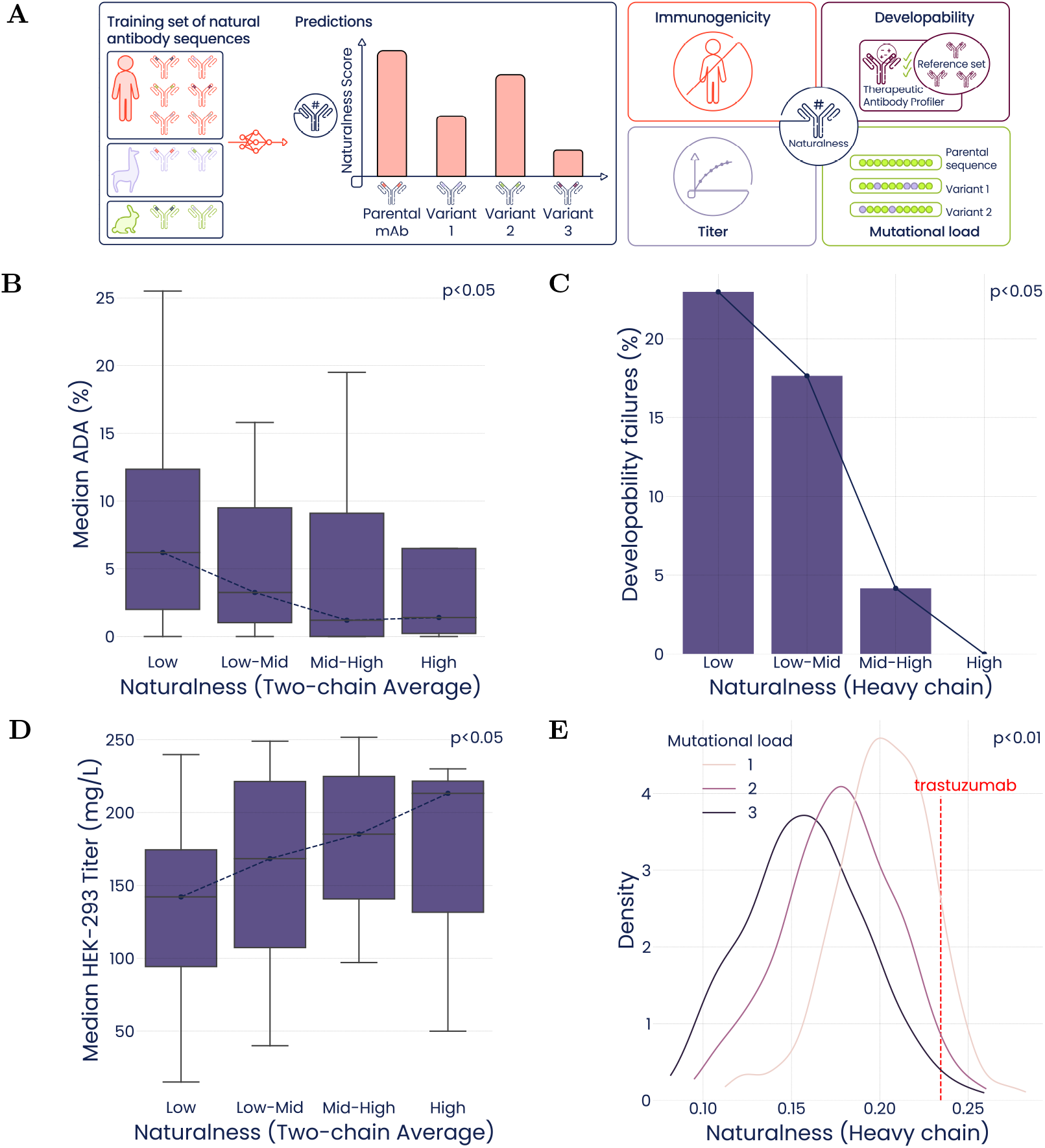
Associations between antibody naturalness, immunogenicity, developability and other properties. **(A)** Language models pre-trained with antibody repertoire sequences can be leveraged to compute the naturalness of an antibody sequence conditioned on a given species. Naturalness scores were investigated for association with four antibody properties: **(B)** *Immunogenicity* using Anti-Drug Antibody (ADA) responses to humanized clinical-stage antibodies reported by Marks et al. [29] (n=97); **(C)** *Developability* failures as predicted by the Therapeutic Antibody Profiler (TAP) for round 3-enriched phage display hits from the Gifford library [30] (n=882); **(D)** *Expression levels* in HEK-293 cells (mg/L) of clinical-stage humanized antibodies from Jain et al. [31] (n=67); **(E)** *Mutational load* of trastuzumab variants using a mutagenesis strategy as the trast-3 dataset (n=6,710,400). The dashed line corresponds to the naturalness of the parental trastuzumab sequence. In all box plots, the four bins (Low, Low-Mid, Mid-High, High) result from dividing the naturalness range into four parts of equal size (see fig. S23). In all panels, p-values were computed using the Jonckheere-Terpstra test for trends across the four bins going from Low to High. Datasets in panels **B** and **D** were scored using the average naturalness of both the heavy and light chains, whereas datasets in panels **C** and **E** comprised only heavy-chain variants and were consistently scored only with the heavy-chain models.

To determine the usefulness of naturalness, we evaluated its association with four antibody properties (fig. 7). The first property studied was immunogenicity, for which Anti-Drug Antibodies (ADA) responses were collected from numerous primary studies on clinical-stage antibodies by Marks et al. [29]. A potential confounding factor in a naturalness-immunogenicity association analysis is that some antibodies have a fully human origin, while others are humanized, chimeric or murine. Scoring antibodies of different origin by naturalness would amount to binning them primarily by species, which would be trivial and uninformative. By contrast, scoring antibodies belonging to the same class would amount to genuinely ranking from most natural to least natural. The only two antibody classes in Marks et al. large enough to support a statistical analysis are human and humanized antibodies. We investigated the latter because their reported immunogenicity is greater [29], thereby providing an ideal case study.

A scatterplot of the fraction of ADA-positive patients vs. naturalness scores reveals a weak, non-significant correlation (fig. S22). However, closer inspection of ADA responses showed that most data points are in the 0-10 % range, with a few outliers above 20 %. We reasoned that outliers could blur the relationship between naturalness and immunogenicity. To mitigate the impact of outliers, we binned naturalness scores (fig. S23A) and computed the median ADA responses per naturalness bin. This analysis revealed that antibodies with higher naturalness scores trigger lower median ADA responses than less natural antibodies (fig. 7B).

The second property considered was developability, which can be estimated with the Therapeutic Antibody Profiler (TAP) [32]. We computed naturalness scores (with our model) and developability scores (with TAP) for the heavy-chain sequences from a high-diversity phage display dataset [30] (“Gifford Library”, fig. S23B) as well as for trastuzumab variants (fig. S23C). In both cases, we found a strong association between naturalness and TAP-determined developability (fig. 7C, fig. S24). This is a remarkable result because naturalness scores were obtained upon training exclusively with examples of naturally occurring antibody sequences, while TAP was calibrated using distributions of five metrics computed on therapeutic antibodies [32]. The association between naturalness and TAP flags suggests that developable antibodies are enriched in human-like antibodies.

The third property investigated was antibody expression level in mammalian (HEK-293) cells, which has been reported for clinical-stage antibodies by Jain et al. [31]. As for immunogenicity, the dataset comprises several classes of antibodies and we again focused on humanized antibodies. We found that antibodies with high naturalness scores were expressed at higher levels than antibodies with low scores (fig. 7D, fig. S23D).

The fourth property considered was mutational load, which is the number of amino acid substitutions in a variant compared to a parental antibody sequence. We computed naturalness scores for 6,710,400 single-, double-, and triple-mutant trastuzumab variants (fig. 7E). We found that naturalness was negatively associated with mutational load. This is consistent with the observation that most mutations have detrimental effects. Since the introduction of mutations can degrade naturalness, it is important to simultaneously optimize naturalness and binding affinity of antibodies.

### Sequence variant generation with desired properties

Antibody optimization can be performed to a limited extent for individual properties using a number of established laboratory approaches. For example, deep mutational scanning has been used to improve the binding affinity of antibody candidates [5]. However, large mutational spaces cannot be exhaustively screened by these methods, limiting the scope of potential improvements. Library screening methods, such as phage display, can overcome this obstacle, but selecting for a single property at a time (such as binding affinity) may negatively affect other properties of interest [19]. For example, we showed that increasing the mutational load often lowers naturalness (fig. 7D).

We exhaustively predicted ACE and naturalness scores of all variants with up to three mutations from trastuzumab. Of the 6.7 million variants, just 46,931 (0.7 %) had predicted ACE scores higher than trastuzumab (fig. S25A). Of these, only 4,003 (8.5 %) had a naturalness score on par or higher than trastuzumab (fig. S25B). Randomly screening this space using the approximately 50,000-member trast-3 library yielded only 60 variants with higher ACE scores and naturalness than trastuzumab.

*In silico* screening provides a way to address this issue by optimizing for multiple properties simultaneously with a designer objective function. We built a genetic algorithm (GA) on top of our affinity and naturalness model oracle that was capable of greatly improving the throughput of our *in silico* screening process. As an example, we could minimize, maximize, or target specific ACE scores in a search space of over 6.7 million sequence variants (fig. 8A), while simultaneously maximizing naturalness (fig. 8B).

**Figure 8.**
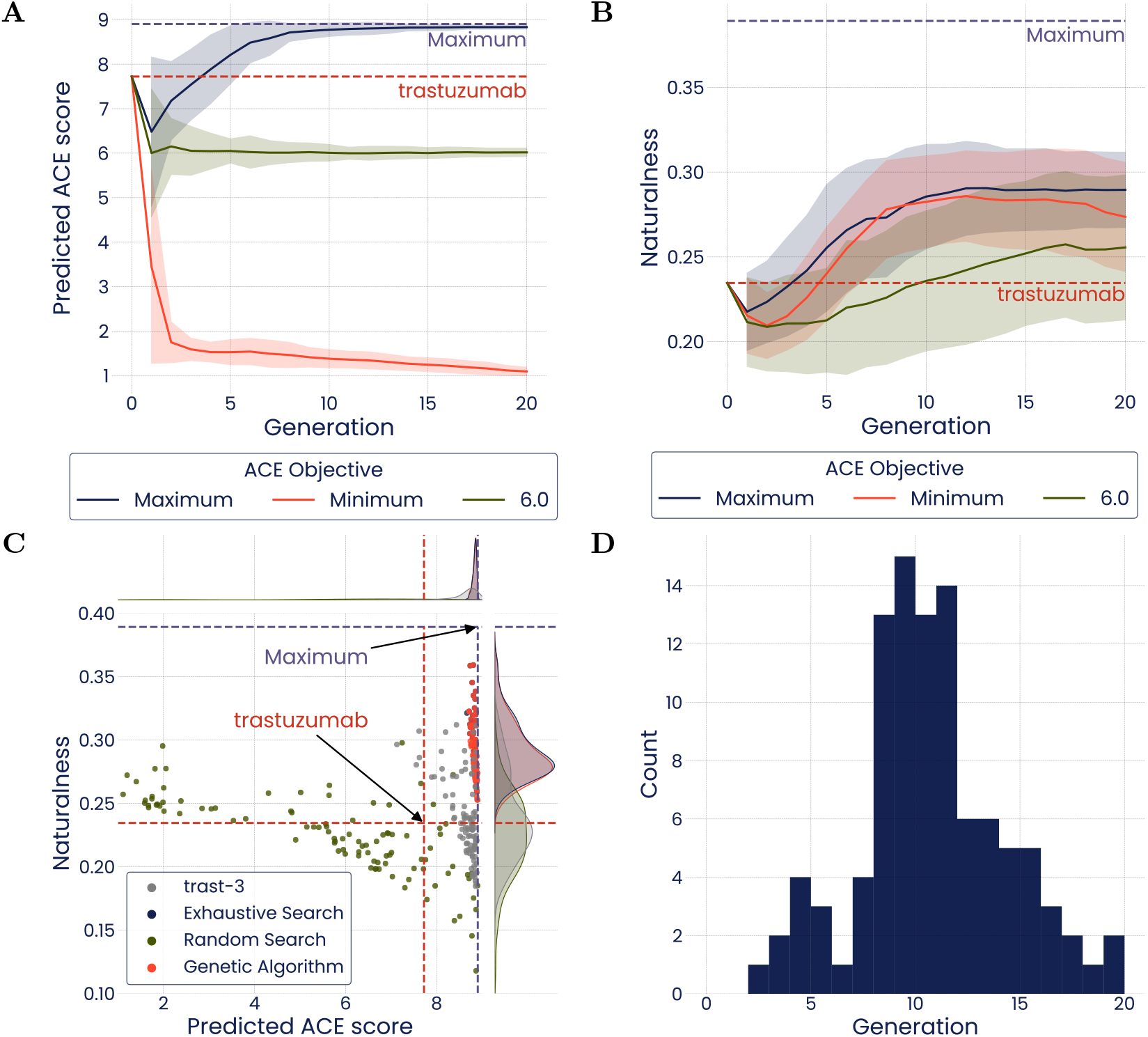
A genetic algorithm can efficiently maximize, minimize, or target specific ACE scores while maximizing naturalness. **(A)** Each line tracks the average predicted ACE score of the best 100 sequences observed across the evolutionary trajectory. Shaded regions indicate the standard deviation. **(B)** Average naturalness of the best 100 sequences observed across the evolutionary trajectory. Shaded regions indicate the standard deviation. **(C)** ACE and naturalness scores of the best 100 sequences determined through three search strategies: Genetic Algorithm, Exhaustive Search, and Random Search. Red dashed lines indicate the scores predicted for trastuzumab. Purple dashed lines indicate maximum scores predicted across the entire combinatorial space. **(D)** Histogram showing the first generation where each of the top 100 sequences observed along the evolutionary trajectory was identified.

After 20 generations, the GA performed nearly as well as an exhaustive search of the mutational space (fig. 8C); 85 of the top 100 variants identified by the GA were among the top 100 variants overall. In addition, all of the top variants identified by the GA were within 5 % of the maximum achievable ACE score (9, resulting from 9 sorting gates) and had higher naturalness scores than trastuzumab. As a baseline, we performed a random search by querying the same number of sequences as the GA. This search was only able to find two sequences with higher ACE score and naturalness than trastuzumab (fig. 8C).

Unlike an exhaustive search of the mutation space, GA-driven optimization is highly efficient. In each generation, the GA sampled 200 new variants, resulting in only 4,000 total sequences sampled across all 20 generations. In addition, over half of the top 100 individuals were selected by the GA in the first 12 generations (fig. 8D). Altogether, these results show that a genetic algorithm built on top of predictive models for binding affinity and naturalness can quickly and efficiently identify a set of top candidates for downstream development. The value of optimization techniques coupled with AI oracles will likely increase as *in silico* design is applied to larger combinatorial sequence spaces.

## Discussion

Deep learning methods have demonstrated rapid progress for the modeling of proteins, including their sequence, structure, and function. Likewise, protein interactions are receiving increased attention for the purposes of therapeutic design. A key limitation for many of these efforts is the ability to synthesize large libraries of proteins and quantitatively assess their attributes. Here we demonstrate that our ACE assay is a powerful complement to deep learning models, providing the throughput and fidelity needed to accurately model antibody binding affinity with up to four/five mutations across two CDRs (combinatorial space: 10^8^-10^10^) in a single experiment. The ACE assay provides advantages over existing methods for large scale antibody variant interrogation such as Tite-Seq [33], SORTCERY [34] and Phage Display [35]. First, the ACE assay utilizes the SoluPro™ *E. coli* B Strain to solubly express antibodies intracellularly, avoiding binding artifacts associated with surface display formats. Additionally, the ACE assay leverages genetic tools available for *E. coli*, enabling faster library generation cycles and increased transformation efficiency compared to other organisms. Finally, the ACE assay is a true screening method where all variants are measured regardless of affinity strength, as opposed to selection methods, such as phage display, where only high-affinity binders are preferentially isolated.

The predictive ability of our deep learning models demonstrated here is enabled by the quantitative data generated by our improved ACE assay, which provides two distinct advantages from a modeling standpoint. The first is the expanded capabilities of models trained on quantitative data for overall increased performance and quantitative predictions, which are particularly useful when the goal is to tune the binding affinity rather than simply maximize it. Secondly, quantitative training data allows for the intelligent selection of sequences for downstream quantification with lower-throughput assays, such as SPR. The sequence space available for bioengineering is enormous and heavily skewed toward deleterious mutations. A common approach to this problem is to bias the mutational library towards specific locations or key mutations, but the strength of epistatic effects identified by our models suggest these approaches systematically miss potentially impactful sequence changes. Our pre-quantification step with the improved ACE assay allows us to access sequences throughout the binding affinity spectrum without bias, which increases the generalization power of the models.

Only a very small fraction of antibody sequences within the enormous combinatorial space have been detected in nature; there are >10^8^ high-quality, unique sequences in the OAS database versus more than 10^120^ possible unique CDR sequences for the longest reported human sequences. Our naturalness model can help determine whether a novel sequence belongs to this category, and we roughly estimate the size of this natural space as 10^60^ (fig. S30). While this estimate has considerable uncertainty, it is clear that the natural space is much larger than can possibly be screened in a lab or *in-silico*. At the same time, these natural sequences are vanishingly rare in screens of random sequence variants. The solution we present here is to apply models trained on both naturalness and affinity data, the intersection of which effectively allows evaluation of a larger whitespace of sequences than can be physically assessed, while also focusing screening on the most relevant “natural” sequences.

In future work, our co-optimization of two antibody properties could be extended to the co-optimization of additional properties relevant to protein interactions and therapeutic potential. Training models on multiple affinity datasets unlocks binding predictions for multiple antigens or antigen variants, as we showed here for CR9114. In principle, multi-antigen predictions could facilitate engineering of breadth (co-optimization for antigen escape variants), specificity (co-optimization to reduce binding to undesired members of a protein family, while increasing binding to desired members of the same family) and species cross-reactivity (co-optimization for human and cynomolgus orthologs), just to name a few.

We demonstrated that pre-training on natural sequences improved the predictive performance of our models. Likewise, the models can continue to improve with the addition of new data, both with respect to new antibodies and with the addition of new performance or developability attributes. Naturalness can be computed extremely rapidly and can complement other scores, with the potential to reduce preclinical or clinical attrition caused by complex properties such as immunogenicity. Additional properties could be added alongside data from their respective assays, such as conditional pH binding, effector function, melting temperature, self-aggregation, viscosity and more. For most of these, a single model trained on a high-quality dataset could serve for diverse antibodies of interest and even improve the power of the binding affinity models through multi-task learning. Importantly, the framework presented facilitates tuning an antibody property toward a desired specification, not necessarily limited to selecting for variants at the extremes of a given range. Moreover, while the models we presented are focused on target affinity and naturalness of antibodies, the approach could in principle be extended to other protein classes.

While AI-assisted optimization of biological sequences can reduce therapeutic development time, it does not by itself offer a fully *in-silico* replacement. To this end, fully generative modeling approaches are needed. However, their training and validation faces an even greater data challenge, since the full *de novo* combinatorial space considered without the anchor of the parental sequence is dramatically larger, and strong selective binders are an infinitesimally small slice of that space. Structure-based approaches are showing increasing capabilities and may be useful for bridging this gap. The language models presented here can serve as *in-silico* oracles within their applicability domain, and might provide an effective training ground for generative models. Harmonizing antibody optimization and *de novo* generation may be the next big step in data-driven therapeutic design.

## Materials and Methods

### Libraries

#### Cloning

Antibody variants were cloned and expressed in Fab format. To produce ACE and SPR datasets meant for model training and evaluation (table 1), we synthesized DNA variants spanning CDRH2 and CDRH3 in a single oligonucleotide using ssDNA oligo pools (Twist Bioscience). Codons were randomly selected from the two most common in *E. coli* B strain [36] for each variant. Two synonymous DNA sequences were synthesized (5 or 10 for parental trastuzumab and positive/negative controls) for each amino acid variant.

Amplification of Twist Bioscience ssDNA oligo pools was carried out by PCR according to Twist Bioscience’s recommendations with the exception that Platinum SuperFi II DNA polymerase (ThermoFisher) was used in place of KAPA polymerase. Briefly, 20 µL reactions consisted of 1x Platinum SuperFi II Mastermix, 0.3 µM each of forward and reverse primers, and 10 ng oligo pool. Reactions were initially denatured for 3 min at 95 °C, followed by 13 cycles of: 95 °C for 20 s; 66 °C for 20 s; 72 °C for 15 s; and a final extension of 72 °C for 1 min. DNA amplification was confirmed by agarose gel electrophoresis, and amplified DNA was subsequently purified (DNA Clean and Concentrate Kit, Zymo Research).

To build libraries meant for SPR validation of model designs in independent experiments, oligonucleotides (59 nt) spanning CDRH3 and the immediate upstream/downstream flanking nucleotides were synthesized by Integrated DNA Technologies (IDT). Codon usage was identical for all variants, except at mutated positions. Olignoucleotides were pooled such that each oligonucleotide was represented in an equimolar fashion within the pool. This single stranded oligonucleotide pool was used directly in cloning reactions (see below) without prior amplification.

To generate linearized vector, a two-step PCR was carried out to split Absci’s plasmid vector carrying fab format trastuzumab into two fragments in a manner that provided cloning overlaps of approximately 30 nucleotides (nt) on the 5’ and 3’ ends of the amplified Twist Bioscience libraries, or 18 nt on the 5’ and 3’ ends of IDT oligonucleotides. Vector linearization reactions were digested with DPN1 (New England Bioloabs) and purified from a 0.8 % agarose gel (Gel DNA Recovery Kit, Zymo Research) to eliminate parental vector carry through. Cloning reactions consisted of 50 fmol of each purified vector fragment, either 100 fmol purified library (Twist Bioscience) or 10 pmol (IDT) insert, and 1x final concentration NEBuilder HiFi DNA Assembly (New England Biolabs). Reactions were incubated at 50 °C for either two hours (Twist Bioscience libraries) or 25 min (IDT library), and subsequently purified (DNA Clean and Concentrate Kit, Zymo Research). Transformax Epi300 (Lucigen) *E. coli* were transformed by electroporation (BioRad MicroPulser) with the purified assembly reactions and grown overnight at 30 °C on LB agar plates containing 50 µg/ml kanamycin. The following morning colonies were scraped from LB plates and plasmids were extracted (Plasmid Midi Kit, Zymo Research) and submitted for QC sequencing.

#### QC

Antibody variant libraries were amplified by PCR across the CDRH2 and CDRH3 region and sequenced with 2×150 nt reads using the Illumina NextSeq 1000 P2 platform with 20 % PhiX. The PCR reaction used 10 nM primer concentration, Q5 2x master mix (NEB) and 1 ng of input DNA diluted in MGH_2_0. Reactions were initially denatured at 98 °C for 3 min; followed by 30 cycles of 98 °C for 10 s, 59 °C for 30 s, 72 °C for 15 s; with a final extension of 72 °C for 2 min.

Sequencing results were analyzed for distribution of mutations, variant representation, library complexity and recovery of expected sequences. Metrics included coefficient of variation of sequence representation, read share of top 1 % most prevalent sequences and percentage of designed library sequences observed within the library.

### Activity-specific Cell-Enrichment (ACE) assay

#### Antibody Expression in SoluPro™ *E. coli* B Strain

SoluPro™ *E. coli* B strain was transformed by electroporation (Bio-Rad MicroPulser). Cells were allowed to recover in 1 ml SOC medium for 90 min at 30 °C with 250 rpm shaking. Recovery outgrowths were centrifuged for 5 min at 8,000 g and the supernatant was removed. Resultant cell pellets were resuspended in 1 mL of induction media (IBM) (4.5 g/L Potassium Phosphate 547 monobasic, 13.8 g/L Ammonium Sulfate, 20.5 g/L yeast extract, 20.5 g/L glycerol, 1.95 g/L Citric Acid) containing inducers and supplements (260 µM Arabinose, 50 µg/mL Kanamycin, 8 mM Magnesium Sulfate, 1 mM Propionate, 1X Korz trace metals) and then added to 100 ml IBM containing inducers and supplements in a 1 L baffled flask. Antibody Fab induction was allowed to proceed at 30 °C with 250 rpm shaking for 24 h. At the end of 24 h, 1 mL aliquots of the induced culture were adjusted to 25 % v/v glycerol and stored at −80 °C.

#### Cell Preparation

High-throughput quantitative selection of antigen-specific Fab-expressing cells was adapted from the approach described in Liu et al. [20]. For staining, an OD_600_ = 2 of thawed glycerol stocks from induced cultures were transferred to 0.7 ml matrix tubes, centrifuged at 3300 g for 3 min, and resulting pelleted cells were washed three times with PBS + 1 mM EDTA. Washed cells were thoroughly resuspended in 250 µL of 33 mM phosphate buffer (Na_2_HPO_4_) by pipetting then fixed by the further addition of 250 µL 32 mM phosphate buffer with 0.5 % paraformaldehyde and 0.04 % glutaraldehyde. After 40 min incubation on ice, cells were washed three times with PBS, resuspended in lysozyme buffer (20 mM Tris, 50 mM glucose, 10 mM EDTA, 5 µg/ml lysozyme) and incubated for 8 min on ice. Fixed and lysozyme-treated cells were equilibrated in stain buffer by washing 3x in 0.1% saponin buffer (1x PBS, 1 mM EDTA, 0.1 % saponin, 1 % heat-inactivated FBS).

#### Staining

Prior to library staining, the Her2 probe was titrated against the reference strain to determine the 75 % effective concentration (EC_75_). After lysozyme treatment and equilibration, the trast-1 library was resuspended in 250 µL saponin buffer and transferred to a new matrix tube. The trast-3 library was incubated for 20 min in AlphaLISA immunoassay assay buffer (Perkin Elmer; 25 mM HEPES, 0.1 % casein, 1 mg/ml dextran-500, 0.5 % Triton X-100, and 0.05 % kathon) for additional permeabilization prior to equilibration and resuspension in saponin buffer. A 2x concentration of stain reagents – 100 nM human HER2:AF647 (Acro Biosystems) and 60 nM anti-kappa light chain:AF488 (BioLegend) – was prepared in saponin buffer, then 250 µL probe solution was transferred to the prepared cells bringing the total stain volume to 500 µL with 50 nM Her2 and 30 nM anti-kappa LC. Libraries were incubated with probe overnight (16 h) with end to end rotation at 4 °C protected from light. After incubation, cells were pelleted, washed 3x with PBS, and then resuspended in 500 µL PBS by thorough pipetting.

#### Sorting

Libraries were sorted on FACSymphony S6 (BD Biosciences) instruments. Immediately prior to sorting, 50 µL prepped sample was transferred to a flow tube containing 1 mL PBS + 3 µL propidium iodide. Aggregates, debris, and impermeable cells were were removed with singlets, size, and PI^+^ parent gating. To reduce expression bias, an additional parent gate was set on the mid 65 % of peak expression positive cells. Collection gates were drawn to evenly sample the log range of binding signal. The far right gate was set to collect the brightest 10,000 events over the allotted sort time, estimated by including the 5 brightest events for every 65,000 in the expression parent gate. Seven additional gates were then set to fractionate the positive binding signal, and one gate collected the binding negative population (fig. S26). Libraries were sorted simultaneously on two instruments with photomultipliers adjusted to normalize fluorescence intensity, and the collected events were processed independently as technical replicates.

#### Next-generation sequencing

Cell material from various gates was collected in a diluted PBS mixture (VWR), in 1.5 mL tubes (Eppendorf). Post sort samples were spun down at 3,800 g and tube volume was normalized to 20 µl. Amplicons for sequencing were generated from one of two methods. The frist method amplifies the CDRH2 and CDRH3 region via a two-phase PCR, using collected cell material directly as template. During the initial PCR phase, unique molecular identifiers (UMIs) and partial Illumina adapters were added to the CDRH2 and CDRH3 amplicon via 4 PCR cycles. The second phase PCR added the remaining portion of the Illumina sequencing adapter and the Illumina i5 and i7 sample indices. The initial PCR reaction used 1 nm UMI primer concentration, Q5 2x master mix (NEB) and 20 µl of sorted cell material input suspended in diluted PBS (VWR). Reactions were initially denatured at 98 °C for 3 min, followed by 4 cycles of 98 °C for 10 s; 59 °C for 30 s; 72 °C for 30 s; with a final extension of 72 °C for 2 min. Following the initial PCR, 0.5 µM of the secondary sample index primers were added to each reaction tube. Reactions were then denatured at 98 °C for 3 min, followed by 29 cycles of 98 °C for 10 s; 62 °C for 30 s; 72 °C for 15 s; with a final extension of 72 °C for 2 min. The second method amplifies the CDRH2 and CDRH3 region without the addition of UMIs. This single phase PCR used 10 nM primer concentration, Q5 2x master mix (NEB) and 20 µl of sorted cell material input suspended in diluted PBS (VWR). Reactions were initially denatured at 98 °C for 3 min, followed by 30 cycles of 98 °C for 10 s; 59 °C for 30 s; 72 °C for 15 s; with a final extension of 72 °C for 2 min. After amplification by either method samples were run on a 2 % agarose gel at 75 V for 60 min and the proper length band was excised and purified using the Zymoclean Gel DNA Recovery Kit (Zymo Research). Resulting DNA samples were quantified by Qubit fluorometer (Invitrogen), normalized and pooled. Pool size was verified via Tapestation 1000 HS and was sequenced on an Illumina NextSeq 1000 P2 (2×150 nt) with 20 % PhiX.

### ACE assay analysis

In order to produce quantitative binding scores from reads, the following processing and quality control steps were performed:

1. Paired-end reads were merged using FLASH2 [37] with the maximum allowed overlap set according to the amplicon size and sequencing reads length (150 bases for all the libraries described in this manuscript).
2. If UMIs were added during amplification, the downstream UMI tag (last 8 bases) was moved to the beginning of the read, and the UMI Collapse tool [38] was used in FASTQ mode to remove any PCR duplicates. Only fully identical sequences were considered to be duplicates and error correction was not performed at this stage.
3. Primers were removed from both ends of the merged read using cutadapt tool [39], and reads were discarded where primers were not detected.
4. Reads were aggregated across all FACS sorting gates and aligned to the reference sequence (parental version of the amplicon) in amino acid space. Alignment was performed using the Needleman–Wunsch algorithm implemented in Biopython [40], with the following parameters: `PairwiseAligner, mode=global, match_score=5, mismatch_score=-4, open_gap_score=-20, extend_gap_score=-1`. Parameters were chosen by manual inspection across a number of processed libraries.
5. Reads were then discarded if (1) the mean base quality was below 20, or (2) a sequence (in DNA space) was seen in fewer than 10 reads across all gates (or in less than 10 unique molecules following UMI deduplication, when available).
6. We also flagged: (1) sequences that align to the reference with a low score (defined as less then 0.6 of the score obtained by aligning the reference to itself); (2) sequences containing stop codons outside of the region of interest and (3) sequences containing frame-shifting insertions or deletions. Flagged sequences were not included in any mutation-related statistics, but were used for count normalization for binding score calculations. FastQC [41] and MultiQC [42] were used to generate sequencing quality control metrics.
7. For each gate, the prevalence of each sequence (read or UMI counts relative to the total number of reads/UMIs from all sequences in that gate) was normalized to 1 million counts.
8. The binding score (ACE score) was assigned to each unique DNA sequence by taking a weighted average of the normalized counts across the sorting gates. For all experiments, weights were assigned linearly using an integer scale: the gate capturing the lowest fluorescence signal was assigned a weight of 1, the next lowest gate was assigned a weight of 2, etc.
9. Any detected sequence which was not present in the originally designed and synthesized library was dropped.
10. For each unique amino acid variant, ACE scores from synonymous DNA sequences were averaged.
11. ACE scores were averaged across independent FACS sorts, dropping sequences for which the standard deviation of replicate measurements was greater than 1.25. An amino acid variant was retained only if we collected at least three independent QC-passing observations between synonymous DNA variants and replicate FACS sorts.

### Surface Plasmon Resonance (SPR)

#### Antibody expression in SoluPro™ *E. coli* B strain

Individual SoluPro™ *E. coli* B strain colonies expressing antibody Fab variants were inoculated in LB media in 96-well deep blocks (Labcon) and grown at 30 °C for 24 h to create seed cultures for inducing expression. Seed cultures were then inoculated in IBM containing inducers and supplements in 96-well deep block and additionally grown at 30 °C for 24 h. Post induction samples were transferred to 96-well plates (Greiner Bio-One), pelleted and lysed in 50 µL lysis buffer (1X BugBuster protein extraction reagent containing 0.01 KU Benzonase Nuclease and 1X Protease inhibitor cocktail). Plates were incubated for 15-20 min at 30 °C then centrifuged to remove insoluble debris. After lysis samples were adjusted with 200 µL SPR running buffer (10 mM HEPES, 150 mM NaCl, 3 mM EDTA, 0.01 % w/v Tween-20, 0.5 mg/mL BSA) to a final volume of 260 µL and filtered into 96-well plates. Lysed samples were then transferred from 96-well plates to 384-well plates for high-throughput SPR using a Hamilton STAR automated liquid handler. Colonies were prepared in two sets of independent replicates prior to lysis and each replicate was measured in two separate experimental runs. In some instances, single replicates were used, as indicated.

#### SPR experiments

High-throughput SPR experiments were conducted on a microfluidic Carterra LSA SPR instrument using SPR running buffer (10 mM HEPES, 150 mM NaCl, 3 mM EDTA, 0.01 % w/v Tween-20, 0.5 mg/mL BSA) and SPR wash buffer (10 mM HEPES, 150 mM NaCl, 3 mM EDTA, 0.01 % w/v Tween-20). Carterra LSA SAD200M chips were pre-functionalized with 20 µg/mL biotinylated antibody capture reagent for 600 s prior to conducting experiments. Lysed samples in 384-well blocks were immobilized onto chip surfaces for 600 s followed by a 60 s washout step for baseline stabilization. Antigen binding was conducted using the non-regeneration kinetics method with a 300 s association phase followed by a 900 s dissociation phase. For analyte injections, six leading blanks were introduced to create a consistent baseline prior to monitoring antigen binding kinetics. After the leading blanks, five concentrations of HER2 extracellular domain antigen (ACRO Biosystems, prepared in three-fold serial dilution from a starting concentration of 500 nM), were injected into the instrument and the time series response was recorded. In most experiments, measurements on individual DNA variants were repeated four times. Typically each experiment run consisted of two complete measurement cycles (ligand immobilization, leading blank injections, analyte injections, chip regeneration) which provided two duplicate measurement attempts per clone per run. In most experiments, technical replicates measured in separate runs further doubled the number of measurement attempts per clone to four.

#### Sensorgram baseline subtraction

Sensorgrams were generated from raw data using the Carterra Kinetics GUI software application provided with the Carterra LSA instrument. Sensorgram response values vs. time for 384 regions of interest (ROIs) on the Carterra chip were corrected using a double-referencing and alignment technique implemented by the Carterra manufacturer. This technique incorporates both the time-synchronous response of an interspot reference region adjacent to the ROI, as well as the non-synchronous response from a leading blank buffer injection flowing over the same ROI during an earlier experiment run cycle, to estimate and subtract a background response. Corrected sensorgrams were exported from the Kinetics software package for offline analysis.

#### Kinetic binding parameters

Kinetic binding parameters were estimated via non-linear regression using a standard 1:1 binding model which was modified by the incorporation of a vector of *t*_*c*_ parameters each unique to one analyte concentration. For a single analyte concentration, the association phase model is:

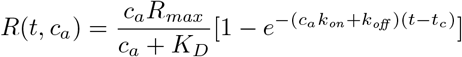

Where

*t* = time

*t*_*c*_ = concentration-dependent time offset

*c*_*a*_ = analyte concentration

**k*_*on*_* = forward (association) reaction rate constant

**k*_*off*_* = backward (dissociation) reaction rate constant

*K*^*D*^ = *k*^*off*^ /*k*_*on*_

*R*_*max*_ = asymptotic maximum instrument response.

The additional concentration-dependent time offset parameter *t*_*c*_ was needed because of the unique measurement system that Carterra uses, in which successive association phase measurements at each new analyte concentration are attempted before the analyte from the previous phase has fully dissociated, leading to response curves which do not begin from zero response at t = 0. The time offset parameters represent the projected time intercept of each association response curve; i.e., the amount of time prior to the start of the association phase, at which the measurement would have had to begin in order to reach the actual observed response at t = 0. The dissociation phase was modeled as a standard decayingexponential curve:

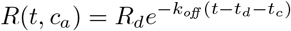

where

*t*_*d*_ = start time of dissociation phase measurement

*R*_*d*_ = final estimated response value *R*(*t*_*d*_, *c*_*a*_) from association equation.

The regression was conducted using R-language [43] scripts. Minpack.lm [44], an R-ported copy of MINPACK-1[45] [46], a FORTRAN-based software package which implements Levenberg-Marquardt [47] [48], the non-linear least squares parameter search algorithm, was used to conduct the parameter search.

#### Next-generation sequencing

To identify the DNA sequence of individual antibody variants evaluated in SPR, NGS was carried out on measured variants. Individual colonies were picked from LB agar plates containing 50 µg/mL Kanamycin (Teknova) into 96 deep well plates containing 1mL LB media (Teknova). The culture plates were grown overnight in a 30 °C shaker incubator. 200 µl of overnight culture was transferred into new 96 well plates (Labcon) and spun down at 3,500 g. A portion of the pelleted material was transferred into 96 well PCR (Thermo-Fisher) plate via pinner (Fisher Scientific) which contained reagents for performing an initial phase PCR of a two-phase PCR for addition of Illumina adapters and sequencing. Reaction volumes used were 25 µl. During the initial PCR phase partial Illumina adapters were added to CDRH2 and CDRH3 amplicon via 4 PCR cycles. The second phase PCR added the remaining portion of the Illumina sequencing adapter and the Illumina i5 and i7 sample indices. The initial PCR reaction used 0.45 µM UMI primer concentration, 12.5 µl Q5 2x master mix (NEB). Reactions were initially denatured at 98 °C for 3 min, followed by 4 cycles of 98 °C for 10 s; 59 °C for 30 s; 72 °C for 30 s; with a final extension of 72 °C for 2 min. Following the initial PCR, 0.5 µM of the secondary sample index primers were added to each reaction tube. Reactions were then denatured at 98 °C for 3 min, followed by 29 cycles of 98 °C for 10 s; 62 °C for 30 s; 72 °C for 15 s; with a final extension of 72 °C for 2 min. Reactions were then pooled into a 1.5 mL tube (Eppendorf). Pooled samples were size selected with a 1x AMPure XP (Beckman Coulter) bead procedure. Resulting DNA samples were quantified by Qubit fluorometer. Pool size was verified via Tapestation 1000 HS and was sequenced on an Illumina MiSeq Micro (2×150 nt) with 20 % PhiX.

After sequencing, amplicon reads were merged corresponding to their sample indices. Merging was performed by custom Python scripts. Scripts merged R1 and R2 reads based on overlapping sequence. Instances of unique amplicon sequences within each sample were counted and tabulated. Next, custom R scripts were applied to calculate sequence frequency ratios and Levenshtein distance between dominant and secondary sequences observed within samples. These calculations were used for quality filtering downstream to ensure clonal SPR measurements. The dominant sequence within each sample was then combined with companion Carterra SPR measurements.

#### QC

SPR fits were excluded if any of the following criteria was satisfied:

- less than 3 analyte concentrations providing usable fits
- handling errors as noted by operator
- non-physical fits (such as an upward-sloping dissociation-phase signal, even after sensorgram baseline subtraction)
- non-convergent fits
- a value of - log_10_ *K*_*D*_ `≤`8.5 coupled with an estimated signal-to-noise ratio, for the highest analyte concentration c_*a*_ included in the fit (typically 500 nM), of less than 10
- a value of- log_10_ *K*_*D*_ > 8.5 coupled with an estimated signal-to-noise ratio, for the highest analyte concentration included in the fit, of less than 70
- a *t*_*c*_ value, for the highest analyte concentration included in the fit, such that *t*_*c*_ < - 300 s or *t*_*c*_ > 0 s
- failed NGS
- non-clonal sequence (dominant sequence less than 100 times as abundant as secondary sequence when the Levenshtein distance between the two is greater than 2)
- sequence does not match any designed variant in the synthesized oligo pool (within a sequence identity tolerance to accommodate sequencing errors)

*K*_*D*_ and *k*_*off*_ were- log^10^ transformed, while *k*_*on*_ was log^10^ transformed. Distributions of kinetic parameters were visually inspected for absence of significant batch effects. Multiple measurements of the same antibody variant (usually (a) duplicate serial measurements of the same clone in the same SPR run; (b) technical replicates of the same clone from duplicate 384-well plates measured in separate runs; (c) two DNA variants with identical translation, when available; and (d) independent clones of a variant) were averaged in log space. Variants whose - log_10_ *K*_*D*_ measurements showed a coefficient of variation greater than 5 % upon aggregation were dropped.

### Observed Antibody Space (OAS) database processing

We downloaded the OAS database [49] of unpaired immunoglobulin chains on February 1st, 2022. From the full database, the following exclusions were applied to the raw OAS data: first, studies whose samples come from another study in the database (`Author` field Bonsignori et al., 2016, Halliley et al., 2015, Thornqvist et al., 2018); second, studies originating from immature B cells (BType field `Immature-B-Cells` and `Pre-B-Cells`) and B cell-associated cancers (`Disease` field `Light Chain Amyloidosis`, `CLL`); and finally, sequences were excluded if any of the following criteria was met:

- Sequence contains a stop codon
- Sequence is non-productive
- V and J segments are out of frame
- Framework region 2 is missing
- Framework region 3 is missing
- CDR3 is longer than 37 amino acids
- J segment sequence identity with closest germline is less than 50 %
- Sequence is missing an amino acid at the beginning or at the end of any CDR
- Conserved cysteine residue is missing
- Locus does not match chain type

From the resulting sequences, and for each of the two (heavy/light) chains, two types of subsequences were extracted: “CDR” and “near-full length (NF)”. In CDR datasets, we extracted only the CDR1, CDR2 and CDR3 segments as defined by the union of the IMGT [50] and Martin [51] labeling schemes. In NF datasets, we included IMGT positions 21 through 128 (127 for light chains and for heavy chains from rabbits and camels).

In all four datasets, duplicated sequences were removed, while tabulating the redundancy information (i.e. the number of times a specific sequence was observed in each study).

Sequences with a redundancy of one (i.e., observed only once in a single study) were dropped on the grounds of insufficient evidence of genuine biological sequence as opposed to sequencing errors.

A flow chart with the number of sequences filtered out and retained after each pre-processing step is shown in fig. S27.

### Model architecture

Protein language models have shown great promise across a variety of protein engineering tasks [17, 52–56]. Our architecture is based on the RoBERTa model [57] and its PyTorch implementation within the Hugging Face framework [58].

The model contains 16 hidden layers, with 12 attention heads per layer. The hidden layer size is 768 and the intermediate layer size is 3072. In total, the model contains 114 million parameters. In a pilot study, we tested larger and smaller models and compared their losses in both a masked language modeling task and a regression task. We noticed that smaller models underperformed whereas larger models did not provide significant performance boost, confirming that the selected model size was appropriate.

### Model training

#### Pre-training with OAS antibody sequences

All models for predicting binding affinity presented in this study were derived from RoBERTa architectures pre-trained on immunoglobulin sequences from the four datasets resulting from the OAS database processing (see Observed Antibody Space (OAS) database processing above). Thus, four models were trained with heavy or light chain, CDR or NF sequences. All training sequences contained species tokens (e.g. `h` for human, `m` for mouse, etc) for conditioning the language model [59]. In addition, input sequences to CDR models contained

CDR-delimiting tokens so that the originally discontinuous CDR segments could be concatenated into a single input sequence.

CDR models were used for all binding affinity and naturalness predictions, except for the CR9114 case study for which NF models were used due to framework mutations.

Model training was performed in a self-supervised manner [49], following a dynamic masking procedure, as described in Wolf et al. [57], whereby 15 % of the tokens in a sequence are randomly masked with a special `[MASK]` token. For masking, the `DataCollatorForLanguageModeling` class from the Hugging Face framework was used which, unlike Wolf et al. [57], simply masks all randomly selected tokens. Training was performed using the LAMB optimizer [60] with Є of 10^-6^, weight decay of 0.003 and a clamp value of 10. The maximum learning rate used was 10^-3^ with linear decay and 1000 steps of warm-up, dropout probability of 0.2, weight decay of 0.01, and a batch size of 416. The models were trained for a maximum of 10 epochs.

#### Fine-tuning with affinity data

Transfer learning was used to leverage the OAS-pre-trained model by adding a dense hidden layer with 768 nodes followed by a projection layer with the required number of outputs. All layers remained unfrozen to update all model parameters during training. Training was performed with the AdamW optimizer [61], with a learning rate of 10^-5^, a weight decay of 0.01, a dropout probability of 0.2, a linear learning rate decay with 100 warm up steps, a batch size of 64, and mean-squared error (MSE) as the loss function.

All models were trained for 25,000 steps. The number of steps, batch size, and learning rate for all runs were determined through a hyperparameter sweep using a pilot dataset. A grid search was run across three learning rates (10^-4^, 10^-5^, 10^-6^), three batch sizes (64, 128, 256), and two numbers of steps (25,000, 50,000). Each hyperparameter set was used to fine-tune the OAS pre-trained RoBERTa model using a 90:10 train:hold-out split from a pilot dataset (fig. S28A), and from a subset of 500 randomly selected sequences from the pilot dataset (fig. S28B). To minimize model training time while maintaining model performance, the final hyperparameters were 10^-5^ for learning rate, a batch size of 64, and 25,000 training steps.

#### Co-training with ACE and SPR data

We designed a model to predict both ACE- and SPR-derived binding affinities from sequences, using a weighted sum of the mean squared errors for each regression task as the loss function. Loss weights were inversely proportional to the dataset size. All models were evaluated using pooled out-of-fold predictions in a 10-fold cross-validation setting.

### Model characterization

#### Baselines

To assess the effectiveness of fine-tuning a pre-trained model, two baselines were evaluated.

First, a RoBERTa model with the same architecture as the pre-trained models was trained with affinity data starting from randomly initialized weights (no OAS pre-training).

Second, an XGBoost [62] model was implemented using a one-hot encoding of amino acids. The following XGBoost hyperparameters were selected using a grid search on a pilot dataset: `eta`=0.05, `gamma`=0, `n_estimators`=1000, `subsample`=0.6, `max_depth`=9, `min_child_weight`=1, `col_sample_by_tree`=1 (fig. S29). Default values were used for all other hyperparameters.

#### Out-of-distribution predictions of binding affinity

To evaluate the predictive power for binding affinities outside of the distribution seen in the training set, we fine-tuned a model by excluding any variant with log_10_ *K*_*D*_ higher than that of parental trastuzumab from the training set. We then tasked the model with predicting affinities of a set of sequences highly enriched in binders stronger than trastuzumab as validated by SPR.

#### Assessing the size and fidelity of training data

Models were trained using subsets of different sizes from datasets of varying fidelity. The trast-3 dataset was treated as the high-fidelity dataset. The low-fidelity dataset was generated by isolating a single DNA variant for each sequence from a single FACS sort, using the same preprocessing workflow. Each training dataset was evenly split into 1, 2, 4, 8, 16, 32, 64 and 128 subsets, respectively. Each training subset was used to both directly train a model with randomly initialized weights, and to fine-tune the OAS pre-trained model. A common hold-out dataset containing 10 % of data from the original trast-3 dataset was used to evaluate all models, regardless of data source or training set size. These sequences were removed from both datasets before constructing the training subsets.

#### Embeddings

Embeddings were generated by taking the mean pool of activations from the last hidden layer of the model, head excluded. The resulting size of the embedding of each sequence was 768. The dimensionality of embeddings was reduced with the Uniform Manifold Approximation and Projection (UMAP) algorithm as implemented in the RAPIDS library [63].

In a first investigation, we compared embeddings from four different models, resulting from presence or absence of OAS pre-training and presence or absence of binding affinity fine-tuning using the trast-2 dataset.

In a second investigation, embeddings were leveraged to cluster variants close in internal representation space. To this aim, dimensionality-reduced embeddings were filtered to retain only strong binders based on predicted ACE scores and 3D embeddings were clustered using HDBSCAN [64], with a minimum cluster size of 40 sequences. Sequence logo plots for each cluster were generated using Logomaker [65].

### Epistasis

Epistatic interactions between mutations were assessed by considering the predicted affinity scores for the double mutant, the constituent single mutants, and the parental antibody sequence. Specifically, the epistatic effect between two mutations, m_1_ and m_2_, was calculated as:

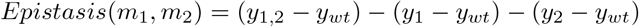

where *y*_*i*_ denotes the predicted ACE score for the mutant with mutation(s) *i*, or the parental sequence in the case of *y*_*wt*_.

### Antibody naturalness

We define the naturalness n_*s*_ of a sequence as the inverse of its pseudo-perplexity according to the definition by Salazar et al. [66] for masked language models (MLMs). Recall that, for a sequence S with N tokens, the pseudo-likelihood that a MLM with parameters ⊝ assigns to this sequence is given by:

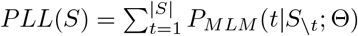

The pseudo-perplexity is obtained by first normalizing the pseudo-likelihood by the sequence length and then applying the negative exponentiation function:

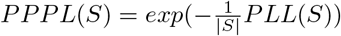

Thus, the sequence naturalness is:

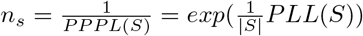

Naturalness scores were computed using the two pre-trained models described above (see Pre-training with OAS antibody sequences). Several antibody properties (immunogenicity, developability, expression level, and mutational load) were analyzed to investigate a potential relationship with sequence naturalness. For datasets in which antibodies exhibit variation in both chains (immunogenicity and expression level), the reported naturalness score was the average of the individual heavy- and light-chain scores. For datasets in which antibodies exhibit variation only in the heavy chain (developability and mutational load), only the heavy-chain naturalness score was computed. In all cases, we report naturalness scores from models trained on CDR datasets (see Pre-training with OAS antibody sequences).

To assess the relationship between naturalness and antibody properties of interest, we binned naturalness values into four equally spaced intervals (low, low-mid, mid-high, high). For each naturalness bin, we plotted the property of interest using boxplots (continuous properties) or barplots (binary properties). For boxplots, we set the whisker parameter to 1.5 and did not show outliers. The rationale for binning continuous variables was to reduce the impact of outliers and noisy data points. To assess statistical significance, we computed the Jonckheere-Terpstra test for trends.

### Immunogenicity

We obtained immunogenic responses, reported as percent of patients with anti-drug antibody (ADA) responses, from Marks et al. [29]. Of all classes of antibodies (human, humanized, chimeric, hybrid, mouse), only humanized antibodies were included in our analysis because inter-class comparisons are trivial, amounting to simple species discrimination, while intra-class comparisons are both challenging and practically relevant; (2) humanized antibodies represent the largest class (n=97) in this dataset, thereby providing the greatest statistical power; and (3) compared to the second largest class in the dataset (human antibodies) humanized antibodies have in principle more immunogenic potential due to the animal origin of their CDRs, thereby providing a practically relevant case study to assess the degree of human naturalness achieved upon engineering/humanization.

### Developability

Sequence developability was defined as a binary variable indicating whether an antibody sequence fails at least one of the developability flags computed by the Therapeutic Antibody Profiler (TAP) tool [32]. See the TAP analysis subsection for a detailed definition of these flags.

We scored hits (positive enrichment in round 3 compared to round 2, heavy-chain sequences, n=882) from the phage display library described in Liu et al. [30], which we refer to as the `Gifford Library`).

We also analyzed trastuzumab variants with up to 3 simultaneous amino-acid replacements in 10 positions of CDRH2 and 10 positions of CDRH3 (according to the same mutagenesis strategy of the trast-3 dataset).

#### Processing of the Gifford Library dataset

We obtained phage display data from the Gifford Library described in Liu et al. [30]. Specifically, we downloaded the raw FASTQ files for rounds 2 (`E1_R2`) and 3 (`E1_R3`) of enrichment from the NIH’s Sequence Read Archive (SRA) under accession number `SRP158510`.

We then followed the guidelines for processing the data as per Liu et al. [30]. First, the flanking DNA sequences of TATTATTGCGCG and TGGGGTCAA were used to pull the CDRH3 sequences. Then, sequences that included `N` or cannot be translated (divisible by 3) were excluded. Next, DNA sequences were translated into protein sequences, and were dropped if they contained a premature stop codon. Then, CDRH3 sequences shorter than 8 or longer than 20 amino acids were filtered out. Lastly, the number of occurrences of each unique sequence was determined, and sequences occurring less than 6 times were considered noise and dropped.

#### TAP analysis

We used the Therapeutic Antibody Profiler (TAP), described in Raybould et al. [32] to calculate developability scores. We used a commercially licensed virtual machine image of the tool, which was last updated on February 7^th^ 2022.

TAP calculates five developability metrics: Total CDR Length, Patches of Surface Hydrophobicity (PSH), Patches of Positive Charge (PPC), Patches of Negative Charge (PNC), and Structural Fv Charge Symmetry Parameter (SFvCSP). Furthermore, it generates flags for whether or not the metric is acceptable relative to a reference set of therapeutic antibodies. Metrics that fell outside the reference distribution are flagged as “red”, whereas metrics that fall within the most extreme 5 % of the distribution are “amber”, and metrics that fall in the main body of the distribution past the 5 % threshold are “green” and acceptable.

Using TAP, we analyzed sequence hits from the Gifford Library dataset as well as trastuzumab variants. TAP flags were used determine if an antibody had acceptable developability scores. An antibody variant was considered a failure if at least one of the TAP flags was not green.

### Antibody expression in HEK-293 cells

We collected clinical-stage antibody expression levels in HEK-293 cells from Jain et al. [31]. The dataset was heterogeneous with regard to antibody type (e.g., human, humanized, chimeric, etc). For the same reasons illustrated for immunogenicity, we focused on humanized antibodies (n=67).

In addition to HEK-293 titer, Jain et al. reported additional biophysical measurements. We did not find associations between naturalness score and biophysical parameters other than titer. However, we note that a dataset of clinical-stage antibodies is necessarily already biased towards antibodies endowed with favorable properties, meaning that distributions of biophysical parameters are strongly depleted of poorly performing antibodies. The availability of positive but not negative examples severely limits the ability to detect associations between biophysical parameters and other metrics such as naturalness.

### Mutational load

Mutational load was defined as the number of amino acid substitutions in an antibody variant compared with its parental sequence. We analyzed the distribution of naturalness scores across 6,710,400 trastuzumab variants with mutational load between 1 and 3 (10 positions in CDRH2 and 10 positions in CDRH3, allowing all natural amino acids except cysteine). We assessed the statistical significance of differences in naturalness score distributions by mutational load using the Jonckheere-Terpstra test for trends.

### Genetic algorithms

To generate sequence variants with desired properties (e.g., high/low/target ACE score and high naturalness), we developed a genetic algorithm (GA) using a tailored version of the DEAP library in Python [67]. In this GA, each individual sequence variant was reduced to its CDR representation described above (union of IMGT and Martin definitions). Each GA run was initialized from a single trastuzumab sequence. The predicted ACE and naturalness scores of each sequence were evaluated using the models described above. A cyclical select-reproduce-mutate-cull process was applied to the starting sequence pool that is common in µ +). GAs [68].

Each offspring pool contained the original 100 parents, along with 200 new, unique individuals. Of the offspring, 30 % were created from a single point mutation of a parent (excluding cysteine), and 70 % were created from two-point crossovers between two parents. Since the GA is initialized from a single sequence, the first offspring pool contained 299 individuals, all of which were created using single point mutations from trastuzumab. All sequences were constrained to remain within the trast-3 library computational space (up to triple mutants in 10 positions in CDRH2 and CDRH3, respectively). If a unique offspring could not be produced within these constraints, a randomly generated individual within the constraints was added to the offspring pool. Tournament selection without replacement (tournament size = 3) was performed to cull the population (size = 300) and select the individuals for the next generation (size = 100).

This process represented one “generation” of the GA, which was always run for 20 generations. To properly balance between the ACE score and naturalness objectives, the fitness objective was defined as:

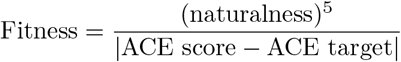

To test the generative capabilities of our models, the GA was run in the following configurations:

- Target ACE score = 9 (maximize ACE score), while maximizing naturalness
- Target ACE score = 1 (minimize ACE score), while maximizing naturalness
- Target ACE score = 6, while maximizing naturalness

Since the GA queries 300 individuals in the first generation, and 200 individuals in each subsequent generation, the GA queries 4,100 (non-unique) sequences across 20 generations. As a baseline, we randomly selected 4,100 sequences from the full mutational search space, and selected the top 100 individuals with the highest fitness as described above. The fitness function was also used to identify the top 100 individuals from the exhaustive search of the mutational space and from the trast-3 dataset.

## Acknowledgments

The authors would like to thank Joseph Sirosh, Andreas Busch, Ivana Magovcevic-Liebisch, Zach Jonasson, Kate Corcoran, Mario Sanches, Daniele Biasci, Deniz Kural, Thomas Wrona and Sarah Korman for critical review of this manuscript. Bob Albrecht, Jovan Cejovic, Joe Kaiser, Jonathan Eads, Kelechi Fletcher, Robert Pfingsten, Chris Rudnicky and Chris Vaillancourt provided engineering, MLOps and DevOps support. The authors appreciate the pilot experiments of Jerome Payet, Chang-Wook Lee and Bailey White, the technical suggestions from Jia Liu and support with schematics and formatting from Marcin Klapczynski and Stephanie Yasko.

## Competing interest statement

The authors are current or former employees, contractors or executives of Absci Corporation and may hold shares in Absci Corporation.

