## Supplementary Information and Figures for "Antibody optimization enabled by artificial intelligence predictions of binding affinity and naturalness"

#### Tite-seq CR9114 dataset

**Dataset processing** The Tite-seq CR9114 dataset [28] includes affinity data for 65,091 variants of the CR9114 bnAb heavy chain against three different influenza hemagglutinin (HA) antigen subtypes (H1, H3, and FluB). Each variant includes binary mutations in up to 16 positions based on the difference between the CR9114 germline and somatic sequences. Out of the 65,091 variants, 63,419 (97 %) bind to H1, 7174 (11 %) bind to H3, and 198 (0.3 %) bind to FluB. We downloaded the dataset from `https://cdn.elifesciences.org/articles/71393/elife-71393-fig1-data1-v3.csv`. The amino acid sequence of each variant was inferred from the binary mutation information using a custom python script, using the germline (~~QVQLVQSGAEVKKPGSSVK~~VSCKASGGTFSSYAISWVRQAPGQGLEWMGGIIPIFGTANYAQKFQGRVTITADK-STSTAYMELSSLRSEDTAVYYCARHGNYYYYYGMDVWGQGTTVTVSS) and somatic (~~QVQLVQSGAEVKKPGSSVK~~VSCKASGGTSNNYAISWVRQAPGQGLEWMGGISPIFGSTAYAQKFQGRVTISADI-FSNTAYMELNSLTSEDTAVYFCARHGNYYYYSGMDVWGQGTTVTVSS) sequences (16 somatic mutations highlighted in red, and trimmed sequences struck through). The first 19 amino acids were trimmed for compatibility with the NF Heavy model (starting from the 21$^{st}$ amino acid in the IMGT numbering scheme).

**Model architecture** We used the NF heavy-chain model initialized with weights trained on the OAS dataset (PT) as well as a model initialized with random weights (NPT). To support predictions for all three antigen targets we used a sum of the mean squared errors for each regression task as the loss function for the regression only models (Reg). In addition, we trained models using a mixture-model combining classification and regression tasks in a joint model (Mix). The loss function for the mixture model was defined as:

$$
\begin{aligned}
\ell(x, y) = L &= \sum_{c=1}^{C} L_c^{Reg} + w^{Cls} L_c^{Cls}, \\
\ell_c^{Reg}(x, y) = L_c^{Reg} &= \frac{1}{N} \sum_{n=1}^{N} l_{n,c}^{Reg}, \\
l_{n,c}^{Reg} &= \left[ x_{n,c}^{Reg} - \left( \sigma(x_{n,c}^{Cls}) \cdot y_{n,c} + (1 - \sigma(x_{n,c}^{Cls})) \cdot B_c \right) \right]^2, \\
\ell_c^{Cls}(x, y) = L_c^{Cls} &= \frac{1}{N} \sum_{n=1}^{N} l_{n,c}^{Cls}, \\
l_{n,c}^{Cls} &= - \left[ p_c y_{n,c}^{Cls} \cdot \log \sigma(x_{n,c}^{Cls}) + (1 - y_{n,c}^{Cls}) \cdot \log(1 - \sigma(x_{n,c}^{Cls})) \right], \\
y_{n,c}^{Cls} &= \begin{cases} 0, & \text{if } y_{n,c} \leq B_c; \\ 1, & \text{if } y_{n,c} > B_c, \end{cases}
\end{aligned}
$$

where $C$ is number of targets (3), $N$ is the number of training examples in a batch (256), $y_{n,c}$ is the measured affinity of sample $n$ to target $c$, $x_{n,c}^{Cls}$ is the predicted binding classification logits score of sample $n$ to target $c$, $x_{n,c}^{Reg}$ is the predicted affinity of sample $n$ to target $c$ given that it is binding $c$, $w^{Cls}$ is the classification loss weight (0.1 for models trained with 10 % and 0.1 % of the dataset, and 0.01 for models trained with 1 % of the dataset), $B_c$ is the lower boundary for each target as determined in the original publication

(7 for H1, and 6 for H3 and FluB), $\sigma$ is the logistic sigmoid function, and $p_c$ is the positive weight score. $p_c$ is calculated dynamically for each training set as the number of negative examples for class $c$ divided by the number of positive examples for class $c$ in the training set (it is set to 1 in cases where there are no positive examples).

**Model training** We trained four types of models (Reg-PT, Reg-NPT, Mix-PT, and Mix-NPT) using three training set sizes (10 %, 1 %, and 0.1 % of 65,091), each using 10 cross-validation folds. For the 1 % and 0.1 % experiments, we randomly selected 10 folds requiring each fold to include at least one positive and one negative example for each target in the training set. To support early-stopping and classifier calibration we allocated 10 % of each test set as a separate validation set. Transfer learning was used to leverage the OAS pre-trained model by adding a dense hidden layer with 768 nodes followed by a projection layer with the required number of outputs. All layers remained unfrozen to update all model parameters during training. Training was performed with the AdamW optimizer with a learning rate of $10^{-5}$, a weight decay of 0.01, a dropout probability of 0.2, a linear learning rate decay with 100 warm up steps, and a batch size of 256. All models were trained until the validation set loss stopped improving for 50, 250, 2500 epochs for training sizes of 10 %, 1 %, and 0.1 % respectively.

**Model evaluation** Unlike typical cross-validation experiments the training sets were smaller than the test sets and therefore each variant was present in multiple test sets. For each variant we randomly selected predictions from a single model instead of using the mean predicted value to avoid introducing an ensemble effect. During inference on the test set the predicted regression values were calculated as $\hat{y}_c^{Reg}(x) = \max\left\{\sigma(x_c^{Cls}) \cdot x_c^{Reg} + (1 - \sigma(x_c^{Cls})) \cdot B_c, B_c\right\}$. Raw classification logits were converted to probabilities and calibrated using the `CalibratedClassifierCV` class of scikit-learn using `cv="prefit"` and `method="isotonic"`. Classification metrics were calculated using scikit-learn functions `balanced_accuracy_score`, `f1_score`, `precision_score`, `recall_score`, and `average_precision_score`. Specifically, *Balanced Accuracy* is defined as $\frac{1}{2}\left(\frac{TP}{TP+FN} + \frac{TN}{TN+FP}\right)$, and *Average Precision* is defined as $\sum_n \left(R_n - R_{n-1}\right) P_n$ where $P_n$ and $R_n$ are the precision and recall at the $n^{\text{th}}$ threshold of the precision-recall curve.

### Supplementary Figures

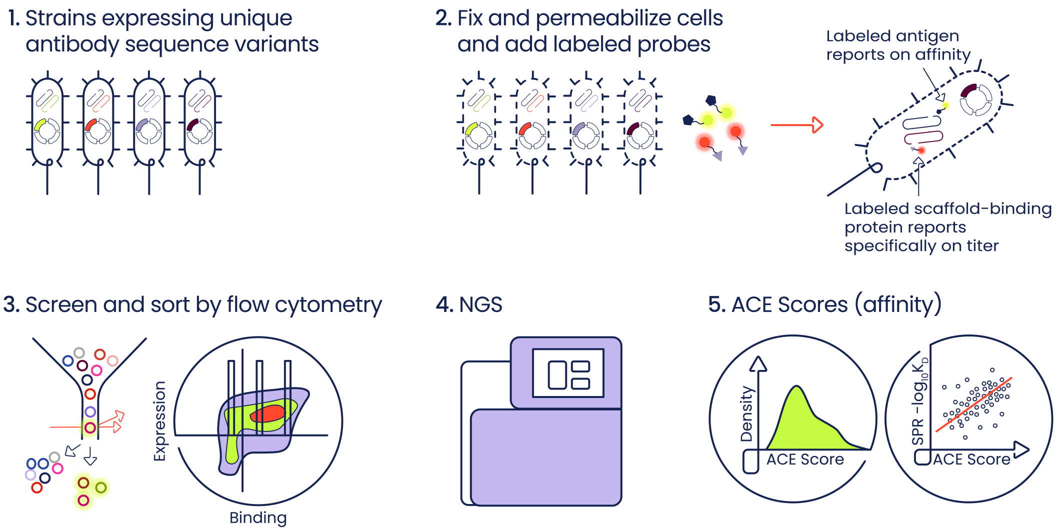


**Figure S1. Activity-specific Cell-Enrichment (ACE) assay.** A schematic representation of the ACE assay workflow. Libraries of antibody variants are expressed in SoluPro$^{TM}$ *E. coli* B Strain. Cell are fixed, permeabilized and stained with fluorescently labeled antigen and a scaffold-binding probe. Cells are then sorted based on expression and affinity levels, followed by sequencing and ACE affinity score computation.

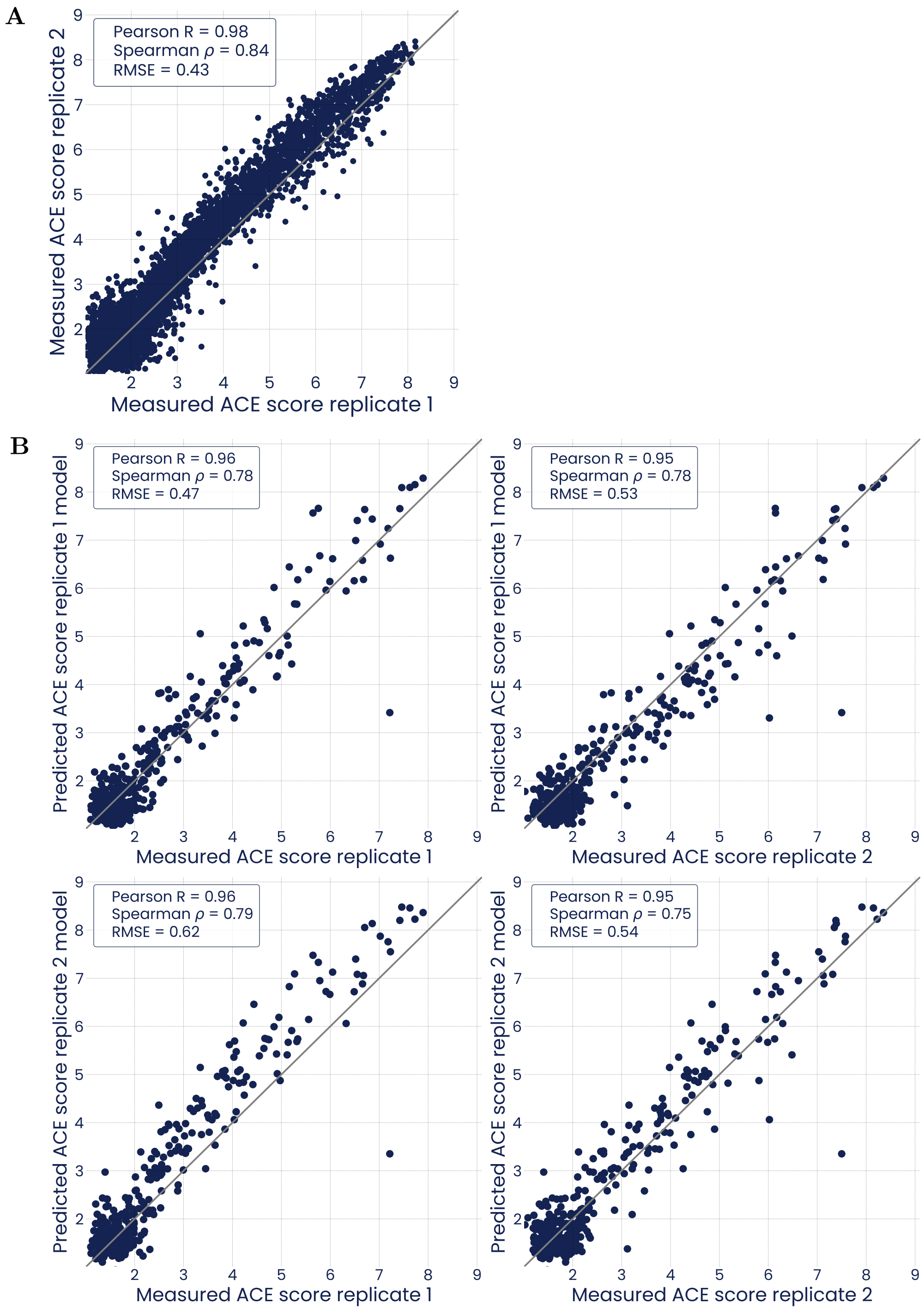


**Figure S2. Comparison of measured and predicted ACE scores between replicates in the trast-1 dataset.** Related to fig. 2C. **(A)** Comparison of ACE scores measured by two replicate FACS sorts. **(B)** All-vs-all comparison of ACE scores measured by one of two replicate FACS sorts against ACE scores predicted by models trained only with data from one of the two replicates. Error bars are 95 % confidence intervals

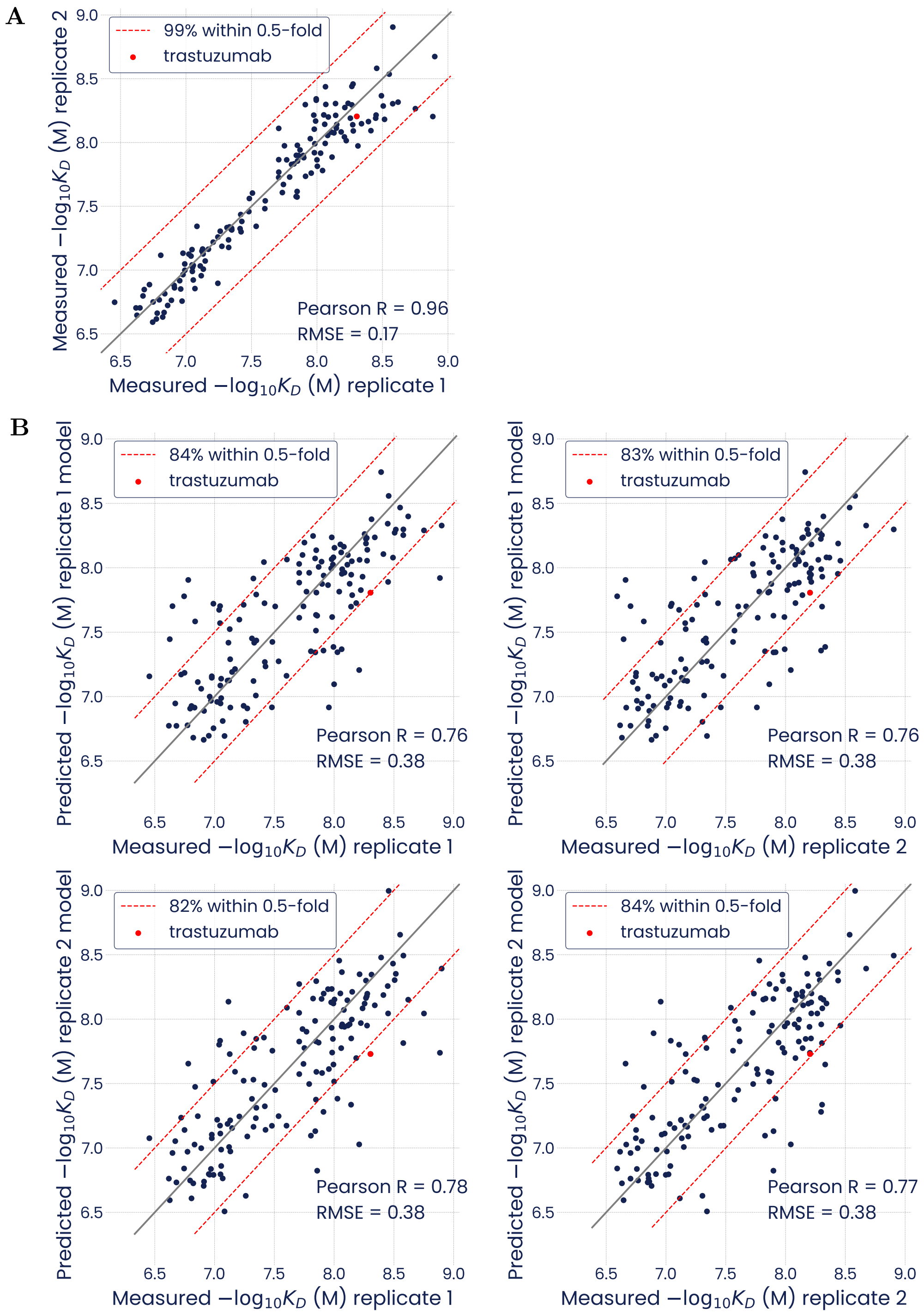


**Figure S3. Comparison of measured and predicted $-\log_{10} K_D$ values between replicates in the trast-2 SPR dataset.** Related to fig. 3B. **(A)** Comparison of $-\log_{10} K_D$ values measured by two SPR experiments. **(B)** All-vs-all comparison of $-\log_{10} K_D$ values measured by one of two replicate SPR experiments against $-\log_{10} K_D$ values predicted by models trained only with data from one of the two replicates. Error bars are 95 % confidence intervals

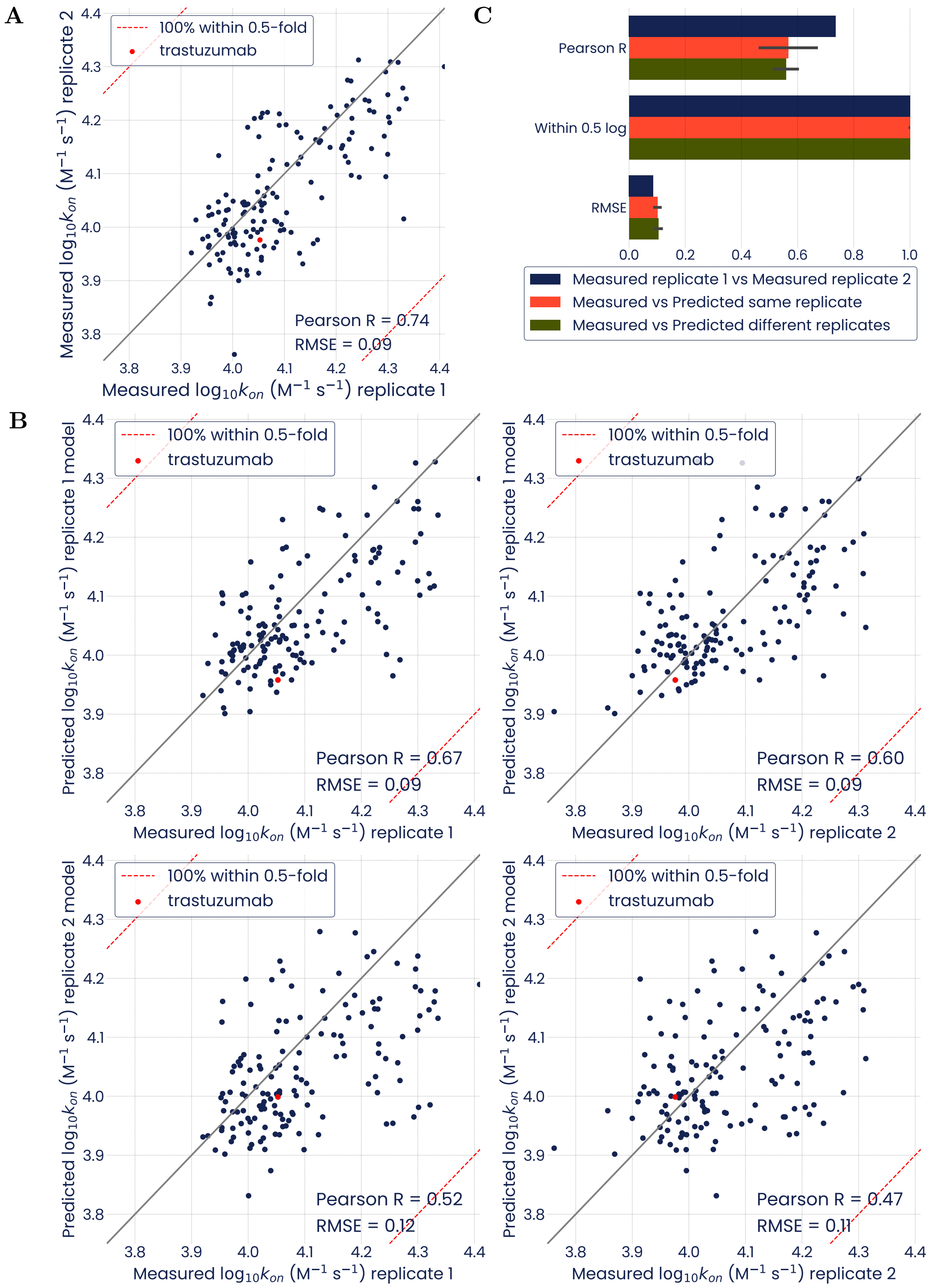


**Figure S4. Comparison of measured and predicted $\log_{10} k_{on}$ values between replicates in the trast-2 SPR dataset.** Related to fig. 3C. **(A)** Comparison of $\log_{10} k_{on}$ values measured by two SPR experiments. **(B)** Comparative analysis of replicate $\log_{10} k_{on}$ measurements and $\log_{10} k_{on}$ predicted from models trained on individual SPR replicates. **(C)** All-vs-all comparison of $\log_{10} k_{on}$ values measured by one of two replicate SPR experiments against $\log_{10} k_{on}$ values predicted by models trained only with data from one of the two replicates.

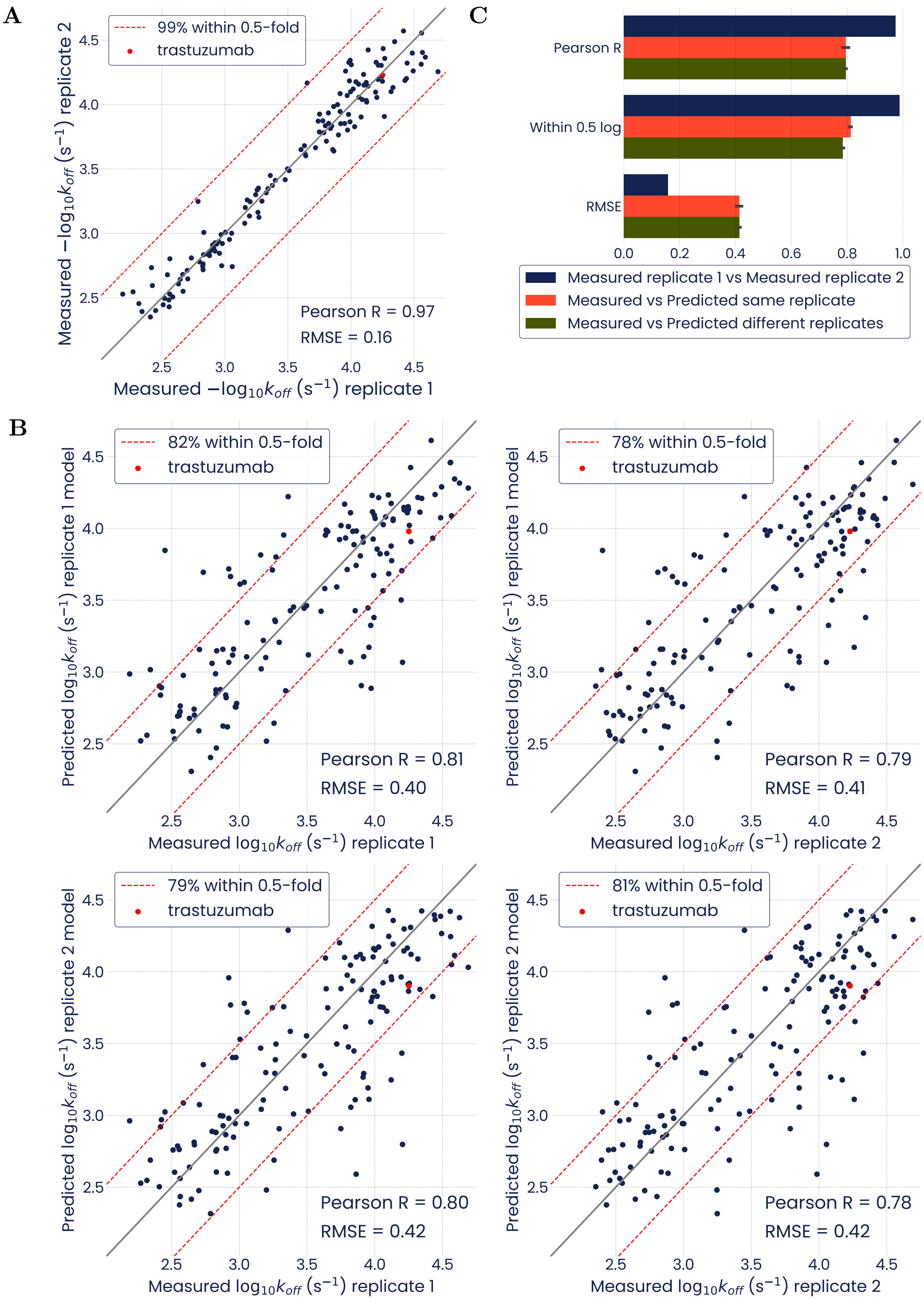


**Figure S5. Comparison of measured and predicted $-\log_{10} k_{off}$ values between replicates in the trast-2 SPR dataset.** Related to fig. 3D. **(A)** Comparison of $-\log_{10} k_{off}$ values measured by two SPR experiments. **(B)** Comparative analysis of replicate $-\log_{10} k_{off}$ measurements and $-\log_{10} k_{off}$ predicted from models trained on individual SPR replicates. **(C)** All-vs-all comparison of $-\log_{10} k_{off}$ values measured by one of two replicate SPR experiments against $-\log_{10} k_{off}$ values predicted by models trained only with data from one of the two replicates.

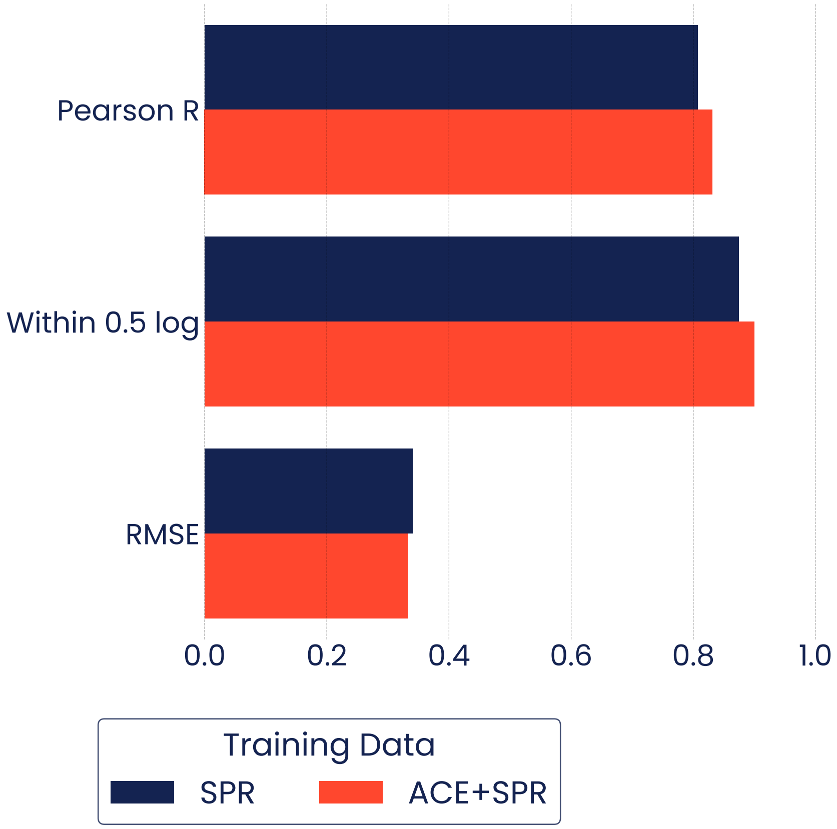


**Figure S6. Prediction of $-\log_{10} K_D$ using SPR training data alone or supplemented by ACE measurements.** Models were either trained using SPR data from trast-2, or co-trained using both ACE (trast-1) and SPR (trast-2) data. Models were evaluated using 10-fold cross-validation, predicting the $-\log_{10} K_D$ values of the sequences in each hold-out set. Pooled out-of-fold predictions were compared against SPR-measured $-\log_{10} K_D$ values.

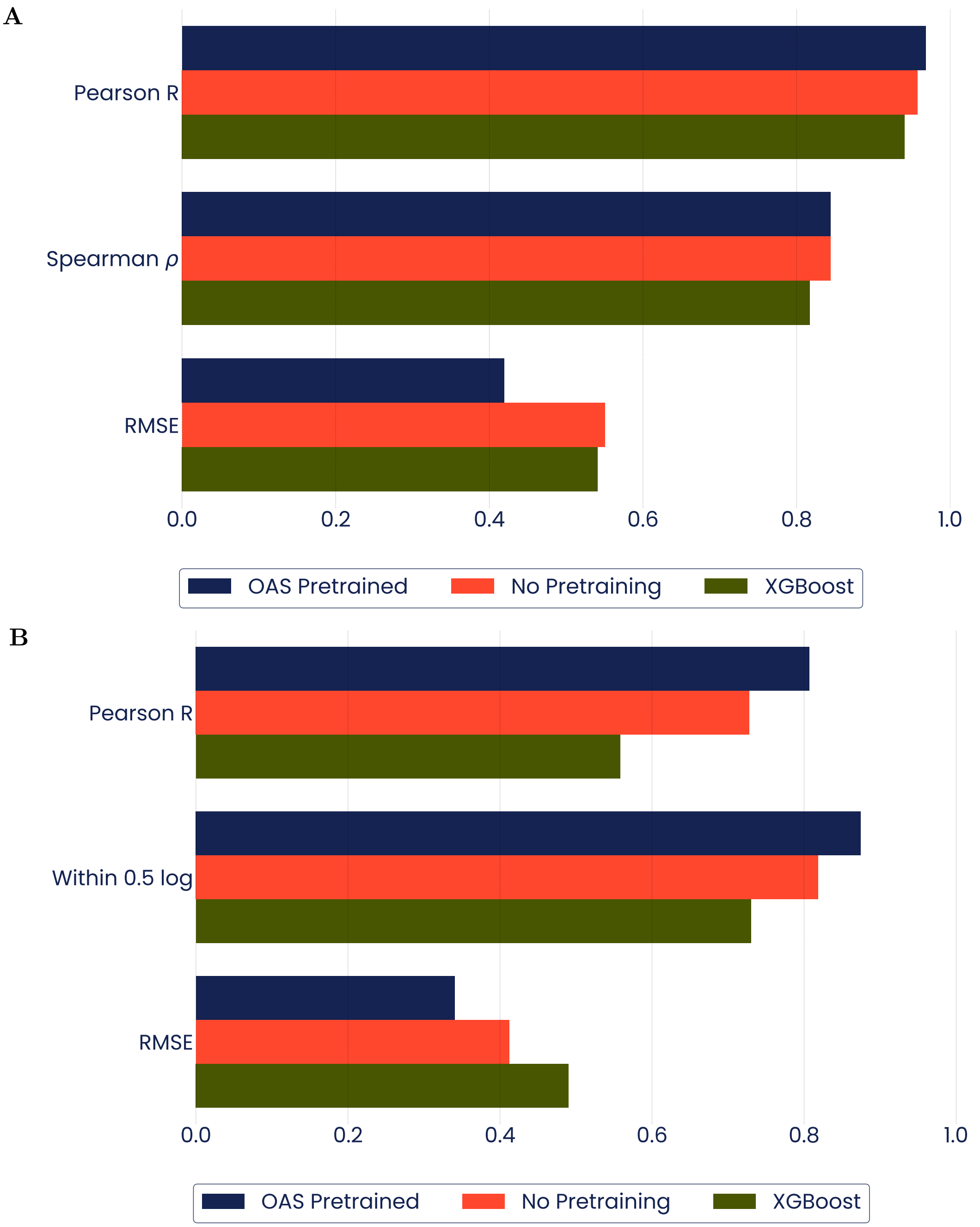


**Figure S7. Comparison of pre-trained language model performance against baselines.** The predictive performance of the OAS pre-trained deep language model was compared against two baselines: *(1)* A deep language model with identical architecture but randomly-initialized weights; and *(2)* an XGBoost model. Models were trained and evaluated using **(A)** a 90:10 train:hold-out split of ACE scores from the trast-1 dataset, or **(B)** 10-fold cross-validation with $-\log_{10} K_D$ values from the trast-2 dataset.
s

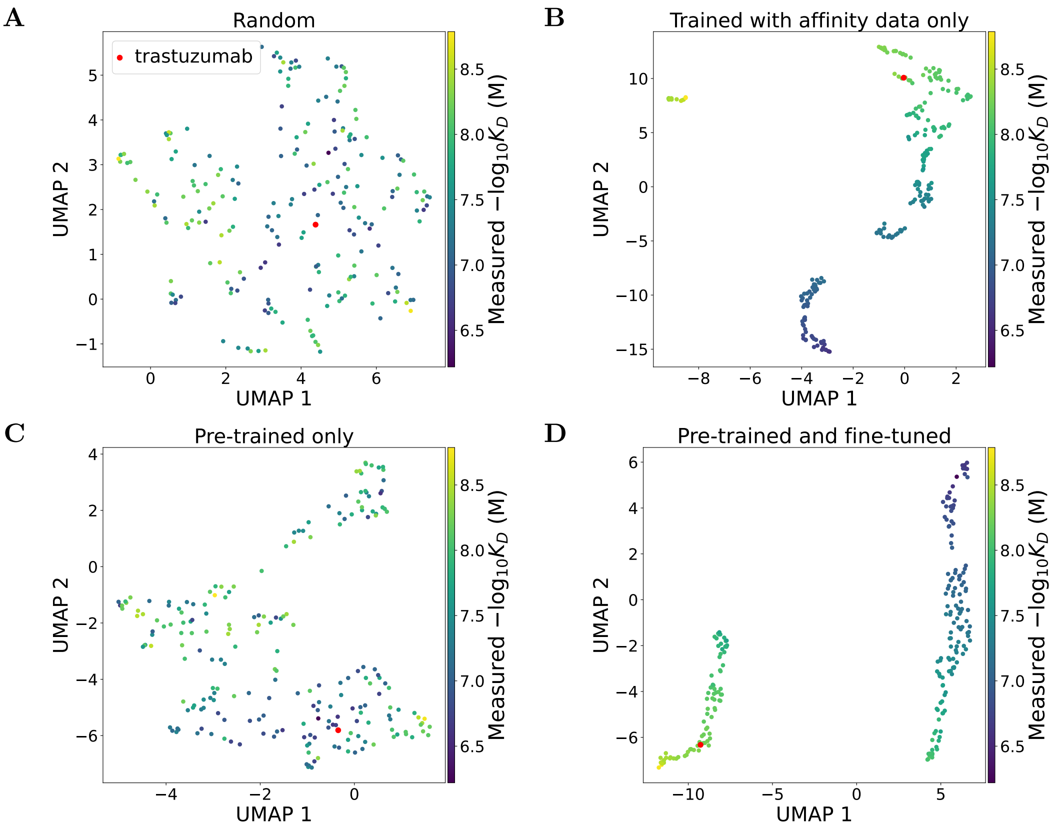


**Figure S8. Structure of model embeddings relative to binding affinities.** Pre-training with OAS-derived sequences was either **(A-B)** not performed or **(C-D)** performed. Fine-tuning with binding affinity data (trast-2 dataset) was either **(A, C)** not performed or **(B, D)** performed. In all cases, embeddings were computed with a forward pass using sequences from the trast-2 dataset, reduced to two dimensions with UMAP and color-coded by measured binding affinity.

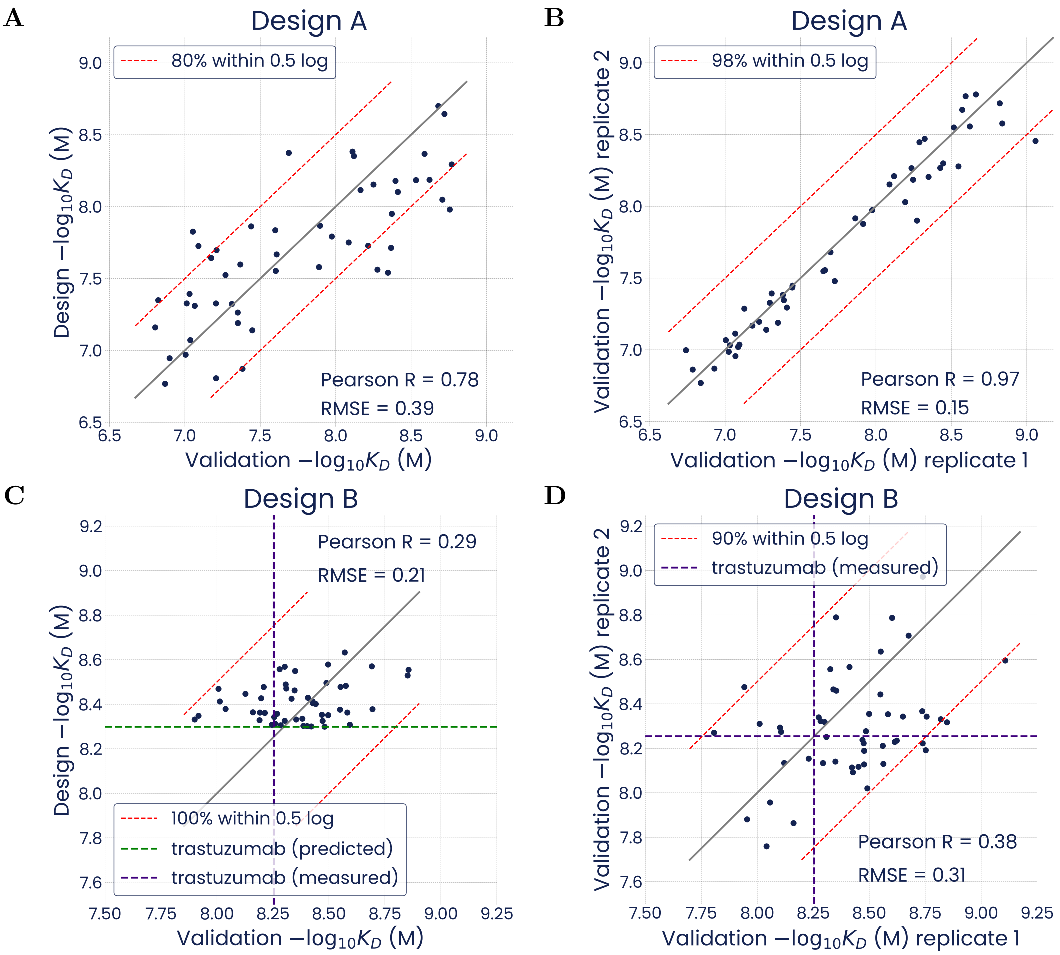


**Figure S9. Deep language models trained with the SPR-generated trast-2 dataset can design unseen sequence variants that validate in independent SPR experiments.** Related to fig. 4. **(A, C)** Scatterplots of predicted (design) and measured (validated) $-\log_{10} K_D$ values. **(B, D)** Scatterplots of measured (validated) $-\log_{10} K_D$ values in individual SPR replicates. **(A-B)** Data refers to design set A. **(C-D)** Data refers to design set B.

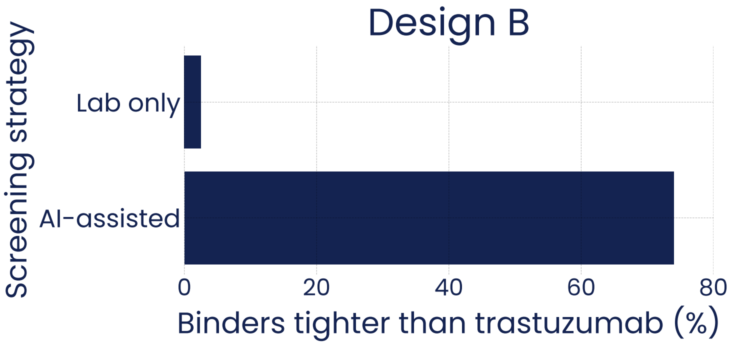


**Figure S10. Model predictions can strongly enrich for variants with desired binding properties relative to naive library screening.** Sequences of interest in design set B are defined as antibody variants binding more tightly to HER2 than parental trastuzumab (top binders). Validation rate of top binders in design set B (AI-assisted screening) as seen in fig. 4C versus prevalence of top binders in the combinatorial space (Lab-only screening) as estimated by the fraction of top binders in the model predictions for the full combinatorial space adjusted for the validation rate of design set B.

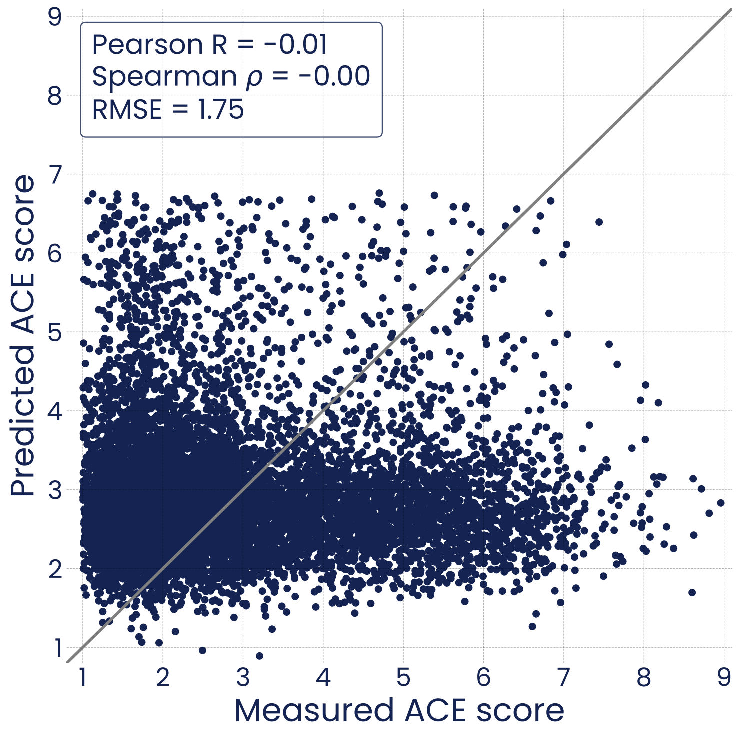


**Figure S11. Performance of a model trained on ACE scores randomized from the trast-3 datase).** The model was unable to predict ACE scores from randomly shuffled data.

**A**

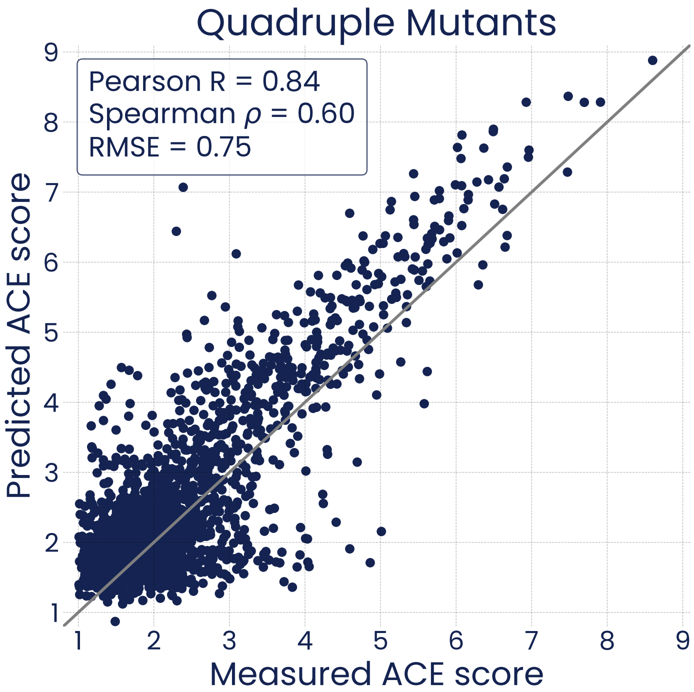


**B**

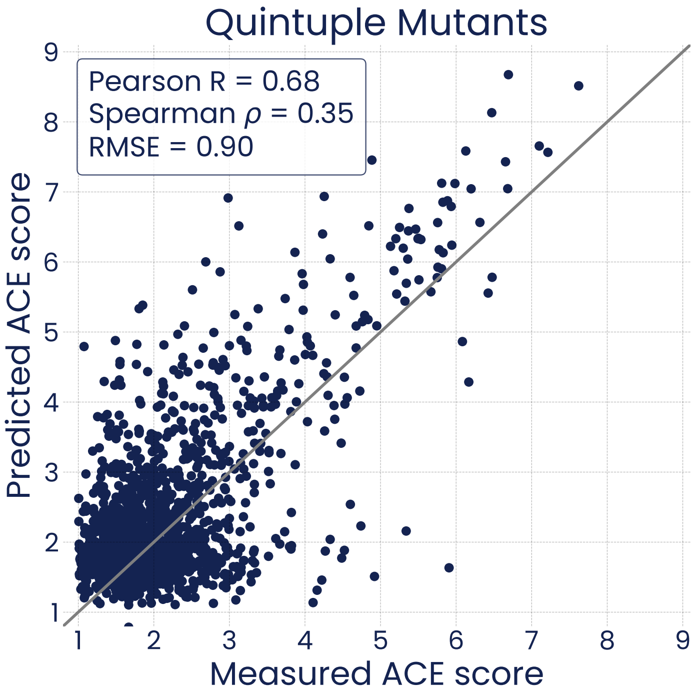


**Figure S12. Extrapolation of predictions to higher mutational loads (trast-3 dataset).** Prediction performance on **(A)** a set of quadruple mutants, and **(B)** a set of quintuple mutants. The model was trained on the trast-3 dataset of up to triple mutants of the parental trastuzumab sequence, and evaluated on a hold-out set containing only quadruple or quintuple mutants, respectively.

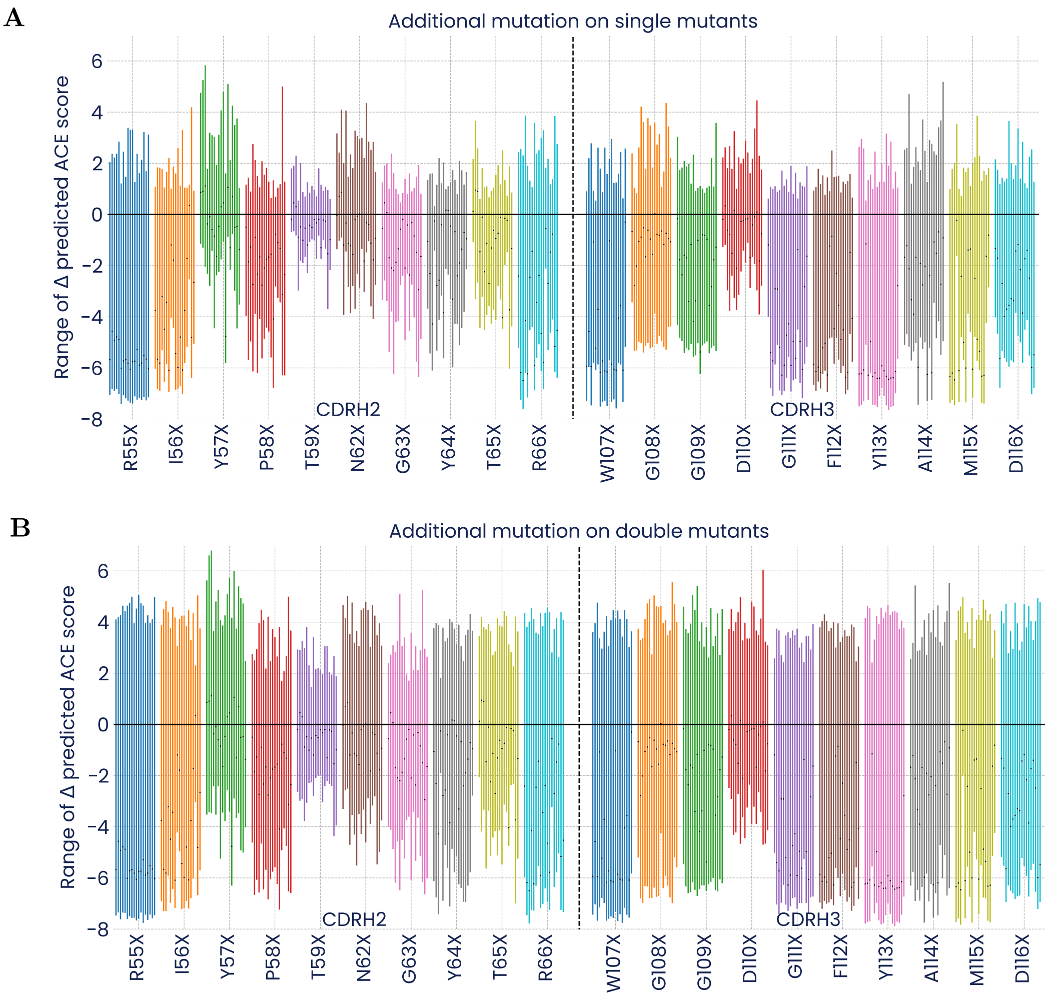


**Figure S13. The effects of individual mutations can vary strongly with the presence of other mutations.** Ranges of incremental effects (minimum to maximum) on predicted binding affinity from a model trained on the trast-3 dataset upon each individual substitution across all possible **(A)** single or **(B)** double mutants of trastuzumab. For reference, the effect of each individual single mutation on trastuzumab is indicated with black dots, identical on both panels. Mutations at each position include all possible substitutions with natural amino acids except cysteine, sorted alphabetically (i.e., $X \in [A, D, E, F, G, H, I, K, L, M, N, P, Q, R, S, T, V, W, Y]$).

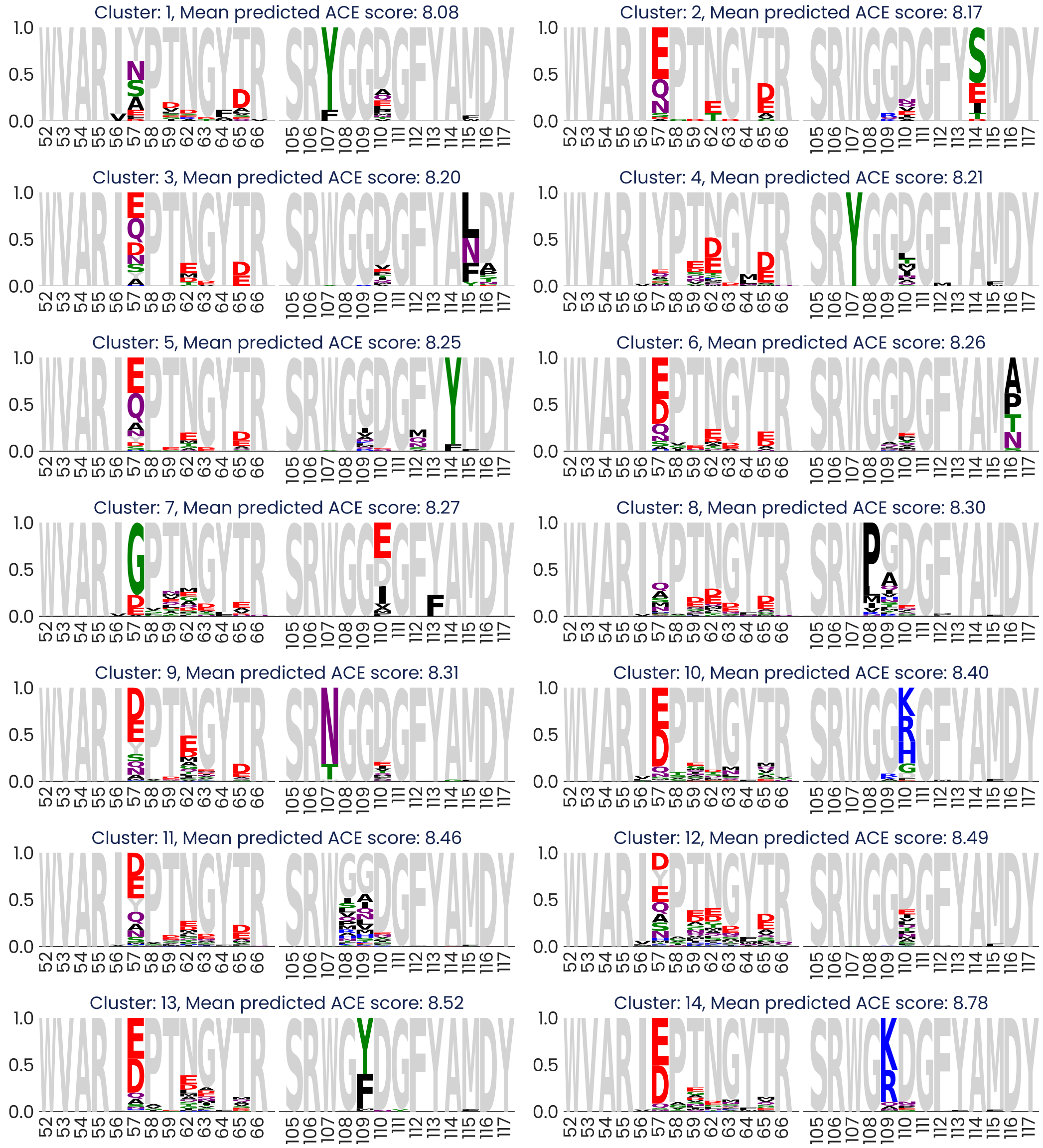


**Figure S14. AI allows identifying diverse clusters of high-affinity variants of trastuzumab.** Sequence logo plots illustrating the composition of high-affinity clusters of embeddings (predicted ACE score >8.0). Clusters were generated by reducing the dimensionality of embeddings followed by HDBSCAN clustering and are sorted by mean predicted ACE score. A minimum of 40 sequences per cluster were required. Logo plots indicate the relative frequency of each specific substitution in the sequences within each cluster. Predictions of binding affinities came from a model trained on the trast-3 dataset.

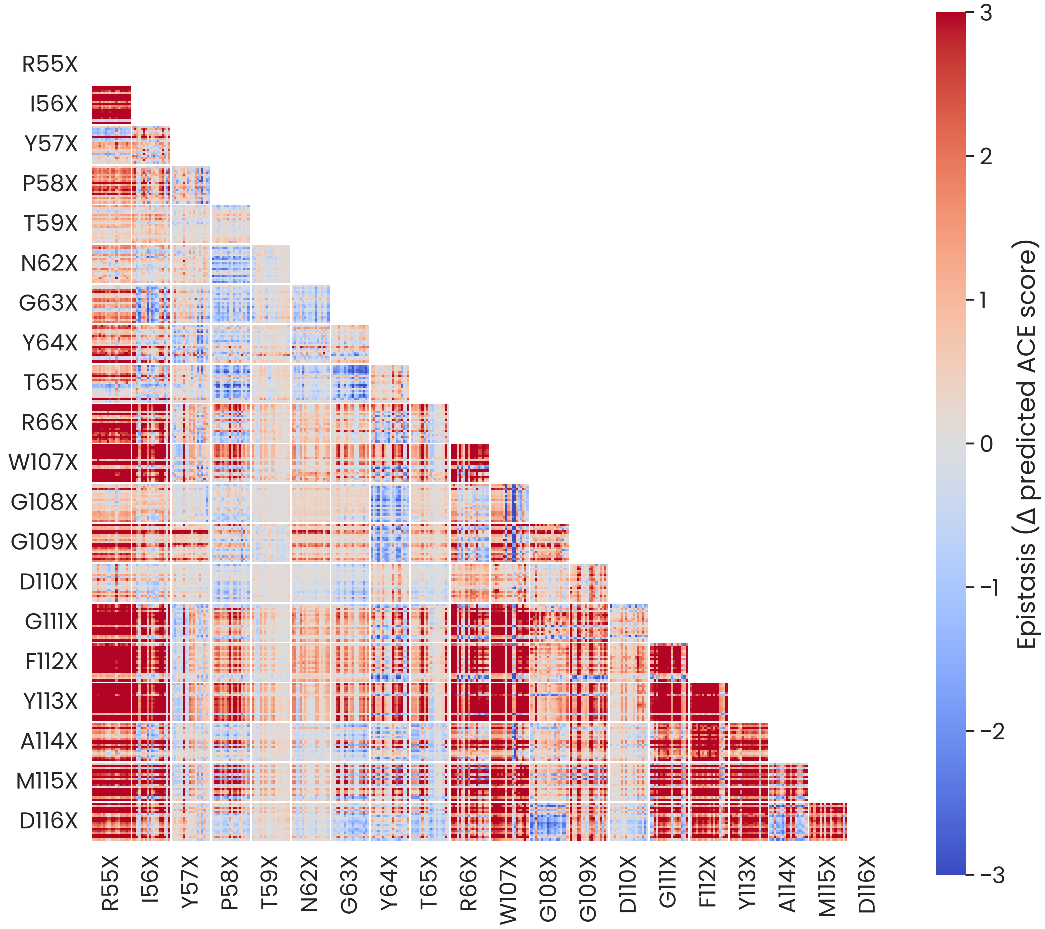


**Figure S15. Antagonistic epistasis is commonly found between key paratope residues in trastuzumab.** Heatmap illustrating epistatic effects across all possible pairs of substitutions. Epistasis refers to the deviation from additivity in the effects of two mutations when they are both present. Antagonistic epistasis refers to a smaller-than-expected change in binding affinity when two mutations co-occur. Mutations at each position include all possible substitutions with natural amino acids except cysteine, sorted alphabetically (i.e., $X \in [A, D, E, F, G, H, I, K, L, M, N, P, Q, R, S, T, V, W, Y]$). Predictions of binding affinities came from a model trained on the trast-3 dataset.

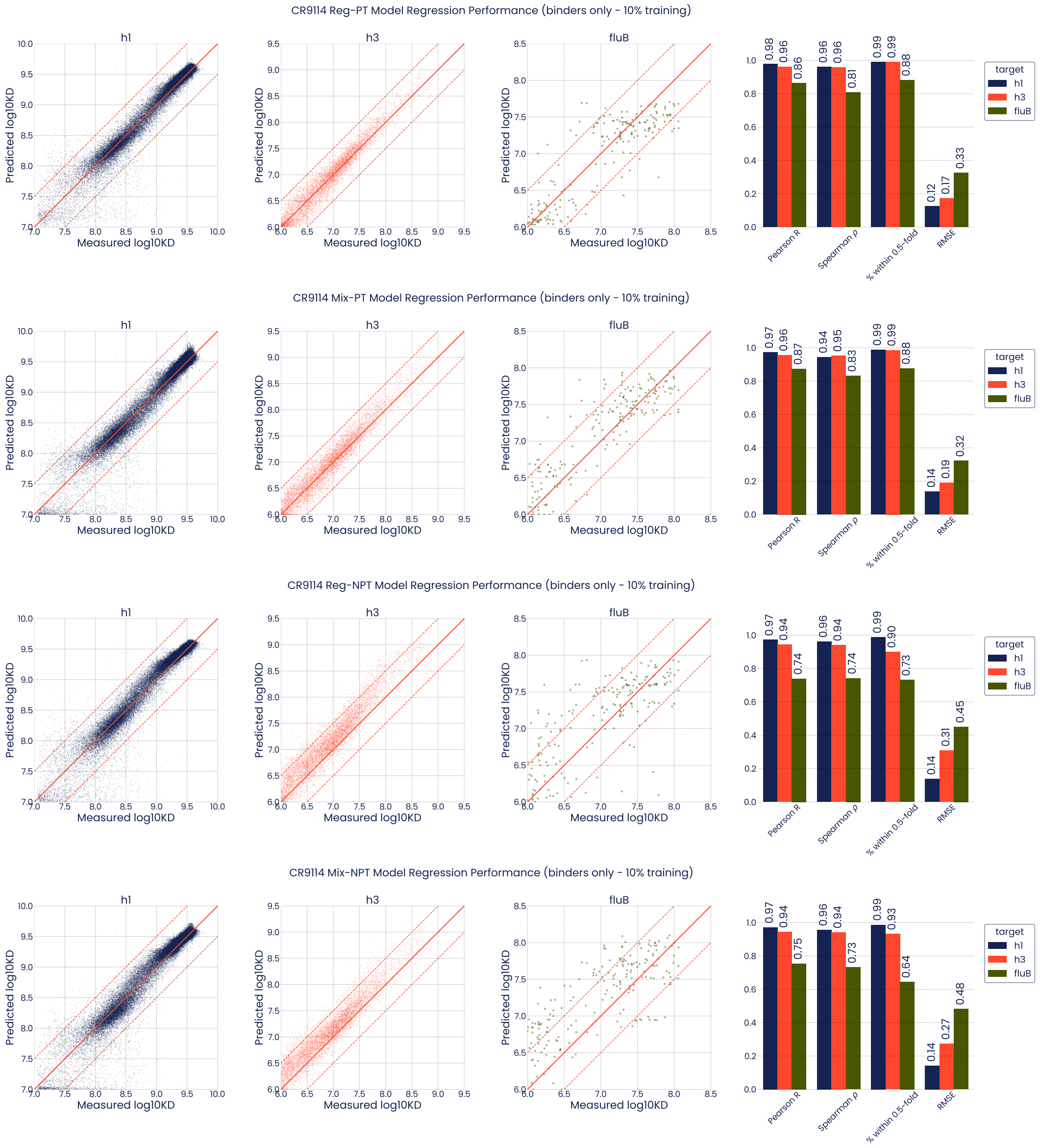


**Figure S16. Regression performance of models trained with 10 % of the CR9114 dataset**. *Reg*: Regression only model; *Mix*: Mixture classification/regression model; *PT*: initialized with pre-trained OAS-model weights; *NPT*: initialized with random weights. Results are shown for these models using pooled CV only for positive binders ($-\log_{10} K_D > B_c$, where $B_c$ is the lower boundary for each target as determined in the original publication; 7 for H1, and 6 for H3 and FluB). The full CR9114 dataset includes 63,419 (97 %) H1, 7,174 (11 %) H3, and 198 (0.3 %) FluB positive binders.

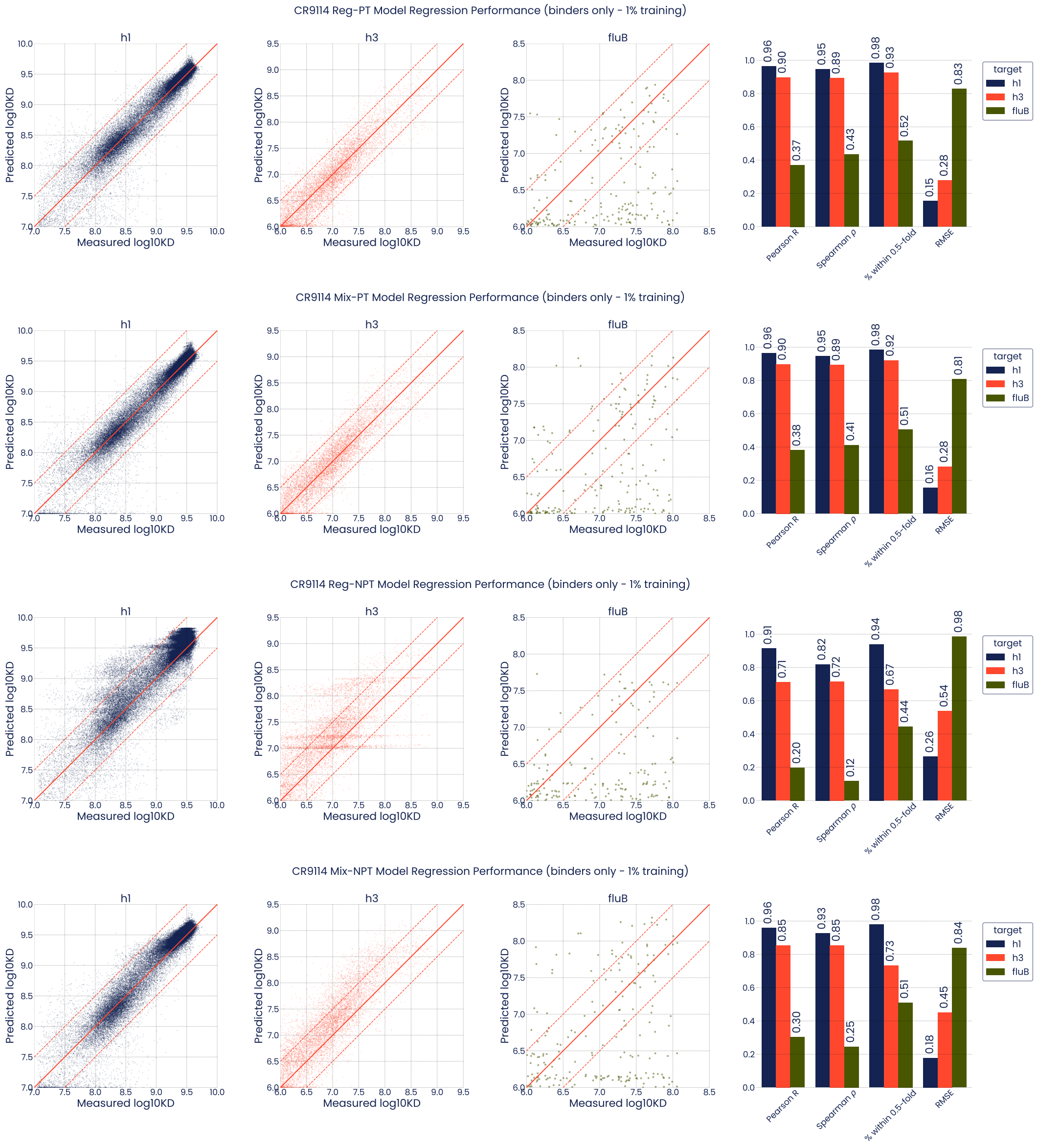


**Figure S17. Regression performance of models trained with 1 % of the CR9114 dataset**. *Reg*: Regression only model; *Mix*: Mixture classification/regression model; *PT*: initialized with pre-trained OAS-model weights; *NPT*: initialized with random weights. Results are shown for these models using pooled CV only for positive binders ($-\log_{10} K_D > B_c$, where $B_c$ is the lower boundary for each target as determined in the original publication; 7 for H1, and 6 for H3 and FluB). The full CR9114 dataset includes 63,419 (97 %) H1, 7,174 (11 %) H3, and 198 (0.3 %) FluB positive binders.

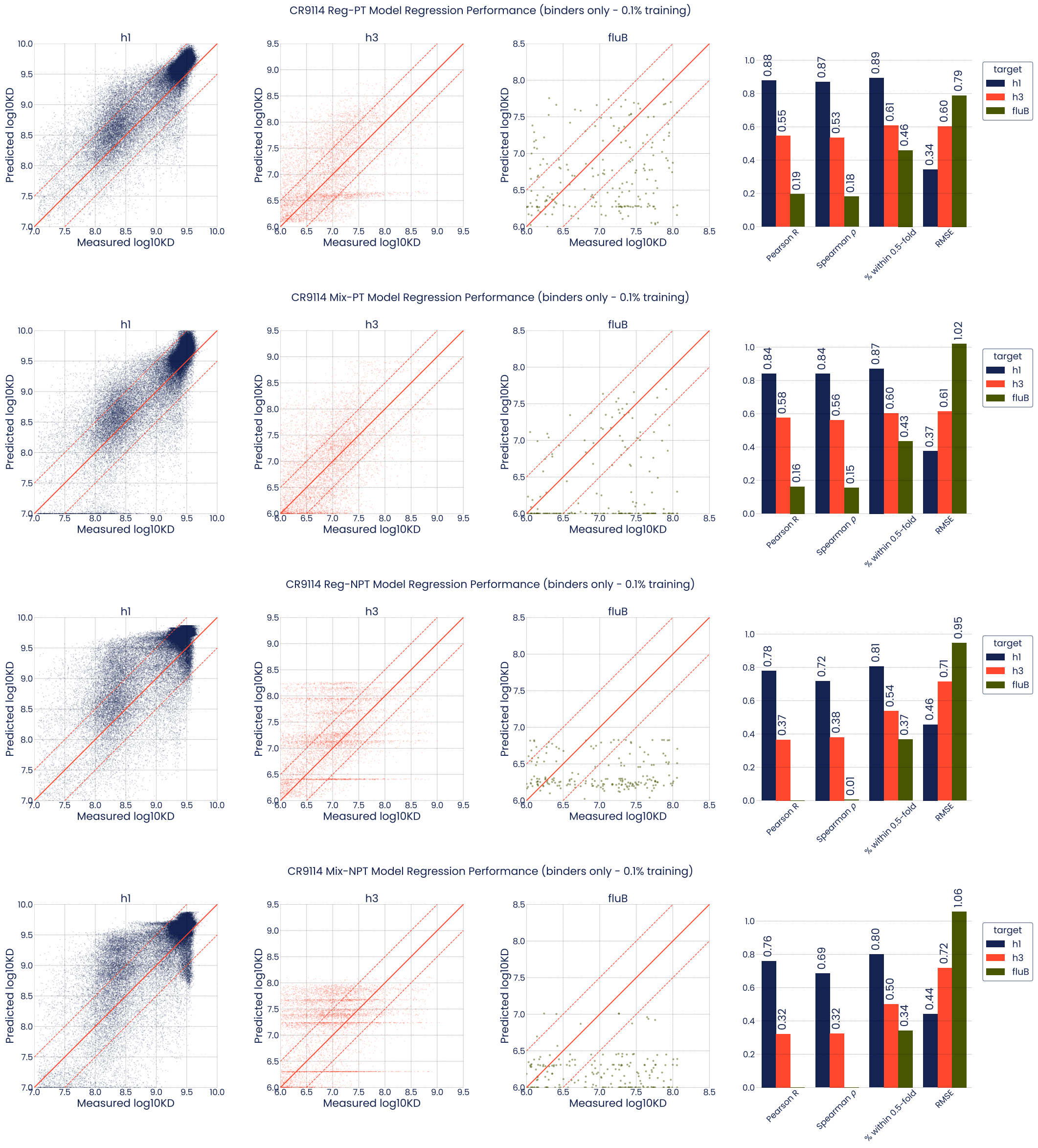


**Figure S18. Regression performance of models trained with 0.1 % of the CR9114 dataset**. *Reg*: Regression only model; *Mix*: Mixture classification/regression model; *PT*: initialized with pre-trained OAS-model weights; *NPT*: initialized with random weights. Results are shown for these models using pooled CV only for positive binders ($-\log_{10} K_D > B_c$, where $B_c$ is the lower boundary for each target as determined in the original publication; 7 for H1, and 6 for H3 and FluB). The full CR9114 dataset includes 63,419 (97 %) H1, 7,174 (11 %) H3, and 198 (0.3 %) FluB positive binders.

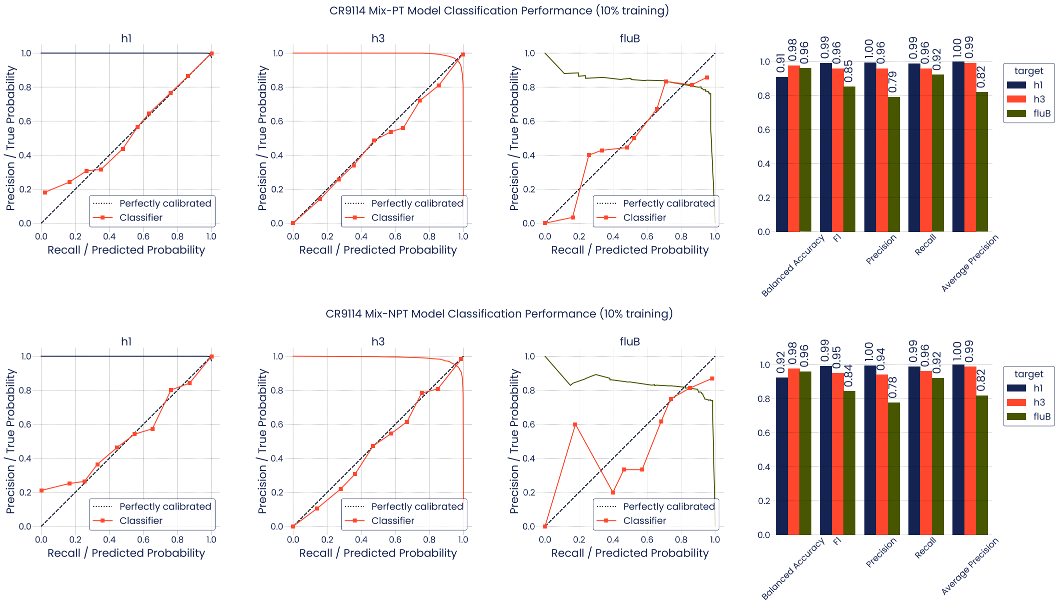


**Figure S19. Classification performance of mixture models trained with 10 % of the CR9114 dataset**. *Mix-PT*: initialized with pre-trained OAS-model weights; *Mix-NPT*: initialized with random weights. Results are shown for these models using pooled CV. For each model and target a precision-recall curve is plotted as well as a calibration curve (true probability vs. predicted probability at different scoring bins).

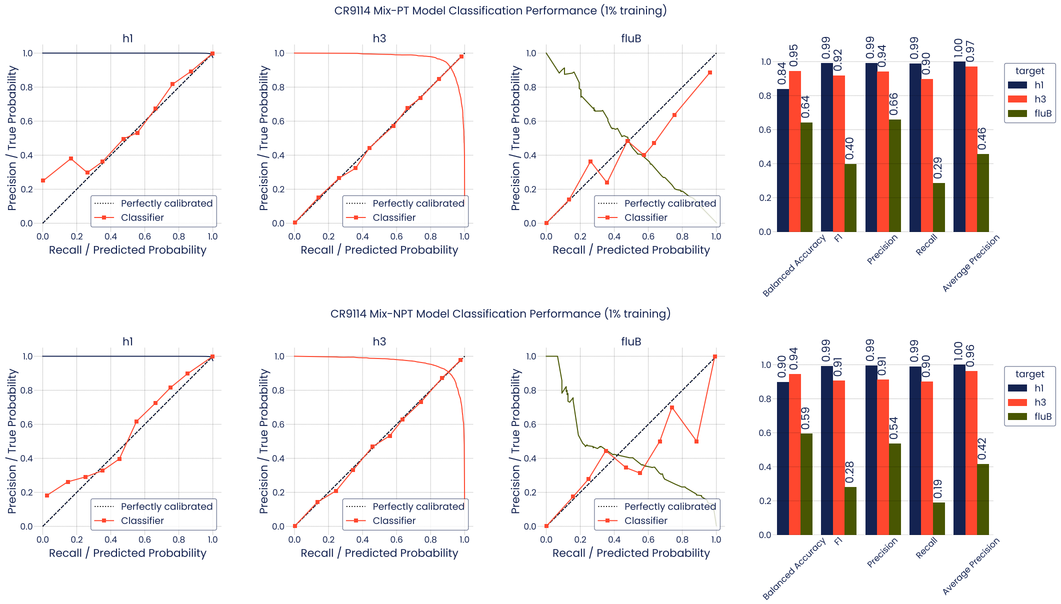


**Figure S20. Classification performance of mixture models trained with 1 % of the CR9114 dataset**. *Mix-PT*: initialized with pre-trained OAS-model weights; *Mix-NPT*: initialized with random weights. Results are shown for these models using pooled CV. For each model and target a precision-recall curve is plotted as well as a calibration curve (true probability vs. predicted probability at different scoring bins).

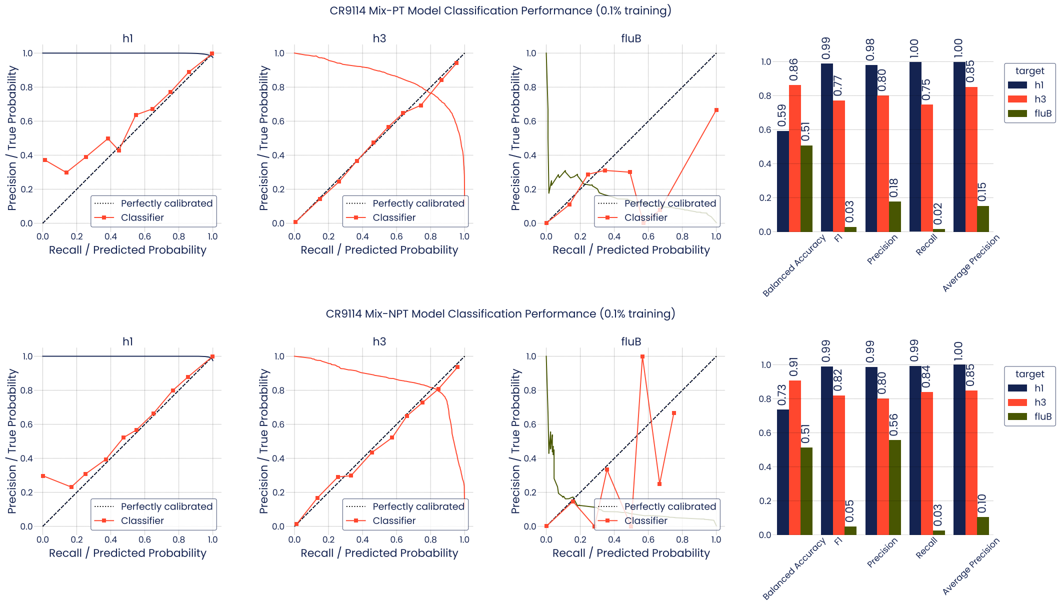


**Figure S21. Classification performance of mixture models trained with 0.1 % of the CR9114 dataset**. *Mix-PT*: initialized with pre-trained OAS-model weights; *Mix-NPT*: initialized with random weights. Results are shown for these models using pooled CV. For each model and target a precision-recall curve is plotted as well as a calibration curve (true probability vs. predicted probability at different scoring bins).

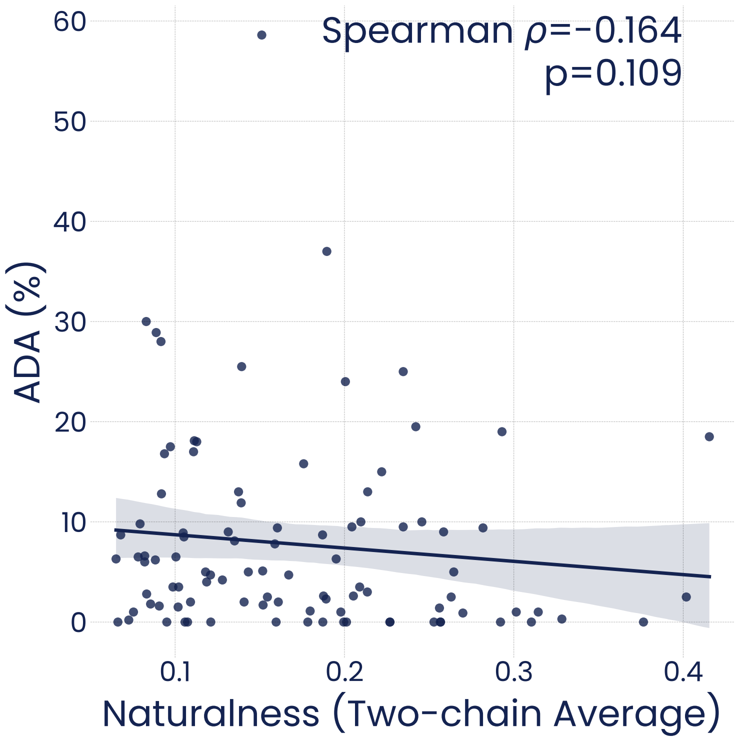


**Figure S22. Correlation between naturalness and antibody immunogenicity.** Scatter plot and Spearman correlation between model-computed naturalness scores and immunogenicity (percentage of patients with Anti-Drug Antibodies, ADA) of clinical-stage humanized antibodies from Marks et al. [29] (n=97). Shading corresponds to the 95 % confidence interval of linear regression.

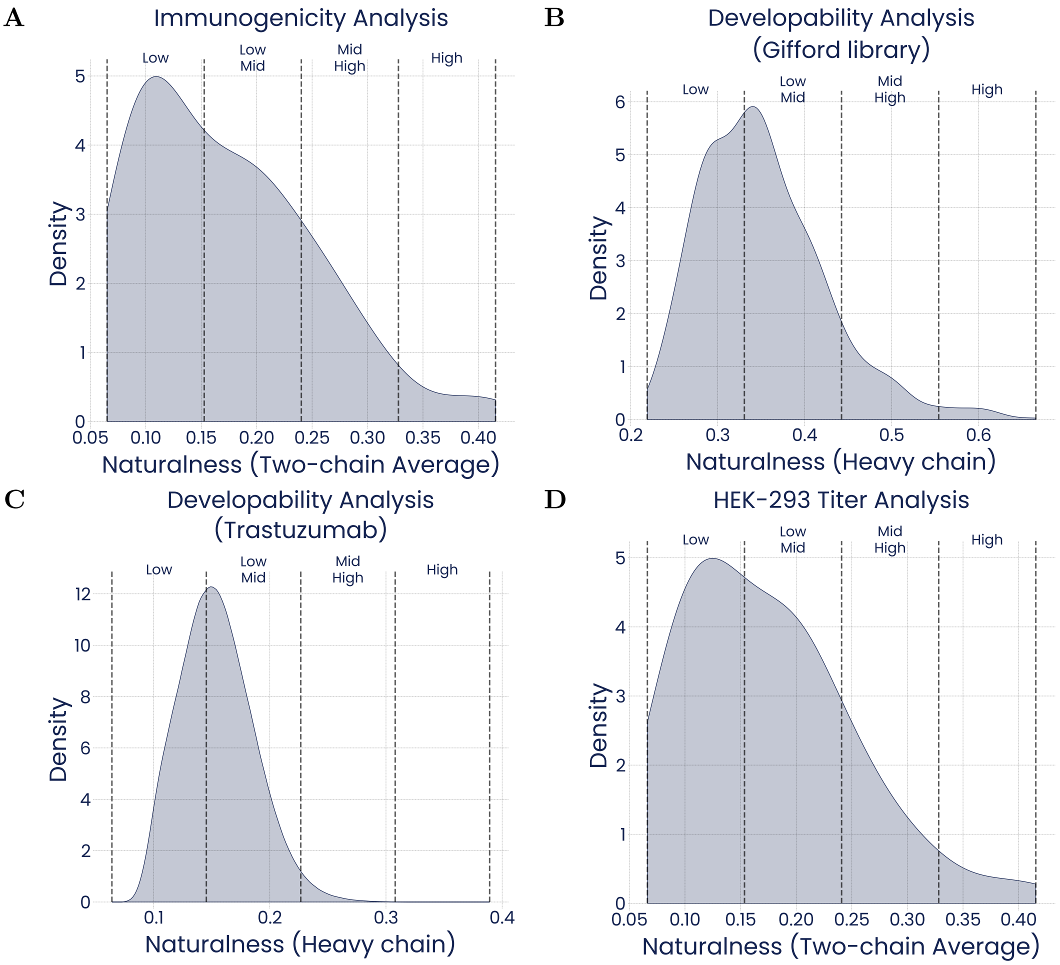


**Figure S23. Distributions of naturalness scores of antibodies from immunogenicity and developability datasets.** Related to fig. 7. **(A)** Naturalness scores of clinical-stage humanized antibodies from Marks et al. [29] (n=97) used in the immunogenicity analysis (fig. 7B). **(B)** Naturalness scores of round 3-enriched phage display hits from the Gifford library [30] (n=882) used in a developability analysis with the Therapeutic Antibody Profiler (TAP) (fig. 7C). **(C)** Naturalness scores of trastuzumab triple mutants from the combinatorial space from which the trast-3 dataset was sampled (table 1) (n=6,710,401) used in a developability analysis with TAP (fig. S24). **(D)** Naturalness scores of clinical-stage humanized antibodies from Jain et al. [31] (n=67) used in the analysis of HEK-293 expression titer (fig. 7D). Bins (Low, Low-Mid, Mid-High, High) split the naturalness range into four parts of equal size.

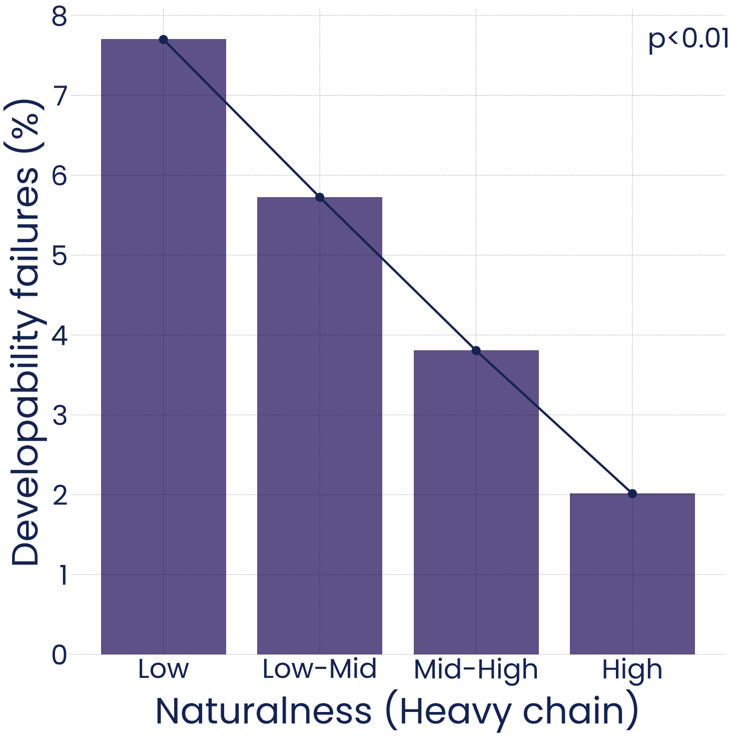


**Figure S24. Association between naturalness and developability failures of trastuzumab variants**. Model-derived naturalness scores (fig. S23C) and Therapeutic Antibody Profiler (TAP)-predicted developability failures for trastuzumab triple mutants from the combinatorial space from which the trast-3 dataset was sampled (table 1) (n=6,710,400). P-values were computed using the Jonckheere-Terpstra test for trends.

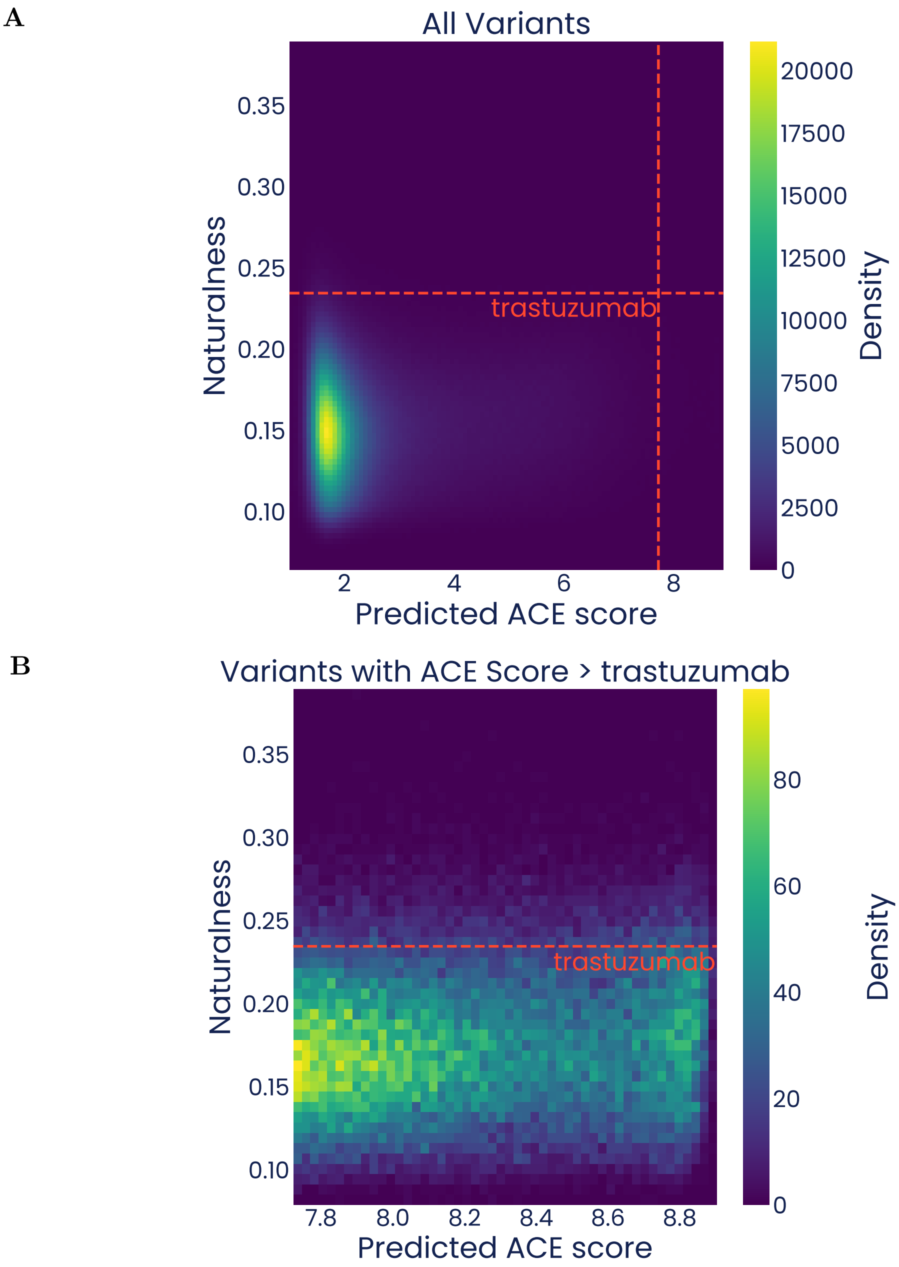


**Figure S25. Density maps of the fitness landscape for the exhaustive trast-3 search space.** Red dashed lines indicate the naturalness and/or predicted ACE score of trastuzumab. (**A**) Density map of the complete search space. (**B**) Density map of variants with predicted ACE score higher than trastuzumab.

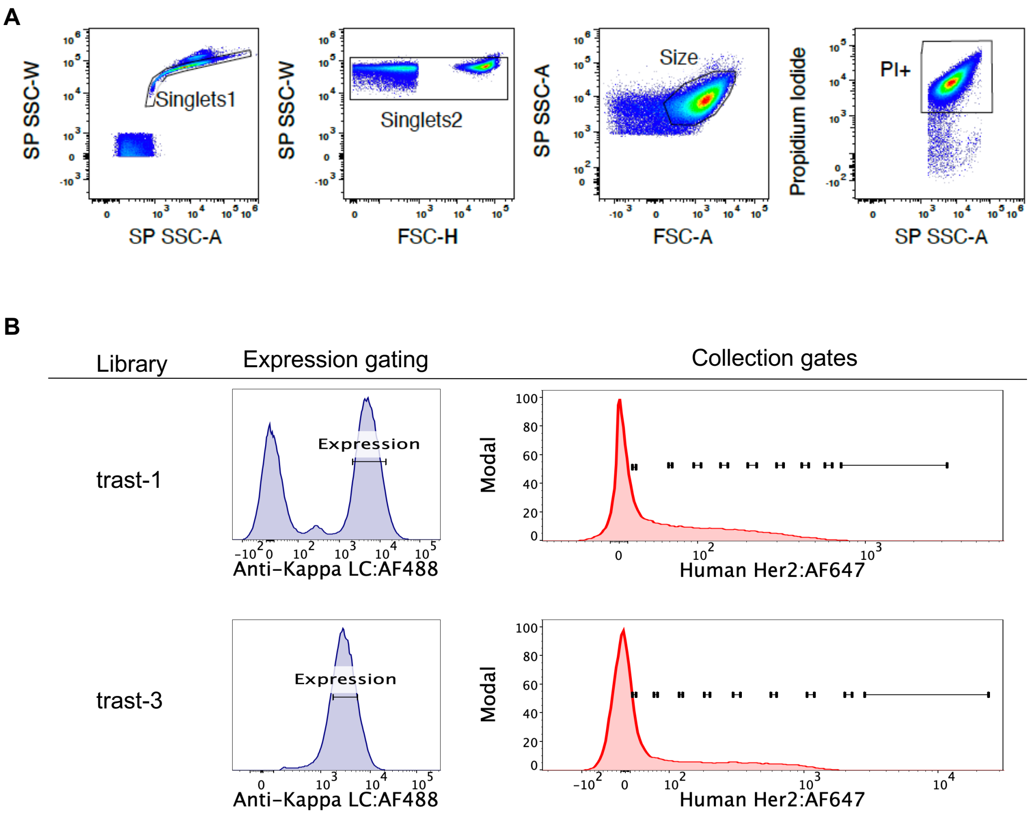


**Figure S26. Gating strategy for ACE sorting. (A)** Representative parent gating for all ACE sorts. The two singlets gates were drawn to exclude SoluPro$^{\text{TM}}$ aggregate regions previously identified by dual fluorescence of GFP and mCherry reporter strains, and propidium iodide was used to exclude unpermeabilized cells. **(B)** Specific expression and collection gating for each ACE library sort. A parent gate containing approximately 65% of the expression positive cells and centered over the peak expression signal was drawn prior to setting collection gates on the probe-specific binding signal.

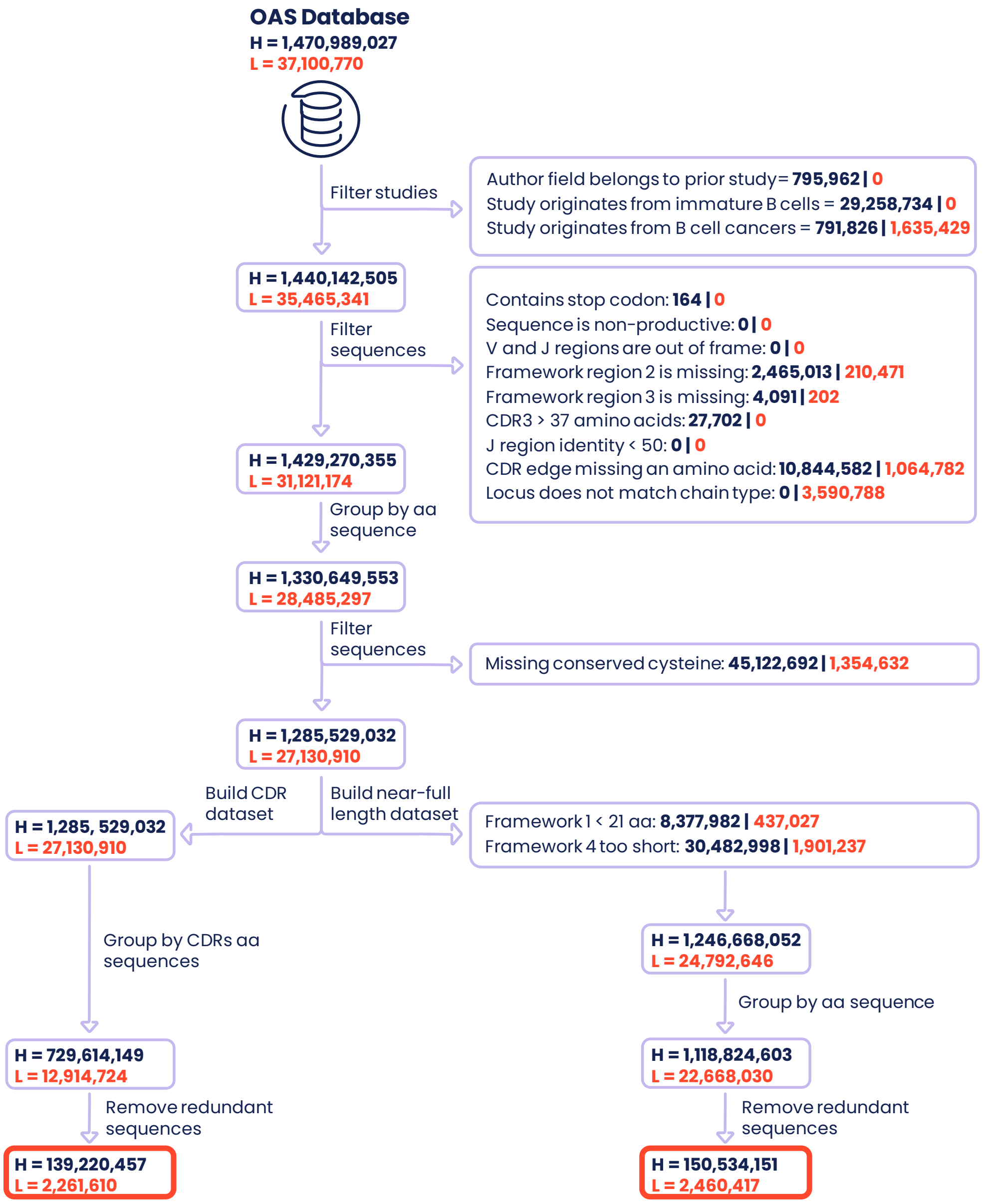


**Figure S27. OAS data preprocessing workflow.** Four datasets were built by processing the OAS dataset: for each of the two chains (heavy and light), two subsets of sequences were extracted (CDR-only and near-full length antibody sequence, see methods for details). Models trained on CDR datasets were used for all binding affinity and naturalness predictions, with the exception of the CR9114 case study for which models trained on near-full length datasets were used due to the location of mutated positions. Numbers in blue and orange correspond to the number of unique heavy (H) and light (L) chain sequences filtered out or retained at each step, respectively.

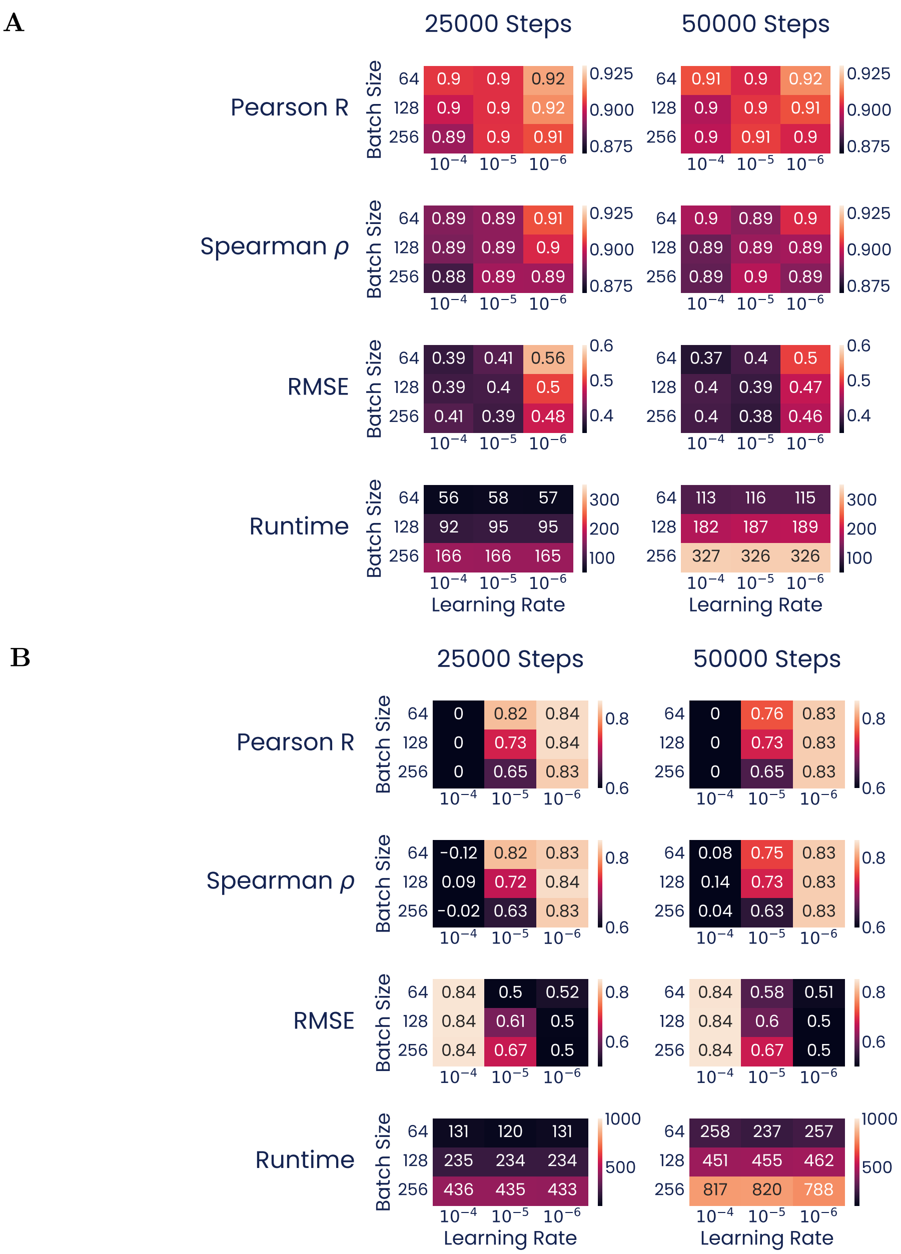


**Figure S28. Hyperparameter sweep on a pilot dataset.** A grid search across hyperparameter values was performed on **(A)** a pilot dataset, and **(B)** a subset of the pilot dataset containing 500 randomly selected sequences. Three metrics of predictive accuracy on the test set are shown for each model, along with the time required to train the model. To minimize training time while maintaining model performance, we selected the following hyperparameters: learning rate = $10^{-5}$, batch size = 64, and training steps = 25,000.

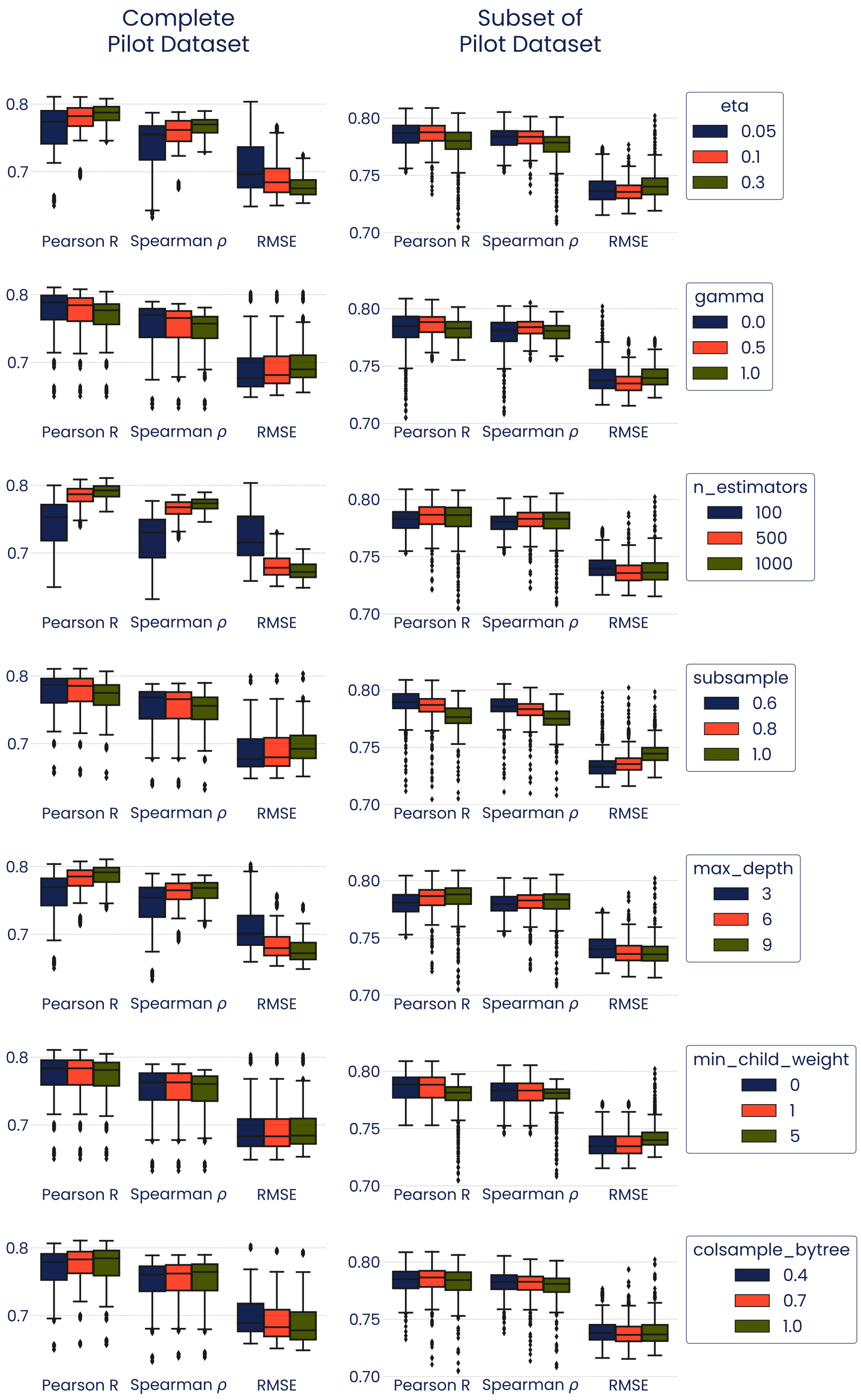


**Figure S29. Hyperparameter optimization for XGBoost baseline on a pilot dataset.** Test set accuracy of 10-fold cross-validation across each of the seven optimized XGBoost hyperparameters. An exhaustive grid search was performed across three values for each hyperparameter using both the complete pilot dataset, and a subset of the pilot dataset containing 500 randomly selected sequences.

**A**

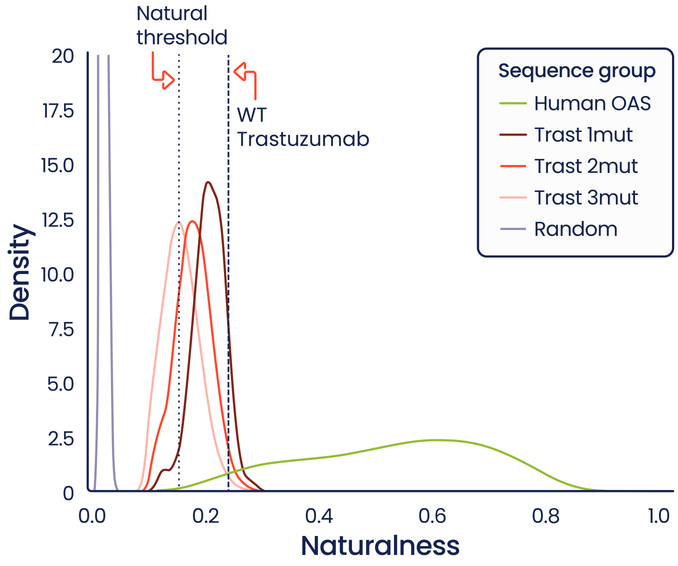


**B**

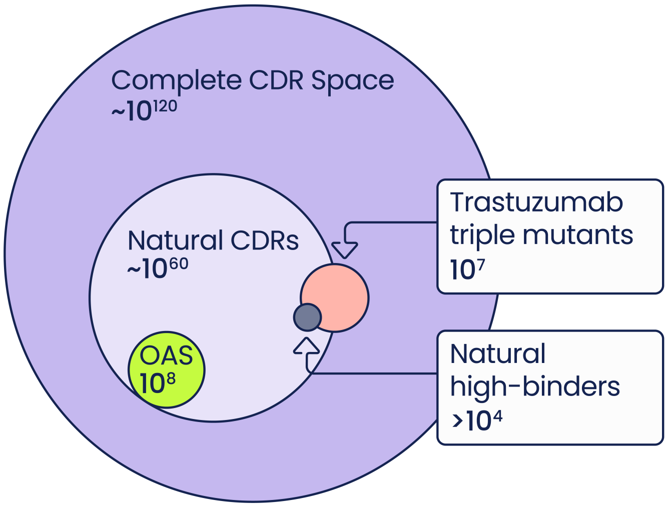


**Figure S30. Estimating sequence space sizes for heavy-chain human CDRs. (A)** A density plot of naturalness distributions for different sequence groups. For the random group, one million sequences were generated with random amino acids matching the length distribution of OAS. Based on the lower tail of OAS naturalness, a threshold of 0.15 was chosen for estimating the size of the natural sequence space. **(B)** A diagram of the relationship between sequence spaces. Circles are not to scale. The size of the total possible sequence space was estimated from 20 amino acid possibilities across 61 positions (the longest human sequence in the filtered OAS dataset). The natural CDR space was roughly approximated by fitting a skew-normal distribution to the random sequences and calculating the fraction which exceeded the naturalness threshold.
